# Optimal Precision and Accuracy in 4Pi-STORM using Dynamic Spline PSF Models

**DOI:** 10.1101/2021.10.19.464803

**Authors:** Mark Bates, Jan Keller-Findeisen, Adrian Przybylski, Andreas Hüper, Till Stephan, Peter Ilgen, Angel R. Cereceda Delgado, Elisa D’Este, Stefan Jakobs, Steffen J. Sahl, Stefan W. Hell

**Affiliations:** Department of NanoBiophotonics, Max Planck Institute for Biophysical Chemistry, 37077 Göttingen, Germany; Department of Optical Nanoscopy, Max Planck Institute for Medical Research, 69120 Heidelberg, Germany; Optical Microscopy Facility, Max Planck Institute for Medical Research, 69120 Heidelberg, Germany; Clinic of Neurology, University Medical Center Göttingen, 37075 Göttingen, Germany

## Abstract

Dual-objective 4Pi fluorescence detection enables single molecule localization microscopy, e.g. PALM and STORM, with sub-10 nanometer spatial resolution in 3D. Despite its outstanding sensitivity, wider application of this technique has been hindered by complex instrumentation requirements and the challenging nature of the data analysis. The point spread function (PSF) of the 4Pi optical system is difficult to model, leading to periodic image artifacts and compromised resolution. In this work we report the development of a 4Pi-STORM microscope which obtains improved resolution and accuracy by modeling the 4Pi PSF dynamically, while using a simpler optical design. We introduce dynamic spline PSF models, which incorporate fluctuations in the modulation phase of the experimentally determined PSF, capturing the temporal dynamics of the optical system. Our method reaches the theoretical limits for localization precision while largely eliminating phase-wrapping artifacts, by making full use of the information content of the data. With a 3D precision as high as 2 – 3 nanometers, 4Pi-STORM achieves new levels of image detail, and extends the range of biological questions that can be addressed by fluorescence nanoscopy, as we demonstrate by investigating protein and nucleic acid organization in primary neurons and mammalian mitochondria.

---

Super-resolution microscopy allows biological samples to be visualized in new ways, by bringing together the benefits of fluorescent labeling with a resolving power far beyond the classical diffraction limit^1–3^. In particular, optical designs incorporating two objective lenses which detect light coherently (4Pi detection)^4–6^ are well suited for three-dimensional (3D) visualization of transparent samples, maximizing photon collection efficiency and resolution, particularly along the optical axis. The high axial sensitivity has allowed single molecule localization microscopy (SMLM) concepts such as PALM and STORM^7–9^, when implemented with 4Pi detection, to achieve a near-isotropic spatial resolution of ∼10 nm^10–13^. Significantly, the ability to resolve 3D structure at biomolecular length scales has enabled the application of 4Pi-SMLM in studies of nanoscale protein localization^14,15^.

The advantages of 4Pi microscopy come at a cost, however. With a design similar to an interferometer, the microscope is sensitive to drift and other optical perturbations. Furthermore, the periodicity of the 4Pi point spread function (PSF) leads to the possibility of phase-wrapping errors during data analysis, which appear as periodic “ghost image” artifacts. Depending on the sample, such artifacts can make it difficult to interpret 4Pi-SMLM images, or restrict the imaging depth to a thin section of only a few hundred nanometers in the *z*-dimension. This problem can be mitigated by the addition of astigmatism to the PSF^13,16^, albeit at the expense of decreased resolution and higher complexity.

In this work we introduce a dynamic spline (DS) model of the 4Pi PSF which accurately captures the full PSF structure, while allowing temporal variations in the PSF phase to be detected and compensated. This model is an essential element of the 4Pi-STORM microscope presented here, which achieves optimal localization precision and artifact-free imaging over a large sample depth, without relying on modifications to the PSF such as astigmatism. We describe a substantially simpler and more accessible optical design which significantly improves performance. We illustrate our method with single- and multicolor nanoscale 3D imaging of nuclear pore complexes, the neuronal cytoskeleton, and mitochondria, and we highlight new analyses which are made possible by the high spatial resolution of each dataset.

## Results

### Numerical representation of the 4Pi PSF

The 4Pi-STORM setup was designed for improved stability and throughput compared to earlier implementations, for robust and routine imaging of biological samples. Details of the microscope design are given in the Methods section (see Supplementary Figs. S1-S9). The microscope employs an optical layout in which fluorescence from the sample is collected coherently by two objective lenses, and divided by polarization (S, P) and phase to be detected as four interference image channels (s_1_, p_1_, s_2_, p_2_) on an EMCCD camera (Fig. 1a). Each channel corresponds to a specific phase delay (0, π/2, π, 3π/2) between the two arms of the optical cavity. In combination, the images may be used to determine an interference phase for each single fluorophore detection event, which is equivalent to a sensitive measurement of the emitter’s *z-*coordinate (modulo the interference fringe period)^10–12^.

**Fig. 1.**
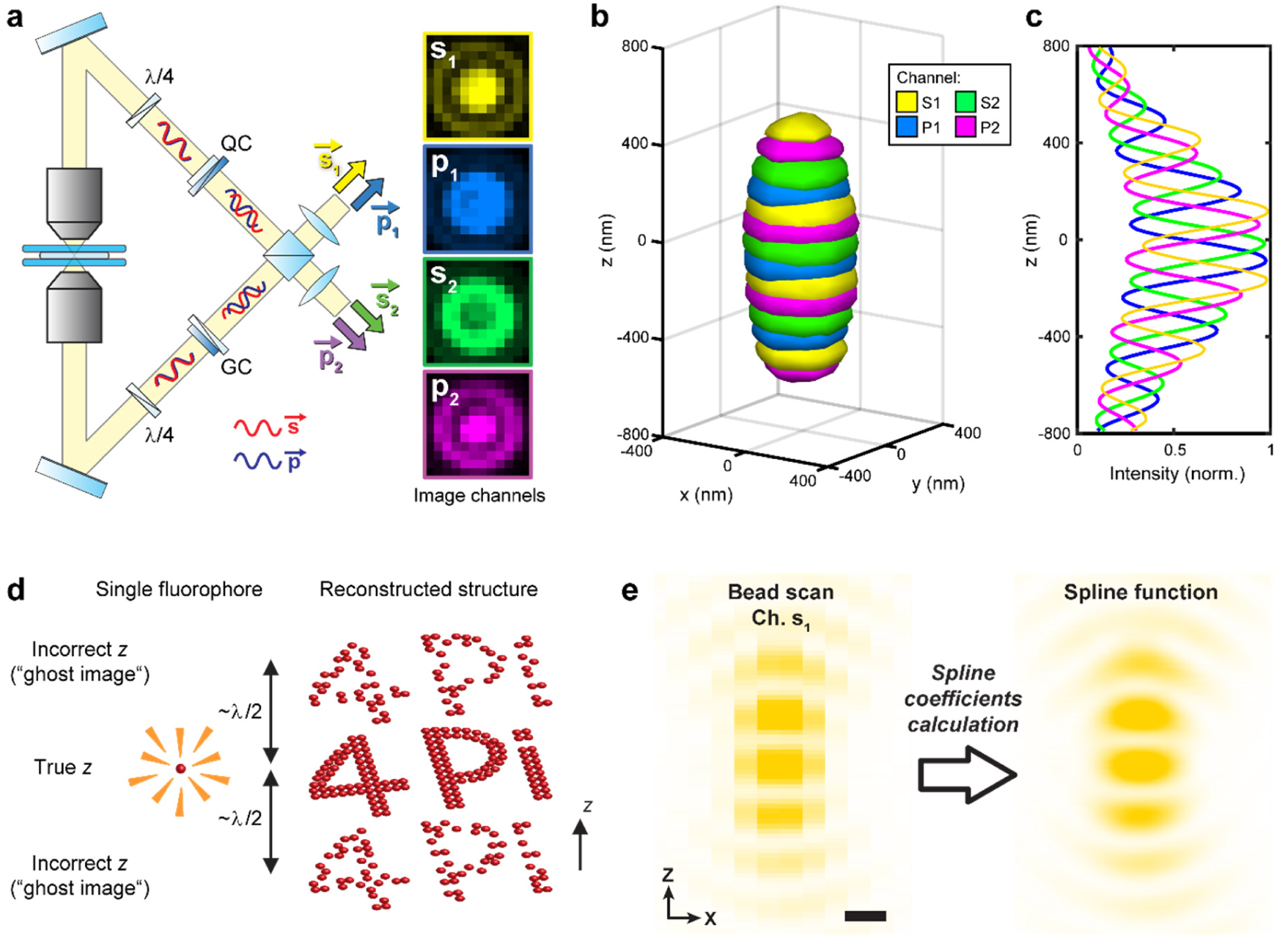
Cubic spline model of the 4Pi PSF. **(a)** Schematic diagram of the 4Pi interferometric cavity. Light from the sample is collected by two objective lenses, and the p-polarized fluorescence is delayed relative to the s-polarized fluorescence by modified Babinet-Soleil compensators (QC and GC). The photons self-interfere at the 50:50 beam splitter, resulting in four beams (s_1_, p_1_, s_2_, p_2_) each with a distinct interference phase. Images of a fluorescent emitter exhibit different patterns in each of the four channels (right panels). Quarter-wave plates, quartz wedge compensator, and glass wedge compensator are labeled by λ/4, QC, and GC respectively. **(b)** Four-channel representation of the microscope PSF, rendered as a 3D isosurface (at 35% of maximum intensity). **(c)** Central intensity profile of the PSF, measured along the *z-*axis, showing the modulation of each channel as the detected signals oscillate between constructive and destructive interference. **(d)** Schematic illustration of ghost image artifacts, arising due to localization errors which occur when an emitter’s *z-*coordinate is assigned to the wrong interference period, shifting its position by a multiple of the fringe spacing (λ/2). **(e)** A cross-section slice (*x-z*) through center of the pixelated PSF measurement (left panel), and the corresponding slice through the PSF spline function (right panel), rendered with a 20-fold higher sampling density, which captures the full detail of the PSF structure. For clarity, only channel s_1_ is shown. Scale bar: 250 nm.

The PSF of the microscope was measured by scanning a small fluorescent bead through the focal plane, which serves as a representation of a freely rotating fluorophore since the bead contains dye molecules with all orientations. The resulting intensity distribution is displayed as a 3D isosurface in Fig. 1b, with the four channels rendered in different colors (also see Supplementary Fig. S10). A central line profile through the PSF (Fig. 1c) reveals the periodic modulation of the fluorescence along the *z-*axis, as each channel oscillates between constructive and destructive interference.

This sharp periodicity is the source of the microscope’s sensitivity^10^, but it can also result in image artifacts. Estimation of the phase of the modulation is not sufficient to unambiguously determine an emitter’s *z-*coordinate, because the phase repeats for each interference fringe, with a period of approximately one half of the fluorescence wavelength (λ/2). Errors in assigning each localization to the correct fringe of the modulation pattern result in ghost images which repeat along the *z*-direction, as illustrated in Fig. 1d, obscuring the true structure of the specimen.

Recently, a new method was proposed for fluorophore localization in SMLM data analysis, in which an analytic model of the PSF, in the form of a cubic spline, is created from an experimental measurement^17,18^. We hypothesized that this approach would be particularly effective for 4Pi-STORM data analysis. As opposed to models which account for only the PSF’s phase^11^ or incorporate approximate shape measures^12,13^, a spline model is capable of fully capturing the complex structural detail of the 4Pi PSF, including asymmetric components which reduce the self-similarity in different *z-*planes.

To create an initial PSF model, we determined the piecewise-continuous 3D cubic polynomial which describes the PSF measurement. The static spline model of the PSF (*h*_4Pi_) is given by

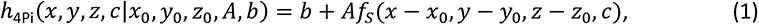

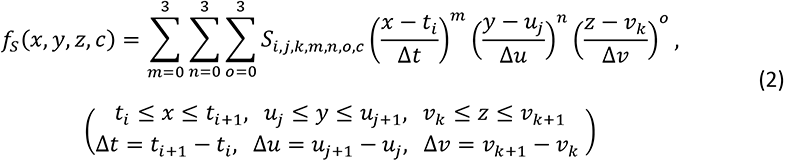

where (*x*_0_, *y*_0_, *z*_0_) is the PSF center coordinate, *A* is the amplitude, and *b* is the background offset. Here, the multichannel 3D spline function is denoted *f_s_*, where (*i*, *j*, *k*) are volume interval indices, (*t*_i_, *u*_j_, *v*_k_) are the interval center coordinates, *c* is the channel index, and *S*_*i,j,k,m,n,o,c*_ are the spline coefficients. The coefficients *S* were calculated directly from the bead scan data (see Methods). A cross-section through the center of the scan is plotted in Fig. 1e (left panel), where only one channel (s_1_) has been shown for clarity. The corresponding spline function (Fig. 1e, right panel) is smooth, reproducing the measured PSF, including the interference modulation. Conceptually, the spline model is sufficient to fully describe the detected signal from a fluorophore at any position in the sample.

The simple analytic form of the model (Eqs. 1 and 2) lends itself well to numerical optimization, but presents a computational bottleneck due to the large number of terms in the polynomial and its derivatives. To avoid this issue, we implemented multichannel 3D spline functions in Gpufit, an open-source GPU-accelerated curve fitting library which we developed previously^19^. Using this approach, we were able to rapidly estimate the spatial coordinates of single emitters by fitting the fluorophore images with the spline function derived from the PSF measurement (see Methods).

### Dynamic Spline PSF model

While the 4Pi interferometric detection concept has advantages in terms of precision, it is highly sensitive to perturbations in the optical cavity, such as thermal drift. A degree of cavity drift is unavoidable, and its effect on the PSF is to cause a shift in the phase of the interference modulation.

We tested for phase drift in our system by measuring the PSF at different time points. Two bead scans recorded 120 minutes apart showed a clear shift in the modulation phase with respect to the center of focus, as shown in Fig. 2a, despite the optical system being fully enclosed and temperature-regulated. This result suggests a potential problem for the spline-based PSF model: if the phase of the system shifts after the bead scan is recorded, the data will no longer be well-described by the model, leading to inaccurate fit results.

**Fig. 2.**
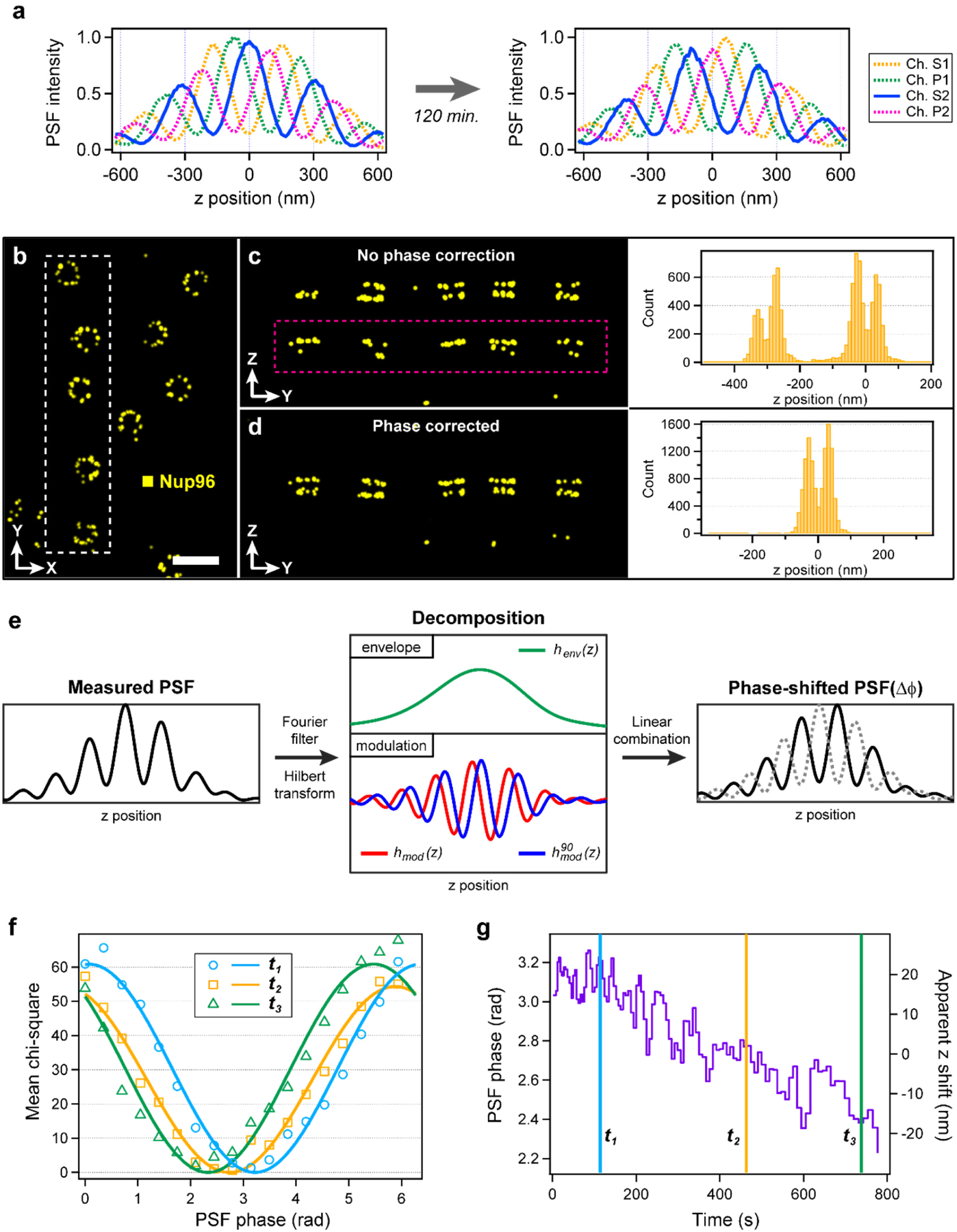
PSF phase estimation and correction. **(a)** Central profile of the 4Pi PSF at two time points during an experiment, showing a shift in the phase of the modulation relative to the focal plane (*z*=0). **(b)** 4Pi-STORM image of nuclear pore complexes, in which SNAP-tagged Nup96 is labeled with Alexa Fluor 647. Top view (*x-y* projection). Scale bar 250 nm. **(c)** Side view (y*-z* projection) of the boxed region in (b), without phase correction, exhibiting a strong ghost image artifact of the Nup96 structure (magenta box). A histogram of the *z-*coordinates of the localizations (right panel) also shows the artifact, separated by ∼300 nm (λ/2) from the true signal. **(d)** The same region shown in (c), after fitting the data with a PSF model with the correct phase. The ghost image is no longer present, and the histogram (right panel) shows only two peaks corresponding to the Nup96 organization in the NPC. **(e)** Illustration of the phase-shift transform. Beginning with the original PSF scan (one channel shown), the PSF is decomposed into envelope (*h*_env_) and modulation (*h*_mod_) components. A Hilbert transform is used to obtain a 90-degree shifted modulation component 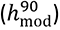. The components are linearly combined with a phase factor Δ φ, to obtain the 4Pi PSF with an arbitrary phase shift. **(f)** Measurement of the cavity phase. Three sets of localizations from narrow time windows (*t_1_, t_2_, t_3_*) were fit with PSF models having a range of phases spanning 360 degrees, and the mean chi-square for each set of fit results is plotted vs. the model phase. The plots exhibit a single minimum at the true phase of the optical cavity in that time window. **(g)** Evolution of the cavity phase. Repeating the analysis shown in (e) for the entire dataset yields the time evolution of the cavity phase over the course of an experiment. The three time-points from (e) are indicated on the plot. In this example, each time window consists of 250 localizations, and the phase measurement precision was < 0.03 radians (see Supplementary Fig. S15).

The effects of PSF phase drift can be seen in the 4Pi-STORM image. Figure 2b shows nuclear pore complexes (NPCs) in which the nucleoporin Nup96 has been labeled with Alexa Fluor 647. The data was first analyzed using the spline model obtained from the bead scan recorded at the start of the experiment. Inspection of the *x-z* profile of the image (Fig. 2c), however, shows a large fraction of localizations assigned incorrectly by the fit, resulting in a strong double image artifact. The upper and lower rings of the core region of the NPC appear twice, as evidenced by the *z-*profile of the dataset (Fig. 2c, right panel). We verified the effect of PSF phase drift using simulations, which showed a strong increase in localization artifacts when the PSF model phase is shifted by > 60 degrees from the true phase of the optical cavity (Supplementary Fig. S11).

To account for the time dependence of the optical system, we implemented a method for numerically shifting the phase of the measured PSF, as shown schematically in Fig. 2e. Using a Fourier low pass filter, we first decomposed the measured PSF into a static envelope function (*h_env_*) and a modulation component (*h_mod_*). Next, a Hilbert transform applied along the *z*-dimension was used to obtain a modulation component shifted by 90 degrees 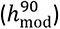. The three components (Eqs. 3-5) can then be recombined, while introducing a phase shift Δφ, to yield a new PSF model 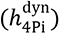 with an arbitrary modulation phase (Eq. 6).

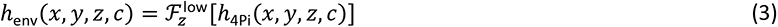

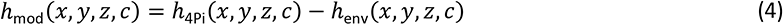

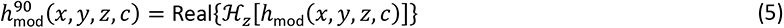

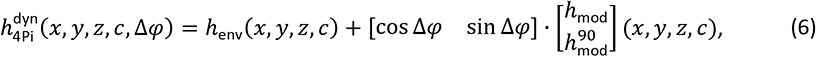

This approach makes no assumptions about the PSF structure, and can be applied to any PSF having a periodic component along the *z*-axis. We verified that the results of our algorithm correspond to physically relevant PSFs, by comparing phase-shifted PSFs with measured PSFs whose phase was altered manually, by changing the length of one cavity arm (Supplementary Figs. S13, S14). Numerical phase shifting of a measured 4Pi PSF is shown in Supplementary Video S1.

Surprisingly, we found that the phase-shifting algorithm also allows a simple means of measuring the PSF phase directly from the data. Taking subsets of molecule images corresponding to short time windows of the experiment, we fit the images with a series of phase-shifted PSF models. The mean value of chi-square 〈χ^2^〉 for each set of fits provides a measure of how well the model describes the data. After varying the phase over 360 degrees, we obtained plots of 〈χ^2^〉 versus phase for three different time windows (Fig. 2f). Each curve exhibits a single minimum corresponding to the PSF phase at that time point.

The full temporal evolution of the PSF phase can be measured using this approach. We found that as few as 250 emitter images were sufficient to measure the phase with a precision < 0.03 radians, allowing for a time resolution as high as 1 second for our data (Supplementary Fig. S15). The time-dependent PSF phase over the course of a full image acquisition, with a duration of 800 seconds, is plotted in Fig. 2g. The phase exhibits short- and long-timescale fluctuations, which introduces a corresponding bias in the estimated *z-*coordinates of the fluorophores (Fig. 2g, right axis).

Knowing the evolution of the cavity phase, we shifted the phase of the initial bead scan to the mean phase estimated for the experiment, and re-analyzed the Nup96 dataset using this new PSF model. Phase drift was corrected by applying a time-dependent *z-*shift to the molecule coordinates which is proportional to the measured phase (Fig. 2g, see Methods). This resulted in a dramatic reduction in localization artifacts in the image data (Fig. 2d).

By including the time-dependent phase as an adjustable parameter of the spline representation, both the structure and dynamics of the 4Pi PSF can be modeled accurately. If left uncorrected, phase fluctuations introduce errors in the localization *z-*coordinates, and large phase shifts cause phase-wrapping artifacts. We refer to our model as a Dynamic Spline PSF (DS-PSF), as it combines the strengths of a parametrized analytic description with a numerical model of the experimentally measured PSF structure.

### Localization Precision and Accuracy

We evaluated the performance of the microscope and the dynamic spline analysis by determining the localization precision and the frequency of localization artifacts. For this, we recorded 4Pi-STORM images of single molecules of Alexa Fluor 647, in which each fluorophore was localized multiple times. Localization clusters for 305 molecules with at least 10 localizations each were co-aligned, and the resulting distribution of position measurements (Fig. 3a) provided an estimate of the measurement precision^9,20,21^. The position distribution has a 3D Gaussian profile with a standard deviation of 3.4 nm in the *x-* and *y*-dimensions, and 2.1 nm in the *z*-dimension (Fig. 3b). The mean number of photons per switching event was 8002 (Supplementary Fig. S17).

**Fig. 3.**
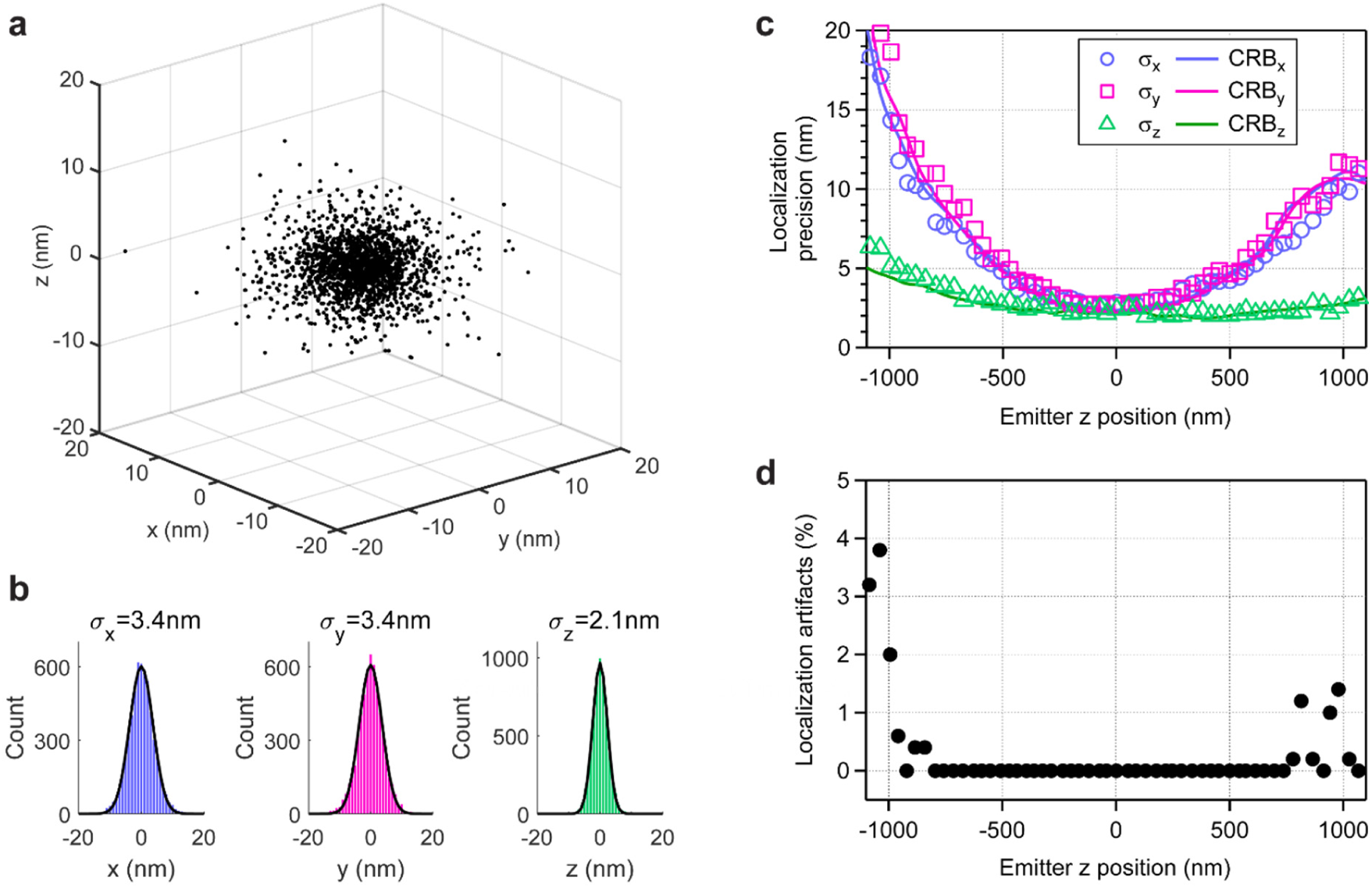
Localization precision and artifacts. **(a)** Distribution of 3D position measurements for single Alexa Fluor 647 molecules, determined using the dynamic spline PSF model (DS-PSF). The size of the distribution provides a direct measure of the localization precision of the microscope. On average, 8002 photons were detected per fluorophore switching event. **(b)** A 3D Gaussian fit to the histogram of the localization distribution yields a standard deviation of 3.4 nm in the *x-* and *y*-dimensions, and 2.1 nm in the *z-*dimension, corresponding to a spatial resolution (FWHM) of 8 nm laterally and 5 nm axially. **(c)** Variation of the localization precision as a function of the *z-*coordinate of the emitter. By stepping a fluorescent bead through the focus of the microscope, the measurement precision was measured over a 2.2 µm range. The bead brightness was similar to a single fluorophore (9200 photons per frame). The Cramér-Rao lower bound, which estimates the theoretical limit of the precision based on the information content of the data, is also shown (solid lines). **(d)** Fraction of localizations erroneously assigned to the wrong interference fringe by the DS-PSF fit, as a function of the *z-*coordinate of the emitter. When such an error occurs, the *z-*position of the emitter is apparently shifted by a multiple of the PSF fringe spacing (λ/2), and this results in periodic ghost artifacts in the final image. Over a central range of ∼1.5 µm, no localization artifacts were measured.

Next, we investigated the localization of emitters outside the focal plane. A fluorescent bead was scanned over a range of 2.2 µm, with the illumination adjusted such that ∼9200 photons were detected per camera frame, similar to a single fluorophore. By scanning the bead in steps of 20 nm (Supplementary Fig. S18), the precision could be measured at each position. We found that the dynamic spline fit reached the optimal precision in three dimensions, as measured by the Cramér–Rao bound (CRB), over the full range of the scan (Fig. 3c), and exhibited no measurable bias (Supplementary Fig. S19).

The same measurement was used to determine the frequency of localization artifacts, which appeared at multiples of ±λ/2 above or below the true *z*-position in the scan. Using the dynamic spline analysis, we found that such artifacts were almost entirely avoided. Over a range of 1.5 µm around the focus, we measured an artifact fraction of 0 %, rising to 5-10 % for positions >1.0 µm away from the focal plane (Fig. 3d).

The localization precision in the focal region (Fig. 3a,b) corresponds to an effective spatial resolution of 8 nm in the lateral (*x*-*y*) direction, and 5 nm along the optical axis (*z*). As a point of comparison, we note that the measured *z-*precision is approximately ten-fold higher than for a standard astigmatic 3D STORM microscope^21^, highlighting the advantages of the 4Pi approach.

### Comparison with Astigmatic 4Pi

In previous implementations of 4Pi-SMLM, astigmatic imaging has been employed as a means to avoid localization artifacts^13,16,22,23^. This method breaks the symmetry of the PSF along the *z*-axis, and allows the interference fringe to be identified according to the ellipticity of each emitter image. To introduce astigmatism into the optical system, hyperbolic curved mirrors^16^, or electronic deformable mirrors^13,22,23^, are incorporated in the 4Pi cavity.

The astigmatic 4Pi approach has drawbacks, however. First, astigmatism causes the lateral extent of the PSF to increase dramatically for emitters positioned above or below the focal plane. This requires a larger area on the CCD detector per emitter image, and can effectively limit the maximum density of fluorophores on the sample, reducing the achievable resolution^24,25^. Second, the specialized mirrors required to shape the PSF add significant complexity and cost to the (already non-trivial) opto-mechanical design of a 4Pi microscope, and add a potential source of instability. Conversely, our measurements of the localization artifact frequency (Fig. 3d), obtained using an unmodified, symmetric 4Pi PSF, suggest that the dynamic spline approach renders astigmatism unnecessary.

In order to compare the methods, we created simulated data based on theoretical models of the symmetric 4Pi PSF and the astigmatic 4Pi PSF (Fig. 4a and Supplementary Fig. S20). For each case, we simulated single fluorescent emitters over a range of *z*-positions, for typical experimental conditions. The data were analyzed with either the dynamic spline model (symmetric PSF) or with the ellipticity-phase analysis^13,22^ (astigmatic PSF). The 3D localization precision obtained for the two analyses are shown in Fig. 4b. Close to the focal plane, both methods yield similar lateral and axial localization precision (3-5 nm in *x-y*, and 2-3 nm in *z*). For |*z*| ≳300 nm, however, the astigmatic 4Pi PSF becomes strongly elliptical, spreading the detected photons over a larger area of the camera, causing the lateral localization precision (in the *x* or *y* dimension) to deteriorate sharply. In contrast, the symmetric 4Pi PSF, in combination with the DS-PSF analysis, exhibits a higher localization precision which is relatively insensitive to the emitter *z*-position. The observed difference is not primarily due to the PSF shape, but results because the DS-PSF more accurately models the data. To support this view, we also created a cubic spline model of the astigmatic PSF, and fit the simulated astigmatic data with this model. We found that using the spline analysis for both cases yields more comparable results (Supplementary Fig. S22).

**Fig. 4.**
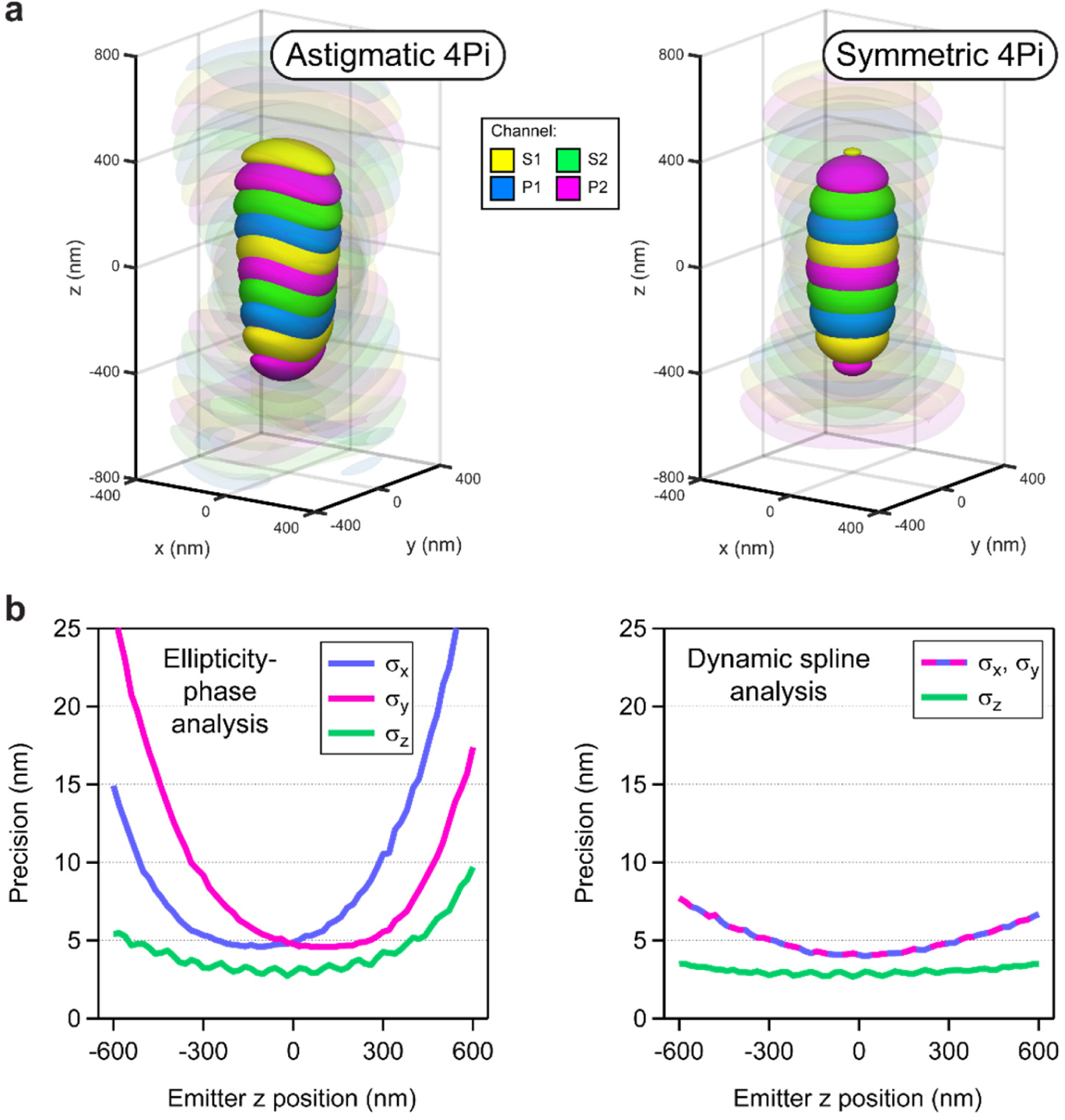
Comparison with Astigmatic 4Pi PSF-based approaches. Previously reported 4Pi-SMLM microscopes employ a shaped astigmatic 4Pi PSF, together with an analysis which measures both the ellipticity and the interference phase of the fluorophore images, to help avoid localization artifacts. (**a**) Side-by-side comparison of simulated astigmatic 4Pi and symmetric 4Pi PSFs, rendered as four-color isosurfaces at 35% and 8% of maximum intensity. **(b)** Localization precision comparison for the astigmatic 4Pi PSF using the ellipticity-phase analysis^13,22^ (left), and the symmetric 4Pi PSF using the dynamic spline analysis (right). Simulated fluorophore images were generated based on the two PSFs, using realistic photon counts (mean 8000 photons) and signal to noise ratio (10 background photons per pixel). The data were analyzed with the indicated method (see Methods and Supplementary Note 3). The results show that while both approaches reach a similar localization precision at the focal plane (*z*=0), for the astigmatic case the precision falls off sharply for positions further than 300 nm from focus. In contrast, the dynamic spline fit maintains a relatively constant precision over the full *z*-range of the simulation. The frequency of localization artifacts for the two methods is evaluated in Supplementary Fig. S21.

Current developments have also followed the direction of numerical PSF models which better describe the data. For example, by including the phase as a free parameter of a 4Pi PSF model, a recent study examined the potential of fitting both the phase and the *z*-position independently for individual fluorophore images in simulated datasets^26^. Also, phase retrieval was used to estimate the aberrations present in each path of a 4Pi optical system^23^, allowing an analytic model of the astigmatic 4Pi PSF which is more accurate than the previous astigmatism analysis, and obtains similar results to the spline fit in terms of localization precision. The dynamic spline method adopted here is unique, however, in its capacity to precisely measure and compensate 4Pi PSF phase evolution (Fig. 2g), as well as being a simple and direct approach.

### 3D imaging of Beta-II Spectrin

To characterize the capabilities of the microscope and the analysis pipeline in real applications, we imaged primary neuron cultures stained with antibodies against beta-II spectrin, a component of the membrane-associated periodic cytoskeleton (MPS)^27,28^. Viewed from above (*x-y* projection), the spectrin images show a characteristic pattern of stripes oriented orthogonally to the axis of the neuronal process (Fig. 5a and Supplementary Fig. S23). Closer inspection of the 3D data reveals spectrin molecules surrounding the axonal circumference, with few puncta inside the axon (Figs. 5b-e). At the surface of the axon, spectrin appeared in some regions as a linear series of double spots (e.g. white arrow, Fig. 5e). A line profile through one such doublet is shown in Fig. 5f, revealing two peaks separated by 22 nm, which is compatible with the known structure of spectrin heterotetramers and the epitope recognized by the antibody^29^. Notably, each view (*x-y*, *x-z*, and *y-z*) shows equivalent structural detail, due to the near-isotropic localization precision (see Supplementary Videos S2 and S3).

**Fig. 5.**
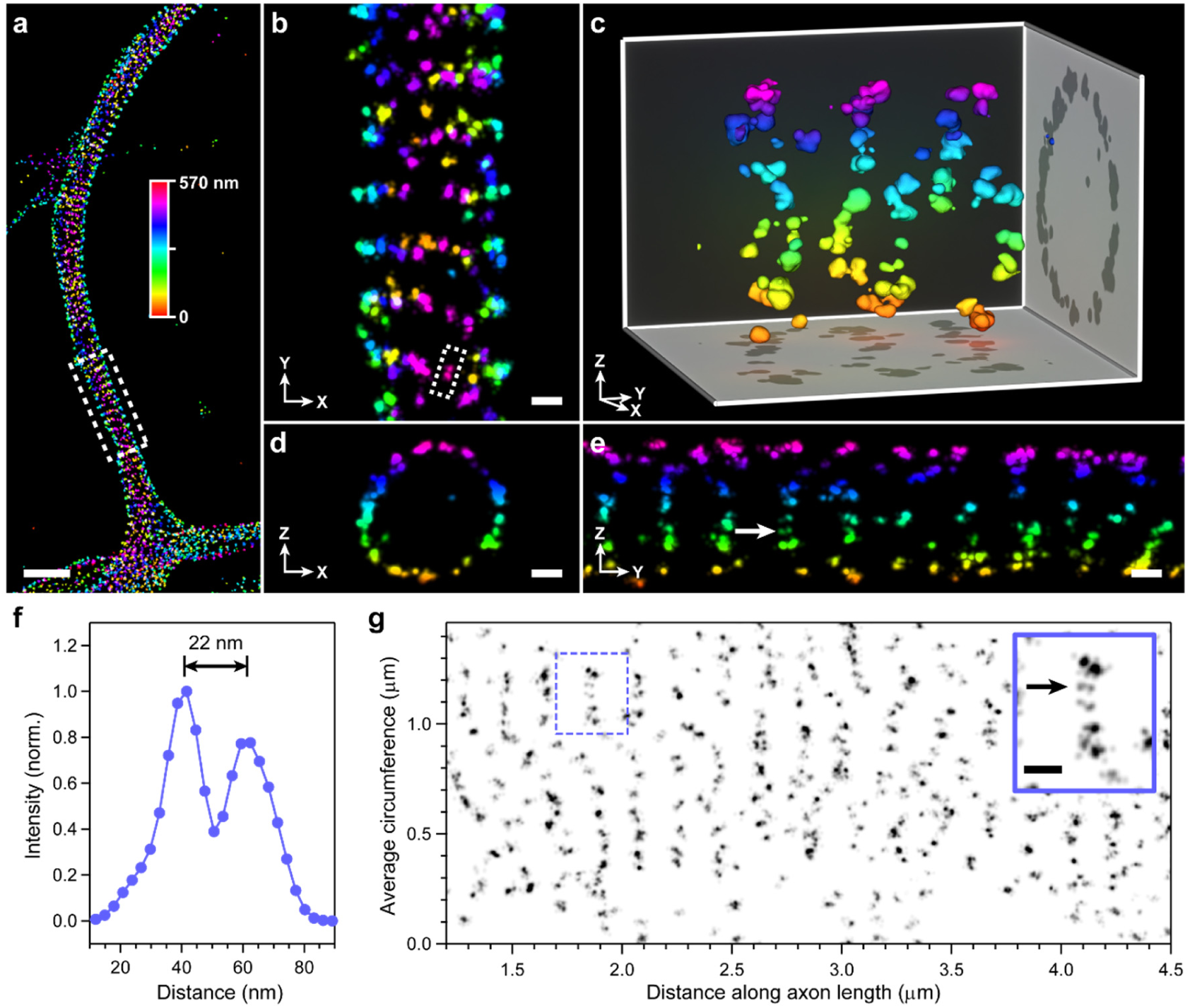
4Pi-STORM imaging of the neuronal cytoskeleton. Beta-II spectrin in the axon of a primary neuron cell, labeled with primary and secondary antibodies. Spectrin forms ring-like patterns around the circumference of the axon, oriented orthogonally to the neuron axis. **(a)** Top view (*x-y*) of the axon, showing periodic spectrin stripes. Localizations have been colored according to their *z-*coordinate, as indicated by the color bar. Scale bar 1 µm. **(b, c, d, e)** Magnified views from the boxed region in (a). Panels (b, d, e) show the neuron as viewed from the top (b), along its axis (d), and from the side (e). The near-isotropic resolution of the microscope allows these views to contain equivalent levels of detail. Close inspection of the top view (a) shows that some stripes are angled relative to the axon axis, appearing to form a spiral. The view along the axis (d) shows the continuity of the ring structure around the circumference of the axon, and the absence of spectrin in the axon interior. Seen from the side, within each stripe the spectrin signal appears to form doublets (panel e, white arrow), which also appear in the top view (panel b, white box). Panel (c) shows a 3D perspective view of a short section of the axon. Scale bars: 100 nm. **(f)** A line profile through a spectrin doublet (panel b, white box) shows a spacing of 22 nm between the spectrin peaks. **(g)** The spectrin signal as a function of its angular position around the axon circumference (y-axis), plotted along the length of the axon (x-axis). This view is equivalent to “unwrapping” the spectrin into a 2D plane, and allows inspection of the spectrin organization with respect to the membrane. Spectrin doublets are visible in the plot (inset, black arrow). Some rings are curved as they wind around the axon, forming a spiral-like structure (e.g. the region from 1.5 µm to 2.5 µm). The top of the axon is located at the midpoint of the *y*-axis, and the bottom of the axon is at position 0.0. The mean circumference of the axon in this region was measured to be ∼1.4 µm. The inset (solid blue box) shows a magnified view of the region in the dashed blue box. Inset scale bar: 100 nm. Longer sections of unwrapped views from this sample are shown in Supplementary Figs. S25 and S26.

We analyzed the topology of the spectrin lattice by unwrapping the curving neuronal membrane onto a 2D plane. A plot of the spectrin density around the circumference of the neuron, versus its position along the neuron’s length, yields a 2D view of the membrane which is simpler to visualize (Fig. 5g, Supplementary Figs. S24-S26, Supplementary Video S4). This revealed details in the local organization of the MPS, such as gaps which were observed at the bottom of the axon, and Y-joints, kinks, and bends, possibly indicating the presence of a splitting actin meshwork, or for localized regulation of the tension and tilting of the MPS^30,31^. Intriguingly, we observe that within the space of few micrometers the topology of the MPS can transition from an ordered series of perpendicular ring shapes, to a seemingly disordered pattern, to a tilted organization which is consistent with a 3D spiral. A short section of this spiral-like structure can be seen in the 3D perspective view (Fig. 5c).

The depth-of-field of the microscope, defined as the *z-*range over which the resolution does not appreciably degrade, is approximately 1 µm (see Fig. 3). Cellular samples are typically several micrometers thick, however, requiring a larger imaging depth. We addressed this issue by implementing step-wise *z-*stage scanning^13,32^. During the measurement, the sample may be periodically shifted in *z*, with a step size of ∼500nm, over a range of 4-5 µm. Although this procedure reduces the numbers of localizations in each plane due to expenditure of switching cycles for undetected fluorophores, we found it to be a simple and effective means of imaging moderately thick specimens. Supplementary Fig. S27 shows an example of a thick neuronal process, imaged over 10 *z-*planes for a total depth of approximately 5 µm.

### Multicolor imaging of Mic60 and Mitochondrial Nucleoids

The mitochondrial inner membrane forms numerous invaginations, with circular or slit-shaped openings, termed crista junctions. Crista junction formation requires the Mitochondrial Contact Site and Cristae Organizing System (MICOS), a large multi protein complex, which is embedded in the inner membrane. The nanoscale spatial organization of Mic60, the core component of MICOS, was recently investigated in human bone osteosarcoma epithelial cells (U-2 OS) using 3D MINFLUX nanoscopy^33^. Using 4Pi-STORM, we imaged Mic60 in U-2 OS cells and monkey kidney fibroblast-like cells (COS-7), to gain insight into the MICOS organization in different cell types.

In U-2 OS cells, Mic60 exhibited a 3D distribution which is mainly punctate (Figs. 6a,b). After determining the 3D envelope of the organelle, we unwrapped the data onto a 2D plane (as before for the MPS) making it easier to analyze its overall distribution (Fig. 6e). The data show that Mic60 puncta consist of 2-6 localization clusters spaced 20-30 nm apart, sometimes arranged as pairs in a short line, or sometimes forming circular groups approximately 30 nm in diameter (Figs. 6b,e). These data further substantiate the view that in human U-2 OS cells, multi-protein arrangements of Mic60 surround, or line, individual circular crista junctions^33^.

**Fig. 6.**
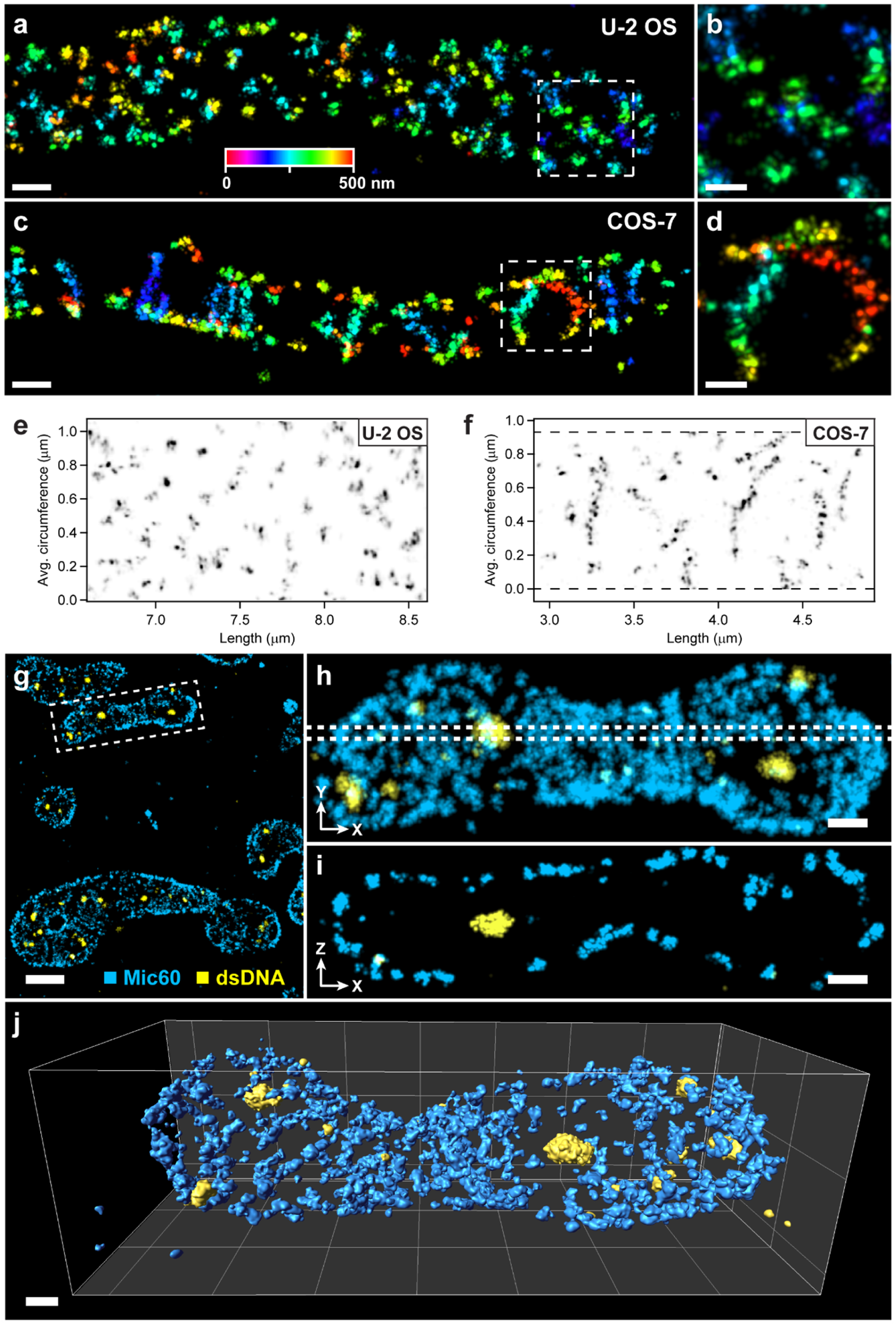
Single- and Multicolor imaging of Mic60 and mitochondrial DNA. Mic60 is a component of the MICOS complex, and is involved in the formation and maintenance of crista junctions between the mitochondrial inner membrane and boundary membrane. **(a)** Mic60 in a U-2 OS cell, labeled with primary and secondary antibodies. The localizations are color-coded according to their *z-*coordinate (see color bar). Scale bar 200 nm. The christa junctions appear as structured, punctate clusters, less than 50 nm in size. **(b)** Magnified view of the boxed region in (a). Scale bar: 50 nm. **(c)** Mic60 in a COS-7 cell, in which the christa junctions exhibit a qualitatively different organization. Here, Mic60 localizations appear in stripes spanning segments of the boundary membrane, approximately 200 to 600 nm in length. Scale bar: 200 nm. **(d)** Magnified view of the boxed region in (c). Scale bar: 50 nm. **(e, f)** Unwrapped views of the Mic60 localization density around the surface of the mitochondria, showing the nanoscale structure of Mic60 clusters and stripes. In U-2 OS cells Mic60 clusters appear predominantly punctate, with pairs or clusters of signal density separated by ∼20-40 nm (Supplementary Figs. S29, S30). In COS-7 cells, stripes of Mic60 appear to have a zig-zag or double-line structure, with a typical width of ∼20 nm (Supplementary Figs. S32, S33). Dashed lines indicate the extent of the data in panel (f). **(g)** Two-color image of Mic60 (blue) and mitochondrial nucleoids (yellow) in a COS-7 cell, stained with antibodies labeled with Alexa Fluor 647 and Cy5.5, respectively. Scale bar: 1 µm. **(h)** Detail view of the boxed region in (g). Mic60 density is lower close to the DNA signal, suggesting a lower number of crista membranes in these regions. **(i)** Cross-section (x-z) slice through the region indicated by the dashed lines in (h), showing Mic60 at the inner boundary membrane surface, and a DNA cluster in the center of the mitochondrion. **(j)** A 3D perspective view of the mitochondrion shown in (h) and (i), where the Mic60 and DNA have been rendered as isosurfaces. Scale bars: 250 nm (h, i, j).

In contrast to the punctuate distribution in U-2 OS cells, we observed that Mic60 often forms extended stripe-like arrangements in the mitochondria of COS-7 cells (Figs. 6c,d). In these cells, Mic60 appeared in linear segments of varying length, and in some places these stripes appeared to be made up of paired clusters of localizations, separated by approximately 25 nm (Figs. 6d,f). The data demonstrate differences in the sub-mitochondrial organization of MICOS in different cell types. Presumably, these differences in the Mic60 distribution have an immediate impact on the cristae structure in these cells. Further visualizations of Mic60 in the two cell types are shown in Supplementary Figs. S28-S33 and Supplementary Videos S5-S6.

The capability for multicolor, target-specific imaging allows the spatial organization of multiple cellular components to be visualized. We implemented multicolor 4Pi-STORM using a ratio-metric detection system^12,34^, and fit the localization data with fluorophore-specific DS-PSF models. This allowed two photo-switchable fluorophores, Alexa Fluor 647 and Cy5.5, to be accurately localized with no significant loss of precision compared with single color imaging, and efficiently discriminated with a color identification accuracy of approximately 93% (see Methods and Supplementary Figs. S34-S39).

We demonstrated multicolor imaging using COS-7 cells labeled for Mic60 and double stranded DNA. Mitochondrial DNA (mtDNA) is compacted into nucleoprotein complexes denoted nucleoids, which form ellipsoid-shaped clusters of approximately 100 nm in size^35,36^. The dual-color images revealed mitochondria with peripheral, stripe-like distributions of Mic60, and dense clusters of DNA within the mitochondrial volume (Figs. 6g-i). Notably, the density of Mic60 labeling was reduced close to the nucleoids, indicating a lower number of cristae in those regions^37^. A detailed view of a single mitochondrion, together with a thin *x-z* cross-section slice (Figs. 6h,i), shows Mic60 localized to the inner boundary membrane, while nucleoids lie within the mitochondrial interior (see Supplementary Video S7).

## Discussion

The strength of the 4Pi concept lies in the structure of the PSF. Self-interference of fluorescence photons, collected from both sides of the sample, introduces a sharp modulation along the optical axis, making the detected signal extremely sensitive to the 3D coordinate of the emitter^10^. Under conditions of spatially invariant illumination, the 4Pi PSF yields the highest localization precision per detected photon of any optical detection scheme^38^.

In this study we developed an analysis framework which makes full use of the information content of 4Pi-STORM data by accurately modeling the PSF structure and dynamics, thereby obtaining optimal localization precision and accuracy. The dynamic spline PSF model allows the precise measurement and correction of phase transients, which is essential for accurate 3D fluorophore localization at single nanometer length scales. For the conditions we tested, we found that the symmetric 4Pi PSF is ideal, achieving a higher localization precision and lower artifact rate compared with the astigmatic 4Pi PSF. Together, these developments enable robust and accurate 4Pi-STORM imaging, with a less complex instrument, and may also facilitate performance improvements for other techniques such as PAINT-based super-resolution microscopy^39^. The software tools developed for this work, including a GPU-accelerated multichannel spline fitting library, and algorithms for the creation of new spline models from 3D data, are published as open-source projects.

The 4Pi-STORM microscope enables the visualization of new structures in biological samples. Images of beta-II spectrin in primary neurons (Fig. 5) showed the nanoscale organization of spectrin, and allowed the neuronal membrane to be mapped such that the topology of the spectrin rings could be digitally unwrapped and inspected. Images of crista junctions in mitochondria (Fig. 6) revealed distinct patterns of Mic60 in different cell types, and resolved the substructure of Mic60 stripes and puncta. We adapted our analysis for multicolor imaging, which is essential for studying the relative organization of multiple cellular components. The recently developed salvaged fluorescence method for 4Pi-SMLM^22^ would complement the methods described here by improving photon collection efficiency and multicolor discrimination. We note that the “salvaged” signal could potentially be integrated as a channel of the DS-PSF model, maximizing the information obtained.

## Methods

### Microscope

#### Microscope design

The 4Pi-STORM microscope was designed and built using an optical layout based on earlier concepts^11,12^. The mechanical design includes new mounting and positioning systems for the objective lenses, mirrors, beam splitters, and sample; a larger interferometric cavity; and the addition of a feedback control system for sample focus. The horizontal layout of the microscope also allows for simpler construction and modification as compared with systems built on a vertically oriented optical breadboard.

#### Objective lenses

The microscope uses two silicone oil immersion objective lenses (Olympus UPLSAPO100XS) with a focal length of 1.8 mm (magnification 100X), a numerical aperture (NA) of 1.35, and a working distance of 0.2 mm. One objective lens (Obj. A) is fixed in position, mounted in a threaded cylinder which is clamped into a V-block (Supplementary Fig. S6). The second objective lens (Obj. B) is fully adjustable in terms of *x*, *y*, *z*-position and tip/tilt. Obj. B is mounted inside the aperture of a 3-axis piezo stage (Physik Instrumente P-733.3DD) as shown in Supplementary Fig. S7. In this arrangement, the piezo stage allows fine control of the objective position in *x*, *y*, and *z*. The entire assembly holding Obj. B is mounted on sliding rails which allow it to be retracted for sample insertion and extraction.

#### Interferometric cavity and fluorescence detection

Fluorescence from the sample travels simultaneously along the upper and lower arms of the 4Pi interferometric cavity. In each of the two arms, the light passes through the objective lenses, dichroic mirrors (Chroma, ZT405/488/561/640/950RPC), quarter-wave plates (B. Halle, RAC3.4.15L), and modified Babinet-Soleil compensators (B. Halle) before reaching the 50:50 beamsplitter (B. Halle TWK1.25), as shown in Supplementary Fig. S1. The dichroic mirrors allow the excitation light and control beams to be introduced into the system. The quarter-wave plates are set at an angle of 45 degrees relative to the plane of the microscope, ensuring that fluorescence photons from the sample are equally distributed across all polarization angles. The compensator in the upper arm of the cavity (QC, Supplementary Fig. S1) consists of a fixed plate of BK7 glass and two adjustable wedges of quartz glass, such that the total thickness of quartz can be varied using a micrometer screw. This element allows a variable phase delay to be introduced between the s-polarized and p-polarized components of the fluorescence. In the lower arm, the compensator (GC) consists of a fixed plate of quartz glass and a variable thickness of BK7 glass, which can be adjusted to equalize the overall optical path length of the upper and lower arms and to correct for dispersion in the cavity. The compensators were adjusted such that the overall phase delay between the s- and p-polarized fluorescence was 90 degrees. The 50:50 beam-splitter was mounted on a custom-made linear translation stage which incorporates a micrometer screw for coarse translation, adjustments for tip/tilt, and a linear piezoelectric actuator (Physik Instrumente P-841.1) for fine adjustment of the beam-splitter position.

The tube lenses (Linos, Achromat VIS, *f* = 500 mm; position L1 in Supplementary Fig. S1) form an image of the sample which was masked by an adjustable rectangular aperture (OWIS SP60). After passing through the polarizing beam-splitter (B. Halle, PTW 1.25) the fluorescence is split into four spatially separable components, consisting of two s-polarized and two p-polarized channels, each with a different interference phase. The telescopes formed by lenses L2 (Linos, Achromat VIS, *f* = 300 mm) and L3 (Linos, Achromat VIS, *f* = 250 mm) form four images of the sample on four quadrants of the EMCCD detector (Andor, Ixon Ultra 897). The effective magnification of the imaging system, including the objective lenses, tube lenses, and telescopes, was 231.48X. Immediately before the camera, the fluorescence was filtered with a combination of spectral filters (Chroma, HQ70075; Semrock, LP647RU and FF01-770SP). The four images formed on the four quadrants of the camera, numbered S1, P1, S2, and P2, correspond to a relative interference phase of 0, 90, 180, and 270 degrees, respectively, such that a fluorophore which interferes constructively in channel S1 exhibits destructive interference in channel S2, and so on.

#### Multicolor imaging configuration

When the microscope is configured for multicolor imaging, two additional filters are added to the detection path. The microscope was designed to discriminate fluorophores with strongly overlapping emission spectra by means of ratiometric spectral detection^12,40^. Two dichroic mirrors (Semrock, FF685-Di02), placed at an acute angle of 34 degrees with respect to the fluorescence beam path, serve as polarization selective long-pass spectral filters. The placement of the filters (DM3) is shown in Supplementary Fig. S1, and the filter spectra are shown in Supplementary Fig. S34. In this design, the ratio of s- to p-polarized photons detected on the camera is approximately 0.6 for Alexa Fluor 647, and 0.9 for Cy5.5, and this ratio may be used to distinguish the two dyes in the data analysis step. In the multicolor configuration, the bandpass filter (Chroma HQ70075) was removed from the detection path.

#### Sample mounting and positioning

Samples were prepared on 18mm diameter glass coverslips (Marienfeld, #1.5H) with a thickness of 170 ± 5 micrometers. Before use, the 18mm coverslips were sputter-coated with a thin aluminum layer on one side, covering one quarter of the glass surface. This coating forms a mirror which was used for initial alignment of the sample and microscope objectives. For imaging, biological samples were prepared on the coated side of the coverslip. The 18mm coverslip holding the sample was mounted onto a second, larger coverslip. For this purpose, 30mm glass coverslips (Marienfeld, #1.5H) were glued onto a custom support consisting of a stainless steel disc with an open aperture. Sample coverslips were then mounted in a sandwich configuration, by placing a droplet of imaging buffer onto the 30mm coverslip, flipping the sample coverslip onto the droplet, and removing excess buffer with vacuum suction. The coverslip sandwich was then sealed around the edge using fast-curing epoxy glue (Uhu). The assembly of the two coverslips, supported by the steel disc, was then magnetically coupled to a sample holder which could be connected to the sample translation stage. An isometric view of the stage assembly if shown in Supplementary Fig. S8.

The sample holder was mounted onto a three-axis translation stage (Newport, M-462-XYZ-M), which was coupled to two servo motors (Thorlabs, Z825B) for motorized sample positioning in *x* and *y*, and to a manual micrometer for initial focusing in *z* (parallel to the optical axis). Fine focus control in *z* was provided by a linear piezo stage (Physik Instrumente, P-752) mounted underneath the top section of the translation stage.

#### Excitation light path

A custom-built laser launch provided light for fluorophore excitation and activation. The following lasers were used for this purpose: Red (642nm, 1.5W, MPB Communications, 2RU-VFL-P-1500-642), Green (560nm, 5W, MPB Communications, 2RU-VFL-P-5000-560), Blue (488nm, 50mW, Coherent Sapphire), and Ultraviolet (405nm, 100mW, Coherent OBIS). The 642nm, 560nm, and 488nm beams were combined using dichroic mirrors and passed through an AOTF (AA optoelectronic) for power control and modulation. Control of the 405nm laser was done via software, and the 405nm beam was combined with the other beams after the AOTF. The four combined beams were coupled into a high-power polarization-maintaining single mode fiber (OZ optics) via an achromatic fiber coupler (Schäfter & Kirchhoff, 60FC-4-RGBV11-47).

The optical layout of the illumination path is shown in Supplementary Fig. S2. Light exiting the fiber is collimated by lens L5 (Linos, Achromat VIS, f=100mm), passes through an adjustable aperture, and is focused by lens L4 (Linos, Achromat VIS, f=500mm) to the back focal plane (BFP) of the fixed objective lens (Obj. A), such that light exits the objective as a narrow collimated beam. By placing the output of the fiber in a plane which is conjugate to the BFP, and translating the fiber laterally with respect to this plane, the angle of the excitation beam at the sample could be adjusted between epifluorescence and near-TIRF illumination modes. Typically, for 4Pi-STORM experiments, the illumination angle was adjusted to obtain the best signal to background ratio for a given sample.

#### Objective alignment control system

Alignment of the two objective lenses must be maintained throughout the course of a 4Pi-STORM experiment. To achieve this, a control system monitors the relative alignment of two lenses via an infrared laser beam, and continuously adjusts the three-axis piezo stage supporting Obj. B, to maintain its lateral and axial position with respect to Obj. A. The optical layout of the control system is shown in Supplementary Fig. S3. Laser light from a fiber-coupled laser diode (830nm, Thorlabs LP830-SF30) was collimated by lens L6 (Linos, f=40mm), and the size of the beam was adjusted by a variable aperture. The telescope formed by lenses L7 (Linos, f=150mm) and L4 expand the beam to a diameter of approximately 6mm. The dichroic mirror DM2 (Chroma, HC670SP) was used to couple the beam into the excitation path. After Obj. A, the beam is focused to a point inside the sample, and is then re-collimated by Obj. B (Supplementary Fig. S3) when the two lenses are co-aligned. The outgoing beam exits the cavity at DM1 in the lower cavity arm and enters the beam deflection analyzer, which consists of a quadrant photodiode (QPD), and two standard photodiodes located behind pinholes^12^.

#### Sample focus control

The distance between the sample and the fixed objective lens is actively monitored during the experiment, in order to correct for sample drift along the *z*-axis (the optical axis). For this purpose, a focus lock system was incorporated into the 4Pi-STORM microscope design. The system consists of an infrared laser (940nm, Thorlabs LP940-SF30) which is collimated and passes through a circular aperture, before being directed to one edge of the rear aperture of the fixed objective lens (Supplementary Fig. S4). A translation stage facilitates adjustment of the incoming beam position with respect to the objective. The beam emerges from the objective at an angle, focuses to the glass-water interface at the sample, and is reflected back into the objective lens. The beam is re-collimated, and returns along the same path until reaching the 50:50 beam-splitter, where part of the light is reflected and focused onto a CCD camera. Any change in the distance between the objective lens and the sample results in a lateral shift in the position of the returning beam, which is detected on the CCD. A software based feedback system detects this shift and corrects for it by adjusting the position of the piezoelectric element built into the sample stage.

#### Data acquisition and control systems

The microscope was controlled with custom software written in LabVIEW. Specifically, LabVIEW interfaces were written to control and monitor the EMCCD camera, excitation lasers, shutters, AOTF, piezoelectric translation stages, servo motors, focus lock CCD, QPD, photodiodes, and other electronic components. Higher-level interfaces and inter-process communication allowed the integration of multiple signals to implement feedback loops such as the focus lock and objective alignment control systems, and user interfaces allowing e.g. sample focus control and stage scanning. Analog voltage inputs and outputs, and digital I/O was managed with two DAQ cards (National Instruments PCI-6229 and PCI-6731).

### Sample preparation

#### Tissue culture and immunofluorescence

Experiments were performed using either standard COS-7 cells or U-2 OS cells (ATCC), or gene edited U-2 OS cells which express a SNAP-tagged version of the nucleoporin Nup107 (CLS Cell Lines Service, U-2OS-ZFN-SNAP-Nup107 clone #294) or Nup96 (CLS Cell Lines Service, U-2OS-CRISPR-NUP96-SNAP clone #33).

COS-7 cells were cultured in DMEM containing 4.5 g/l glucose and GlutaMAX™additive (Thermo Fischer Scientific, Waltham, MA, USA) supplemented with 100 U/ml penicillin and 100 μg/ml streptomycin (Merck Millipore,Burlington, MA, USA), 1 mM sodium pyruvate (Sigma Aldrich, St. Louis, MO, USA) and 10% (v/v) fetal bovine serum (Merck Millipore, Burlington, MA, USA). U-2 OS cells (ATCC, Manassas, VA, USA) were cultured in McCoy’s medium (Thermo Fischer Scientific, Waltham, MA, USA). The growth medium was supplemented with 10% FBS, 1% Pen/Strep, and 1% Sodium Pyruvate.

For antibody and SNAP staining, cell samples were plated on glass coverslips (18mm, #1.5H, Marienfeld) which were previously ¼ coated with an aluminum mirror surface (see above). Cells were grown in culture medium overnight. Prior to fixation, samples were washed twice in warm PBS.

For labeling of mitochondria, COS-7 cells were chemically fixed by adding 2 ml of a pre-warmed solution of 8% formaldehyde in PBS (137 mM NaCl, 2.68 mM KCl and 10mM Na2HPO4, pH 7.4, 37 °C) to 2 ml of the culture medium. After 5 minutes we exchanged the solution for 4% formaldehyde in PBS and incubated for 5 minutes. U-2 OS cells were chemically fixed by replacing the culture medium with 2 ml pre-warmed 4% formaldehyde in PBS for 5 min. The cells were permeabilized by applying 0.5% [v/v] Triton X-100 in PBS for 5 minutes and subsequently blocked with 5% [w/v] BSA in 0.1 M PBS/Glycine for 20 minutes. For antibody staining, we diluted primary antibodies against Mic60 (Proteintech, Rosemont, IL, USA, dilution 1:100), or double-stranded DNA (Abcam, dilution 1:3000) in 0.1 M PBS/Glycine containing 5% BSA [w/v]. The samples were incubated for 1 h at room temperature before washing six times with PBS to remove any unbound labels. Primary antibodies were detected using Fab fragments coupled to Alexa Fluor 647 (Thermo Fisher, dilution 1:500) or secondary antibodies coupled to Cy5.5 (Jackson Immunoresearch, custom labeled, dilution 1:200).

For nuclear pore complex labeling, U-2 OS cells were chemically fixed by replacing the culture medium with 2 ml pre-warmed 2.4% paraformaldehyde for 15 minutes, and briefly quenched with 50mM NH_4_Cl. The cells were then washed with PBS and permeabilized with 0.2% Triton X-100 for 15 minutes. After permeabilization, SNAP-tag expressing cells were washed three times in PBS for 5 minutes, followed by blocking in Image-IT (Thermo-Fisher, R37602). The cells were then incubated with the SNAP substrate (Alexa 647 – benzylguanine, 1 µM, NEB#S9136S) in 0.5% BSA + 1mM DTT at room temperature for 2 hours. Cells were then washed three times in PBS for 5 minutes, and stored in PBS.

#### Neuronal culture preparation and labeling

Cultures of dissociated rat hippocampal primary neurons were prepared from postnatal P0-P2 Wistar rats of either sex and cultured on glass coverslips coated with 100 µg/mL poly-ornithine (Merck KGaA) and 1 µg /mL laminin (BD Biosciences). Procedures were performed in accordance with the Animal Welfare Act of the Federal Republic of Germany (Tierschutzgesetz der Bundesrepublik Deutschland, TierSchG) and the Animal Welfare Laboratory Animal Regulations (Tierschutzversuchsverordnung). According to the TierSchG and the Tierschutzversuchsverordnung no ethical approval from the ethics committee was required for the procedure of sacrificing rodents for subsequent extraction of tissues, as performed in this study. The procedure for sacrificing P0-P2 rats performed in this study was supervised by animal welfare officers of the Max Planck Institute for Medical Research (MPImF) and conducted and documented according to the guidelines of the TierSchG (permit number assigned by the MPImF: MPI/T-35/18).

Cells were grown in presence of 1-β-D-Arabinofuranosyl-cytosin (Merck KGaA) at 37°C and 5% CO_2_. Cultures were fixed at 19 days *in vitro* in 4% PFA in PBS, pH 7.4, for 15 min and quenched for 5 min in PBS supplemented with 100 mM glycine and 100 mM ammonium chloride. Cells were permeabilised for 5 min in 0.1% Triton X-100, blocked with 1% BSA for 1 hour, and incubated with anti-beta-II spectrin primary antibody (BD Biosciences, cat. 612563, 1:200 dilution) overnight at 4°C. After washing in PBS, samples were incubated with secondary antibody (anti-mouse AlexaFluor 647 Fab Fragment, Thermo Fisher, A21237, 1:500 dilution) for 1.5 hours at room temperature and further washed. Samples were post-fixed in PFA 2% for 10 min, quenched for 5 min, rinsed, and mounted for imaging.

#### Calibration and alignment markers

Before mounting the sample, fluorescent beads were added on the surface of the coverslip, which act as fiducial markers for registration of the 4Pi image channels, and for the purpose of measuring the microscope PSF prior to imaging. Far-red fluorescent beads with a diameter of 100nm (Thermo-Fisher, F-8798) were added to the sample in PBS at a high dilution (1 : 2x10^6^), such that the beads were sparsely distributed on the coverslip surface. Beads were bound to the sample by briefly rinsing with 10mM Tris pH 8 + 50mM MgCl_2_. The sample was then washed three times in PBS and stored in PBS.

#### Imaging buffer

Prior to imaging, samples were mounted in STORM imaging buffer, consisting of Tris (50mM, pH 8.0), sodium chloride (10 mM), glucose (10 % w/v), β-mercaptoethanol (143mM, Sigma, M3148), and an enzymatic oxygen scavenger system (1% v/v). The enzymatic oxygen scavenging system was added to the buffer immediately before use, and the 100X stock solution was prepared by mixing pyranose oxidase powder^41^ (10 mg, Sigma, P4234) with catalase slurry (80μL, Sigma, C100) in PBS (170μL), and centrifuging the mixture at 13000rpm for one minute.

### Data acquisition

#### 4Pi-STORM image data acquisition

After mounting the sample, recordings were made in order to measure the registration parameters for the four image channels, and also to measure the PSF of the optical system. First, an isolated fluorescent bead was found on the sample (bound to the glass surface), and images of the bead were recorded at multiple positions in the field of view. This set of images defines the transform which maps the channels onto each other, and is used for co-registration. Next, an isolated fluorescent bead was centered in the field of view and scanned through the common focal plane of the objectives while recording a sequence of images on the camera. The depth of the scan was 3 micrometers, recorded over 3000 steps. The resulting bead image stack serves as an initial measurement of the PSF.

After locating a region of the sample to be imaged, the sample piezo stage was adjusted to bring the sample into focus and the focus lock control system was started. Next, the intensity of the red illumination was increased, causing the fluorophores to switch off and induce stochastic re-activation due to the red light. The angle of the illumination was adjusted to maximize the signal to background ratio of the image data. The EMCCD camera recording was started, with a typical recording duration of 80k-100k frames at 100Hz. The EMCCD was operated in 2x2 binning mode, making the effective pixel size 32 x 32 micrometers, corresponding to an effective imaging region of 138.2 x 138.2nm per pixel. During the measurement, the 405nm laser was switched on, and its intensity slowly increased, in order to maintain a constant rate of fluorophore switching events over time.

### Data analysis

#### Channel transform calculation

Images of fluorescent beads recorded at different positions in the field of view were used to determine a polynomial transformation, which relates coordinates in each of the image channels to every other channel. The 4Pi-STORM data consists of four image channels: two for S-polarized fluorescence and two for P-polarized fluorescence, numbered S1, P1, S2, P2. A sequence of images of bright fluorescent beads were segmented and fit as described above to determine the bead coordinates in each channel, creating a list of equivalent coordinate pairs between the channels. The POLYWARP function (IDL) was used to calculate a second-order (quadratic) polynomial mapping between each of the channel pairs using the coordinate list.

#### Image transformation

The raw STORM image stacks, and the PSF calibration stack, were transformed to map the four image channels to a common coordinate system. Making use of the polynomial mapping calculated in the previous step, we transformed the image data from each channel to the coordinate system of channel S1. This was done using the POLY_2D function (IDL), which employs cubic convolution for interpolation. After this step, image data from the four channels was precisely co-registered to within a root-mean-square deviation of 2 nm or less.

#### Calculation of spline coefficients for PSF model

The bead calibration image stack was converted to a set of cubic spline coefficients describing the PSF of the microscope. First, a 17x17 pixel region of the image stack, centered on the bead, was cropped out of the stack. This data was down-sampled along the *z-*axis by a factor of 10, using an averaging filter. Next, the stack was smoothed along the *z-*direction using a boxcar averaging filter with a spatial width of 50nm. After this procedure, the size of the image stack containing the bead scan was 17x17x300 pixels, corresponding to a physical volume of 2.35 x 2.35 x 3.00 micrometers.

This data was converted into a set of cubic spline coefficients, using standard linear algebra and an algorithm adapted from Babcock et al.^17^ The cubic spline PSF model has four channels, which were generated by running the spline coefficients calculator four times, once for each channel of the bead image data. A 3D cubic spline function has 64 polynomial terms for each volume interval, and therefore the total number of spline coefficients needed to describe the four-channel 17x17x300 voxel bead image stack was 4 x 64 x 16 x 16 x 299 ≈ 1.96 x 10^7^ coefficients. These values define an analytic function which represents a model of the microscope point spread function, and we refer to it here as the PSF spline. A new PSF spline was measured before each set of 4Pi-STORM measurements.

#### Numerical method for 4Pi PSF phase shifting

For the purpose of determining the phase of the interference modulation of the PSF, with respect to the sample focus, a method is needed to shift the phase of the PSF spline by an arbitrary amount.

The spline model of the PSF is based on experimental data, rather than an analytic model, and hence there is no phase parameter which can be simply adjusted. To dynamically shift the phase of the measured spline, we devised the following approach. First, the initial PSF spline was rendered as a four-channel 3D stack over the original range of the data, and with the original sampling frequency. This data was then divided into two components: a modulation component (*h*_mod_) which corresponds to the interference of the fluorescence, and a slowly varying envelope function (*h*_env_). The envelope function (Eq. 3) was measured using a Fourier step filter applied to each column of pixels in the PSF data stack, along the *z*-direction. For our measurements, the cutoff frequency of the filter was set to remove frequency components with a period shorter than 500nm, well above the oscillation period of the interference pattern (∼λ/2, where λ is the fluorescence wavelength).

The modulation component (Eq. 4) was determined by subtracting the envelope function from the original stack. To shift the phase, we used a Hilbert transform applied to the modulation stack, along the *z*-direction (Eq. 5). Taking the real part of the transform results in a new image stack 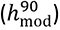 in which all sinusoidal components of the signal are shifted by 90 degrees. A linear combination of *h*_mod_ and 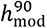 allows an arbitrary phase shift to be introduced, and the phase-shifted PSF is obtained by adding back the envelope function (Eq. 6). The dynamic spline PSF model (DS-PSF) is created by calculating the new set of spline coefficients corresponding to 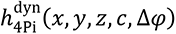.

#### Initial segmentation of 4Pi-STORM data

Initial 2D localization of fluorophore switching events was done by summing the four data channels (after transformation), resulting in an image stack which is effectively free of interference effects. This stack was segmented to identify initial peak candidates, and each peak was fit with a 2D Gaussian function to obtain an initial measure of its centroid, brightness, and width. Fluorophore switching events which persisted over multiple camera exposures were identified and grouped.

#### Single molecule localization with the DS-PSF model

Fluorophore switching events identified in the previous step were extracted from the image stack by cropping square 7x7 pixel regions containing the molecule images out of the transformed, four-channel raw data stack. Switching events which persisted for multiple frames were summed along the time axis. Each image was fit with the DS-PSF spline model, in order to find the optimal alignment in *x*, *y*, and *z* between the molecule image and the model, thus determining the 3D coordinate of the emitter. For this purpose, a multi-channel cubic spline model function was implemented for the Gpufit curve fitting library^19^ (see Supplementary Note 8). This model function accepts an a set of N-channel, 3D spline coefficients, and minimizes the sum of the squared deviation between the model and the data (chi-square, χ^2^), allowing the estimation of the 3D spatial offset, baseline, and amplitude of the spline function which best matches the emitter image. Using a PC with an Intel i7-5820K CPU and an Nvidia GTX 1080 GPU, the DS-PSF fits were executed at a typical rate of 80000 fits per second.

Due to the periodic nature of the PSF, multiple local minima exist in the chi-square landscape when fitting the PSF model to the emitter image. In order to sample the full parameter space of the fit, the algorithm was started with multiple initial *z*-offsets, spaced evenly along the *z*-axis, spanning the full range of the spline model. The spacing of the starting points was smaller than the periodicity of the PSF, ensuring that all of the local minima were sampled during fitting. The best fit between the spline and the emitter image was determined by selecting the fit result with the lowest chi-square value.

Accurate fit results depend on using a PSF model which correctly describes the experimental data, particularly with respect to the phase of the PSF modulation. For this reason, the time-dependent PSF phase was measured for each experiment, as described below. The PSF spline was then shifted to the mean experimental phase, and used for fitting the images of all emitters in the dataset.

#### Optical cavity phase determination

A set of emitter images acts as an effective reporter of the PSF phase at any time point during the experiment. To measure the phase, the measured PSF was used to generate a series of 12 phase-shifted PSF models, equally spaced in phase between 0 and 2π. The set of emitter images were then fit with each model, recording the mean chi-square value 〈χ^2^〉 obtained in each case. A plot of 〈χ^2^〉 vs. phase shift (Δϕ) exhibits a sinusoidal pattern, with a minimum at the phase corresponding to the experimental PSF phase. The position of the minimum was determined by fitting a sine function to the 〈χ^2^〉 vs. Δϕ data, as shown in Fig. 2.

The time evolution of the PSF phase was measured by repeating the phase determination for multiple time windows spanning the full dataset. The precision of this algorithm was dependent on the number of localizations per window, as shown in Supplementary Fig. S15.

#### Phase drift correction

In order to correct the effects of PSF phase drift, the proportionality factor between phase shift and apparent *z-*coordinate shift was determined for each measured 4Pi PSF in the following manner. Data slices taken from the PSF, corresponding to individual *z-*planes, were fit with phase-shifted versions of the full PSF, with phase shifts spanning a range of 2 radians. For each phase shift, the fit returned a shifted *z-*coordinate. A plot of phase shift vs. *z-*shift exhibits a linear relationship, and the slope of this curve yields the proportionality between *z-*shift and PSF phase. After determining the time-dependent experimental PSF phase as described above, phase drift was rescaled to *z-*drift, and subtracted from the *z-*coordinates of the localizations.

We note that the lateral variation of the PSF phase across the field of view (Supplementary Fig. S16) is typically small enough (+/- 40 degrees) so as not to present a problem with respect to localization artifacts. As long as the DS-PSF model used for fitting the localization data is within 60 degrees of the true detection PSF phase, no increase in artifacts will occur (Supplementary Figs. S11 and S12).

For datasets with large phase drift (greater than 60 degrees) over the course of a single measurement, localization artifacts can be avoided by dividing the dataset into time windows, and analyzing each with a DS-PSF model having a phase shift optimal for that window. This was not typically necessary for our data, with the exception of multi-step scanning measurements (see below).

#### Sample drift correction

Sample drift was measured by dividing the set of fluorophore localizations into time windows, rendering a 4Pi-STORM image for each time window, and calculating the 3D correlation function between images from different time points. The centroid of the peak of the correlation function was determined by fitting with a Gaussian function, and this coordinate represents the spatial offset between different time points, as described previously^20^. We used the robust cross correlation (RCC) approach, measuring the correlation between all pairs of time windows, to obtain the optimal 3D drift trajectory, which was then subtracted from the localization coordinates^42^.

#### Wavelength and refractive index corrections

Several factors necessitated corrections to the *z*-coordinates. First, we accounted for the difference in the mean wavelengths of the fluorescence detected from the calibration bead (712nm) and the fluorophores (Alexa Fluor 647: 680 nm, Cy5.5: 699 nm), which results in a small change in the period of the PSF modulation. The second correction is for the difference in the refractive indices of the sample (1.346) and the immersion oil (1.406). This is relevant due to the way in which the PSF was measured, by scanning the bead using the piezo stage rather than moving an emitter within the sample medium. Both corrections were implemented as a linear rescaling of the *z-*coordinates obtained from the DS-PSF fit.

#### Multi-step scanning data analysis

Multi-step stage scanning was used to record 4Pi-STORM images of thick regions of the sample, as shown in Supplementary Fig. S27. During the recording, the z-position of the sample stage was periodically shifted up or down by uniform increments in a staircase pattern, covering the depth of the desired imaging region. The typical step size was ∼400 nm, such that the 4Pi-STORM image data from neighboring steps would overlap partially, for later registration.

When the sample is shifted, the phase of the PSF also shifts, and this must be accounted for in data analysis. The approximate PSF phase shift Δφ, for a shift in the sample position by Δ*z*, is given by

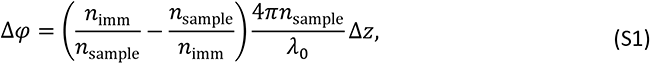

where *n*_imm_ and *n*_sample_ are the refractive indices of the immersion oil and the sample, respectively, and *λ*_0_ is the fluorescence wavelength in vacuum. This result is due to a sum of two effects, namely the shift of the focal plane when the sample is shifted, and the shift of the phase of the interferometric detection. We note that this result is only approximate, as it assumes low numerical aperture objective lenses. In most 4Pi-SMLM microscopes with an index mismatch between the immersion oil and the sample, this is not the case.

We account for sample shifts by treating each step of the multi-step scan as an independent measurement. For each step in the scan, the experimental PSF phase was determined separately during data analysis. The data for that step was then fit with a DS-PSF model with the specific phase for that step, and phase correction was applied as described. In this manner, the PSF phase at each step was determined precisely without using Eq. S1. Small shifts in registration between 3D volume images from different scan steps were resolved during drift correction.

#### Multicolor data analysis

Multicolor 4Pi-STORM data was recorded with additional filters in the detection path (DM3 in Supplementary Fig. S1) which act as a polarization-sensitive spectral filter. For each localization event, the ratio of S-polarized to P-polarized photons was determined, and this ratio was used for initial assignment of the localization as one of the two fluorophore species (Alexa Fluor 647 or Cy5.5) according to a simple threshold (see Supplementary Fig. S35).

The additional filter in the multicolor detection case causes each fluorophore to have a distinct detection PSF. In particular, with the filters in place, we found that the amplitudes of the S-polarized channels for the Alexa Fluor 647 PSF are attenuated by a factor of approximately 0.6 relative to the P-polarized channels. For Cy5.5, the S-polarized channels were attenuated by a factor of approximately 0.9. These values were measured directly from the data, by fitting a sample of localizations from each color species with a series of DS-PSF models with varying S/P amplitude ratio (see Supplementary Fig. S36).

After determining the correct PSF scaling for each fluorophore, final assignment of the fluorophore species for each localization was made by fitting with the DS-PSF model. For this purpose, the localizations were fit with each fluorophore-specific (rescaled) PSF, and the color was identified by selecting the best-fitting result (lowest chi-square value). This procedure minimizes crosstalk and ensures that the DS-PSF matches the true detection PSF for each localization, which is necessary to avoid introducing localization bias errors.

Using simulations, we investigated the effect of scaling the PSF model to have the correct amplitude ratio. Using a realistic PSF model (derived from a real PSF measurement) we generated simulated datasets for the two fluorophores, having the S/P amplitude ratio of 0.6 for Alexa Fluor 647 and 0.9 for Cy5.5. We then fit the data with either a correctly scaled PSF model or an incorrectly scaled model. Supplementary Fig. S37 shows that when the incorrect model is used to fit the data (e.g. data ratio 0.6, model ratio 0.9) a z-bias error is introduced into the estimated localization coordinate. This bias is *z*-position dependent, meaning that it would cause distortions on the order of 5-10 nm which vary over length scales of ∼100 – 200 nm, and cannot be simply corrected by an offset. Fitting the data with the correctly scaled PSF, results in zero bias error, however.

We validated the simulation results by analyzing experimental data which was fit with a correctly scaled or an incorrectly scaled DS-PSF model. A sample labeled with both fluorophores was analyzed by fitting all localization data with both versions of the PSF model. The fluorophore identity for each localization was determined using the S/P photon ratio, as above. For each set of fluorophore results, the difference in the estimated *z*-coordinate between the fit with the correct and incorrectly scaled PSF model is shown as a function of the *z*-coordinate (assuming the correct result is obtained with the correctly scaled model) in Supplementary Fig. S38. This experimental analysis reproduces a similar degree of bias error as seen in the simulations.

To explore the differences between the single color and multicolor optical configurations, we simulated Alexa Fluor 647 and Cy5.5 molecules with photon counts and background levels obtained from the experimental data (see Supplementary Table S1). The predicted localization precision vs. the emitter *z*-coordinate for each case is plotted in Supplementary Fig. S39. The results show that the localization precision for Alexa Fluor 647 is relatively unaffected by the configuration, with an average difference of 4% between the single color and multicolor modes. In the multicolor mode, the *z*-localization precision for Alexa Fluor 647 is modulated as a function of the *z*-coordinate (amplitude approx. 1 nm) due to the uneven distribution of fluorescence between the S- and P-channels. We note that this modulation would be avoided when using a different multicolor detection method, such as salvaged fluorescence^22^. Finally, comparing the performance of the two fluorophores used for multicolor imaging, the simulation shows that on average Cy5.5 exhibits a 10% higher localization uncertainty than Alexa Fluor 647, due to its intrinsically lower brightness.

Despite the relatively small difference in the emission spectra of Alexa Fluor 647 and Cy5.5, some degree of chromatic aberration may still be present, which could introduce an overall *z*-shift between the color channels. To correct for residual chromatic offset, a calibration sample was prepared in which the same structure (Nup107 in nuclear pore complexes) was labeled with both fluorophores, allowing the effect to be measured from the data. The *z*-offset between the fluorophores was determined to be approximately 10 nm, and subtracted from the data.

## Data availability statement

The data that support the findings of this study are available from the corresponding authors M.B. and S.W.H. upon request.

## Code availability statement

Full source code for the spline calculation and fitting routines are provided, under MIT license, at www.github.com/gpufit/Gpufit and www.github.com/gpufit/Gpuspline.

## Supporting information

Supplementary Video Descriptions

## Acknowledgements

We thank Tanja Koenen and Dr. Ellen Rothermel for support with cell culture and sample preparation. We thank Rainer Pick and the mechanical and electronics workshops of the Max Planck Institute for Biophysical Chemistry for design assistance and precision-machining of opto-mechanical components for the microscope. We thank Jaydev Jethwa for assistance with the microscope setup. We thank Matthias Kleinhans and Matthias Kulp for assistance with optical coatings. We thank Dr. Marco Roose for IT support. We thank Dr. Hazen Babcock (Harvard University) for helpful discussions concerning cubic spline modeling. The work has been supported by the Deutsche Forschungsgemeinschaft (DFG) (SFB1286/A07 to E.D.). M.B. gratefully acknowledges funding from the European Molecular Biology Organization (ALTF 800-2010), the Manfred Eigen Förderstiftung, and the Max Planck Society.

## Author contributions

M.B. and S.W.H. designed the project. M.B. and A.H. designed and constructed the 4Pi-STORM microscope. M.B. performed the experiments. M.B. and J.K.-F. developed algorithms and software for data analysis, and analyzed the data. A.P. implemented spline calculation algorithms, and fitting routines within the Gpufit framework. P.I. contributed to data visualization. T.S. and S.J. provided samples for the mitochondrial protein imaging and helped interpret the results. A.R.C.D. and E.D. provided samples for the neuronal cytoskeleton imaging and helped interpret the results. M.B., J.K.-F., S.J.S., and S.W.H. guided the project and wrote the manuscript. All authors discussed the data and commented on the final version of the manuscript.

## Competing interests statement

The authors declare no competing interests.

## Supplementary Notes

### 1 Bead step scan analysis

The bead step scan (Supplementary Fig. S18) featured 250 frames recorded at every scan position. In each recorded frame, a region of 7x7 pixels including the bead image was cut out and fit with the DS-PSF model as described above. In order to estimate the localization precision, the standard deviation of the *z-*positions estimated within a scan step were calculated from the squared differences in coordinates {*z*_i_} of consecutive estimations:

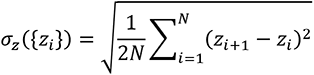

A localization was classified as localization artifact if the estimated *z-*position was more than 70 nm away from the expected *z-*position. Sample drift during the scan was corrected by subtracting a low-pass filtered smooth curve from the estimated *z-*positions over the recording time (for display only).

### 2 Cramér-Rao lower bound of the position estimation

We followed a standard approach, for example as outlined in Balzarotti et al.^1^ Because the fluorescence was detected by an EMCCD camera, we modeled the data as Poisson distributed with a variance that is approximately two times the expected number of photons, and a pixel readout noise term. The probability to obtain a measurement *d*(*x*, *y*) is then given by

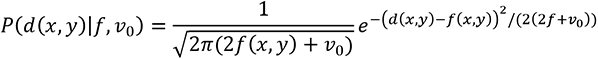

with the model *f*(*x*, *y*) defined as in Eq. 1, and the readout noise *v*_0_. The entries of the Fisher information matrix *I*(*θ*) for the fit parameters *θ* (amplitude, background, *xyz-*positions) were then computed as expectation values

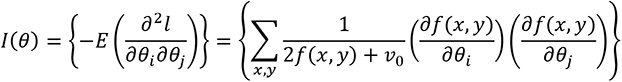

### 3 Astigmatism-based 4Pi-SMLM position estimation

Astigmatism-based 4Pi-SMLM data analysis was reported previously^2,3^, and this method was reproduced in our study for comparison purposes. We refer to the method as the ellipticity-phase analysis, because it is based on a measurement of both the ellipticity and the interference phase for each fluorophore image. For the ellipticity-phase analysis, the transformed images of the emitter in each channel were summed, and an elliptical Gaussian function was fit to the summed image, resulting in *x-y* coordinates of the localized event, and fitted widths *σ_x_* and *σ_y._* From these peak widths, a monotonic measure *m* was calculated as:

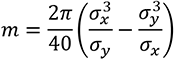

This measure is monotonic within ∼1.2 µm for the simulated astigmatic 4Pi PSF used in our study. A phase for the central Gaussian moment (φ_0_) was also computed, and the combination of *m* and φ_0_ allowed the unambiguous determination of the emitter’s *z-*position. Simplifying the analysis somewhat, a 4Pi PSF with the known correct PSF phase was used in the analysis of simulated data and the *z-*coordinate of the emitter was determined by finding the *z-*value which minimizes:

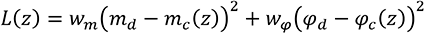

where *m_d_* is the monotonic measures of the data, *m*_c_(*z*) are the monotonic measures of the 4Pi PSF, φ_d_ is the zero order central moment of the data, φ_c_(*z*) are the zero order central moments of the 4Pi PSF, and *w*_m_, *w*_φ_ are empirically determined weighting factors (*w*_φ_ = 10*w*_m_).

### 4 Generation of simulated data

To simulate molecule event data for molecules at a certain position and with a certain brightness and background level, the mean expected signal within a region of camera pixels was computed by interpolating a PSF spline model with suitable center, amplitude and offset parameter values on a grid reflecting the camera pixel positions. From this model, noisy data was created by drawing from a Normal distribution with the model value as mean and a variance two times the model value plus camera readout noise term, independently for each pixel. The additional variance effectively simulates data from an EMCCD camera. The center positions were chosen so that the molecule location was uniformly distributed within the area of the peak camera pixel.

### 5 Symmetric and astigmatic 4Pi-STORM PSF calculation

We employed vectorial diffraction theory to compute the propagation of the electric field of a dipole emitter through our optical system^4–7^. We considered a dipole emitter located close to the common focus of two high NA objective lenses, whose collected signal is detected on a pixelated, planar detector. Aberrations arising due to mismatch between experimentally present and design values of refractive indices or thicknesses of the different layers (sample, glass, immersion oil) of the optical setup were modeled with a Gibson and Lanni phase term^5^. We do not assume rotational symmetry in the aperture, to allow for the introduction of an additional astigmatic phase distortion for the astigmatic PSF case. The foci of both objective lenses were overlaid in *z*, such that the points of maximal detection efficiency coincide. The thickness of the sample layer was assumed to be 20µm with the focus being positioned at 2µm inside the sample. The emitter position was then moved from 0 to 4µm inside the sample. We averaged over all emitter dipole positions by taking the mean intensity for an *x*, *y* and *z-*oriented emission dipole, effectively simulating a freely rotating fluorophore or a fluorescent bead. The calculated PSFs without astigmatism were compared to measurements of the experimental 4Pi PSF of the microscope setup, for each objective lens individually, as well as the interference of both. The modulation frequency and the axial and lateral FWHMs of the PSF envelope were found to match well with the simulation if the numerical aperture of both lenses was lowered to 1.3. The general asymmetric behavior of the 4Pi PSF with respect to the *z-*position that is visible in the experimental PSF was qualitatively be reproduced.

For the introduction of astigmatism an additional phase distortion proportional to 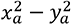 (aperture coordinates) was introduced after collection of the light by the objective lenses and before focusing by the tube lens. The strength of the astigmatism was adjusted so that the distance of the *z-*position of minimal FWHM in *x* and minimal FWHM in *y* matched a previously reported astigmatic 4Pi PSF^2^. This resulted in the phase apodization term 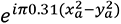. The corresponding symmetric 4Pi PSF without astigmatism simply did not feature such a phase apodization term.

### 6 Estimation of the localization artifact rate in Nup96 measurements

In the central 6x6 µm region of the measurement of a single cell nucleus (a section of which is shown in Fig. 2b) the nuclear envelope is slightly curved and can be accurately estimated as a surface from the Nup96 data alone. For this, a distance weighted average *z-*position of all localizations within the central region was computed. In a second iteration, localizations far away from this surface were disregarded. The Gaussian weighting factor acted as a smoothing parameter for the computed surface. The *z-*position of the nuclear envelope estimation was then subtracted from each localization’s *z-*position, to place all the data at a common reference point in the *z-*direction. The reference point was further shifted so that the histogram part referring to artifact free localizations was located at *z*=0 on average.

Non-specific labelling was visible everywhere in the sample. To control for it, a mask in the *x*-*y* plane was created that contained all nuclear pores. The nuclear pore mask included ∼20% of the area and ∼95% of the available localizations. Histograms of all surface corrected *z-*positions within that mask and outside of the mask were calculated. The histogram from outside of the mask was rescaled to account for difference in area and subtracted from the histogram of *z-*positions within the mask. The resulting histograms are shown in Figs. 2c and 2d.

The localization artifact rate was computed by comparing the number of surface corrected localizations within a *z*=±0.15 µm range to that in a *z* = ±0.45 µm range with the localizations in the 0.15-0.45µm range being the localizations in the next 4Pi PSF fringe, i.e. the most likely mis-localizations. The surface and nuclear pore mask detection for the PSF phase with the lowest artifact rate was used for all tested PSF phases.

### 7 Unwrapping of Beta-II Spectrin and Mic60 data

To create the unwrapped views of the beta-II spectrin distributions along an axon, and of the Mic60 distributions in mitochondria, we modeled the distributions as tubular structures that locally are represented by a rotated, elliptical shell. First, we iteratively refined our model of the center line, i.e. the smooth line that passes along the center of an axon or a section of the mitochondrion, together with the model of the elliptical shape of the membrane at each position of the center line. Initially, an approximation of the centerline was chosen manually. The centerline was then smoothed and divided into sections of equal distance (40 nm). Next, a Cartesian coordinate system (reference system) was chosen from a tangential vector to the centerline and two perpendicular vectors in the transversal plane. The last two vectors were chosen so that initially one pointed upwards, i.e. in the positive *z-*direction, and that the change in vectors between consecutive centerline points was as small as possible. That allowed smoothly sliding the reference system along the centerline. At each centerline position the volume rendered data was linearly interpolated on a rectangular grid in the associated coordinate system and a slice of certain thickness (100-200nm) along the centerline direction was added up. To this an elliptical shell of a certain thickness (typically 80-100nm thick) was fit to the image of the data in the reference system (by maximizing the normalized cross-correlation) and a shift in the center in *x*, *y* as well as the lengths of the major and minor axes and the orientation angle of the ellipse were obtained. This was done for each position of the centerline independently, and the new centers as well as major and minor axis lengths and ellipse orientations were smoothed along the centerline length coordinate. This ellipse fit was repeated until the process converged, i.e. the centerline did not change further, and the fitted ellipse sizes were stable (typically ∼10 iterations were performed).

In a final step, unwrapped views were created by progressing along the centerline in small steps (4 nm pixel size) and calculating a grid of an elliptical shell with a certain diameter (80-100nm) in the transverse plane using the ellipse parameters determined in the previous step. The grid had a radial and an azimuthal direction where the azimuthal positions were equally spaced on the circumference of the fitted ellipse that represents the center of the elliptic shell. An equal spacing of positions along the circumferential coordinate is achieved using the incomplete elliptic integral of the second kind. The data was again linearly interpolated on this grid (see Supplementary Video S4) and added up along the radial direction, resulting in a summed up signal of the beta-II spectrin or Mic60 around the circumference of the axon or mitochondrion. The aspect ratio of the unwrapped views was chosen so that the mean circumferential coordinate and the length along the centerline are scaled equally.

### 8 Gpufit and Gpuspline software libraries

Two open source software libraries were written to calculate spline coefficients and interpolated spline values, and to perform nonlinear least-squares curve fitting with the spline model functions.

Gpuspline (www.github.com/gpufit/Gpuspline) was used to create a cubic spline representation of the 4Pi PSF from pixelated bead scan data. The API function *calculate_coefficients_3d* calculates a set of coefficients for a single channel 3D pixel stack representing the x,y,z intensity distribution of the PSF in one channel. We called this function four times, once for each image channel, to obtain the spline coefficients describing the four-channel 4Pi PSF. The order of the spline coefficients representing a dataset and the corresponding Matlab or Python binding calls are described in the Gpuspline documentation (www.gpuspline.readthedocs.io/en/latest/index.html). Examples of the calculation of spline coefficients from a dataset, as well as interpolation of a dataset using the calculated spline coefficients, are included with the project source code.

Gpufit (www.github.com/gpufit/Gpufit) was used to fit the experimental data, using the cubic spline models generated with Gpuspline. For this purpose, we created new fit model functions for general purpose multichannel, multidimensional spline fitting. The arrays of spline coefficients for each image channel were concatenated, and served as the definition of the multichannel spline models in Gpufit. The fit model function *SPLINE_3D_MULTICHANNEL* was used for 4Pi STORM data analysis, and the concatenated spline coefficients were passed in using the *user_info* parameter of the *gpufit* API function call. In addition to fitting with multiple image channels, the dynamic phase of the 4Pi PSF (from Eq. 6 in the main text) was implemented as a model parameter in another Gpufit model function *SPLINE_3D_PHASE_MULTICHANNEL,* which requires three sets of multichannel spline coefficients passed via the *user_info* parameter, corresponding to the PSF envelope *h*_env_, the modulation *h*_mod_, and the shifted modulation *h*^90^ as defined in Eqs. 3-5. The definitions of the spline fit model functions, the structure of additional user information for the fit models and the corresponding Matlab or Python binding calls are described in the Gpufit documentation (www.gpufit.readthedocs.io/en/latest/index.html). Examples for the use of a spline representation of a model function to fit data with Gpufit are included in the project source code (www.github.com/gpufit/Gpufit/tree/master/Gpufit/matlab/examples).

### 9 Supplementary analysis software and data

An additional software package is provided with the manuscript in order to demonstrate 4Pi-STORM data analysis using the DS-PSF model. This package is written in Matlab, and contains experimental data including a fluorescent bead scan (experimental PSF), and the emitter images for the Nup96 dataset shown in Fig. 2. The package executes each step in the DS-PSF analysis, including the creation of the spline representation of the PSF (using Gpuspline), estimation of the phase evolution of the PSF and fitting the localization events with the optimal PSF phase (using Gpufit), correction for phase drift, and correction for sample drift.

The supplementary software and data are included with the supplementary materials of the manuscript, as a zip file: *supplementary_analysis_software.zip*. Usage instructions are contained in the file *README.txt*. This package includes Windows binary files for the Gpufit and Gpuspline libraries. Note that a CUDA-supported graphics processing unit (GPU) is required to run the software, as the fits are executed in parallel on the GPU.

### 10 Supplementary mechanical drawings

Digital mechanical designs are provided as supplementary data with the manuscript. Full 3D mechanical drawings for the fixed objective mount, movable objective mount, and sample stage (Supplementary Figs. S6, S7, and S8) are saved in STEP format, a widely used format which is accessible from many CAD software environments. The drawings are contained in the zip file: *supplementary_mechanical_drawings.zip*. All mechanical designs are released under the terms of the CERN Open Hardware License Version 2 (Strongly Reciprocal).

---

**Figure S1:**
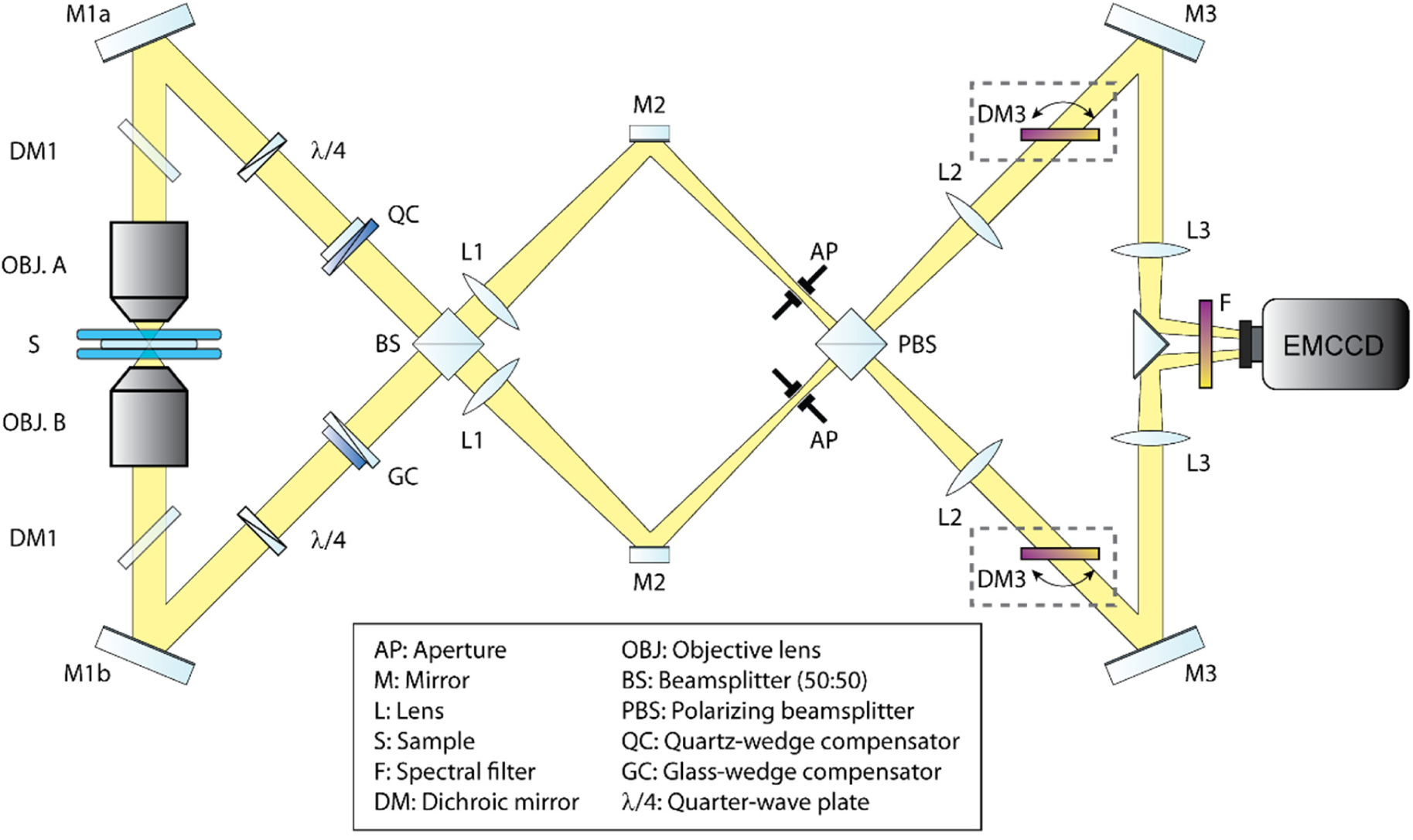
4Pi-STORM detection path. Optical layout of the 4Pi-STORM interferometric cavity and fluorescence detection path. Fluorescence emitted by the sample (S) is collected by the two objective lenses (OBJ. A and B), and travels along the arms of the cavity passing through the dichroic mirrors (DM1), quarter wave plates (λ/4), and Babinet-Soleil compensators (QC and GC). The fluorescence photons meet and self-interfere at the 50:50 beam-splitter cube (BS). Tube lenses (L1) form an image of the sample at the apertures (AP). After passing through the polarizing beam-splitter cube (PBS), the S-polarized fluorescence is separated from the P-polarized fluorescence. The telescopes formed by lenses L2 and L3 create four images of the sample on four quadrants of the EMCCD camera. The fluorescence signal is filtered by spectral filters placed before the camera (F). In the multicolor detection configuration, dichroic mirrors DM3 are placed in the detection path after lenses L2. These dichroic mirrors act as polarization sensitive long-pass spectral filters, and are mounted on rotation mounts so that their angle can be precisely adjusted to set the cutoff wavelength of the filter.

**Figure S2:**
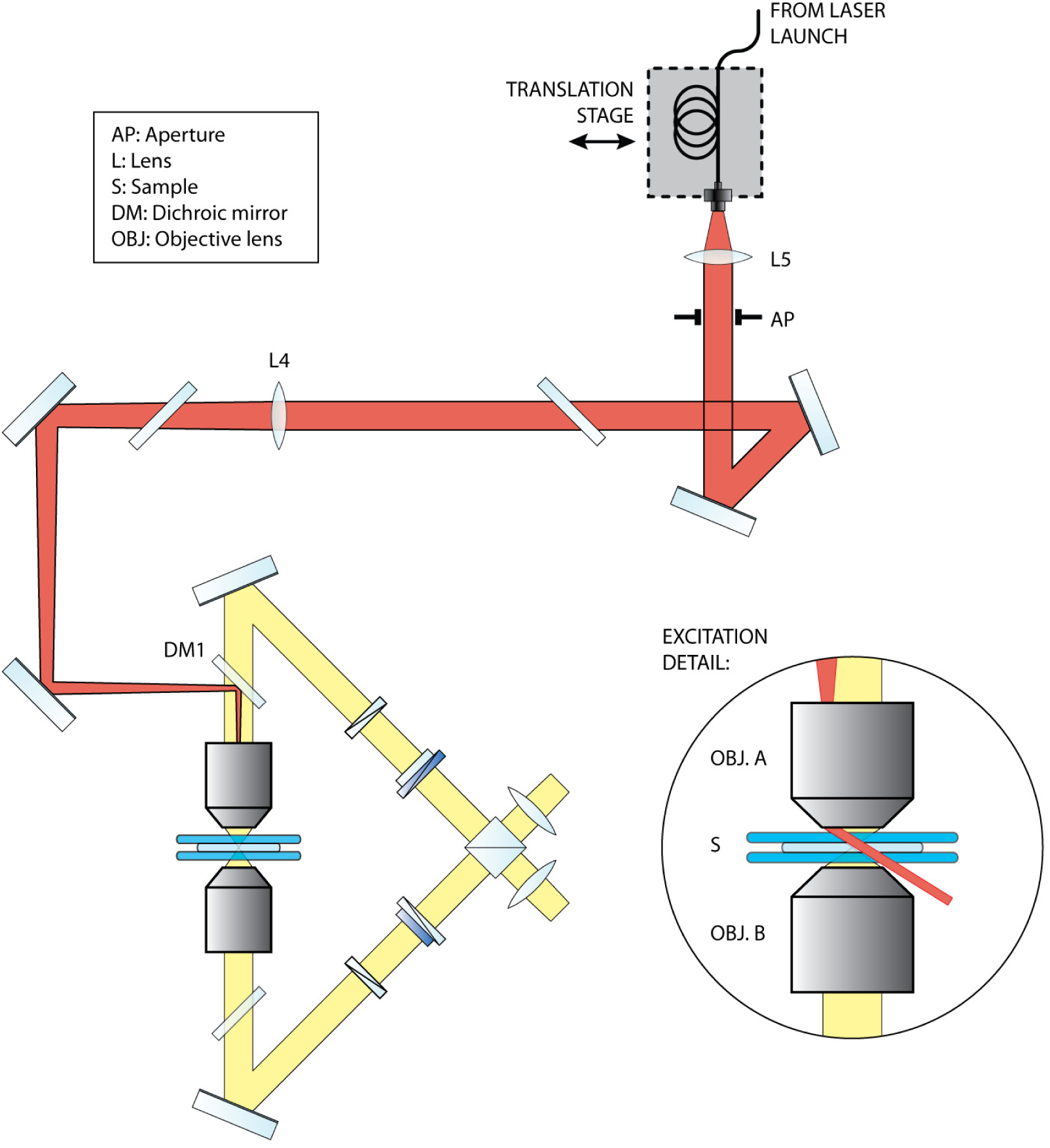
4Pi-STORM excitation path. Optical layout for the sample illumination path. Laser light is coupled into a high-power, polarization-maintaining single mode optical fiber, and the output of the fiber is mounted on a translation stage. Light from the fiber is collimated by lens L5. Lens L4 focuses the beam onto the back focal plane of the objective lens (OBJ A). The excitation beam is coupled into the 4Pi cavity at the dichroic mirror DM1. The adjustable aperture (AP) is located in a plane which is conjugate to the sample plane, and may be adjusted to limit the extent of the illumination area. The output of the fiber is located in a plane which is conjugate to the back focal plane of the objective lens, and by translating the fiber output laterally, the angle of the excitation beam at the sample may be adjusted between epi-illumination and near-TIRF illumination modes (see excitation detail, inset).

**Figure S3:**
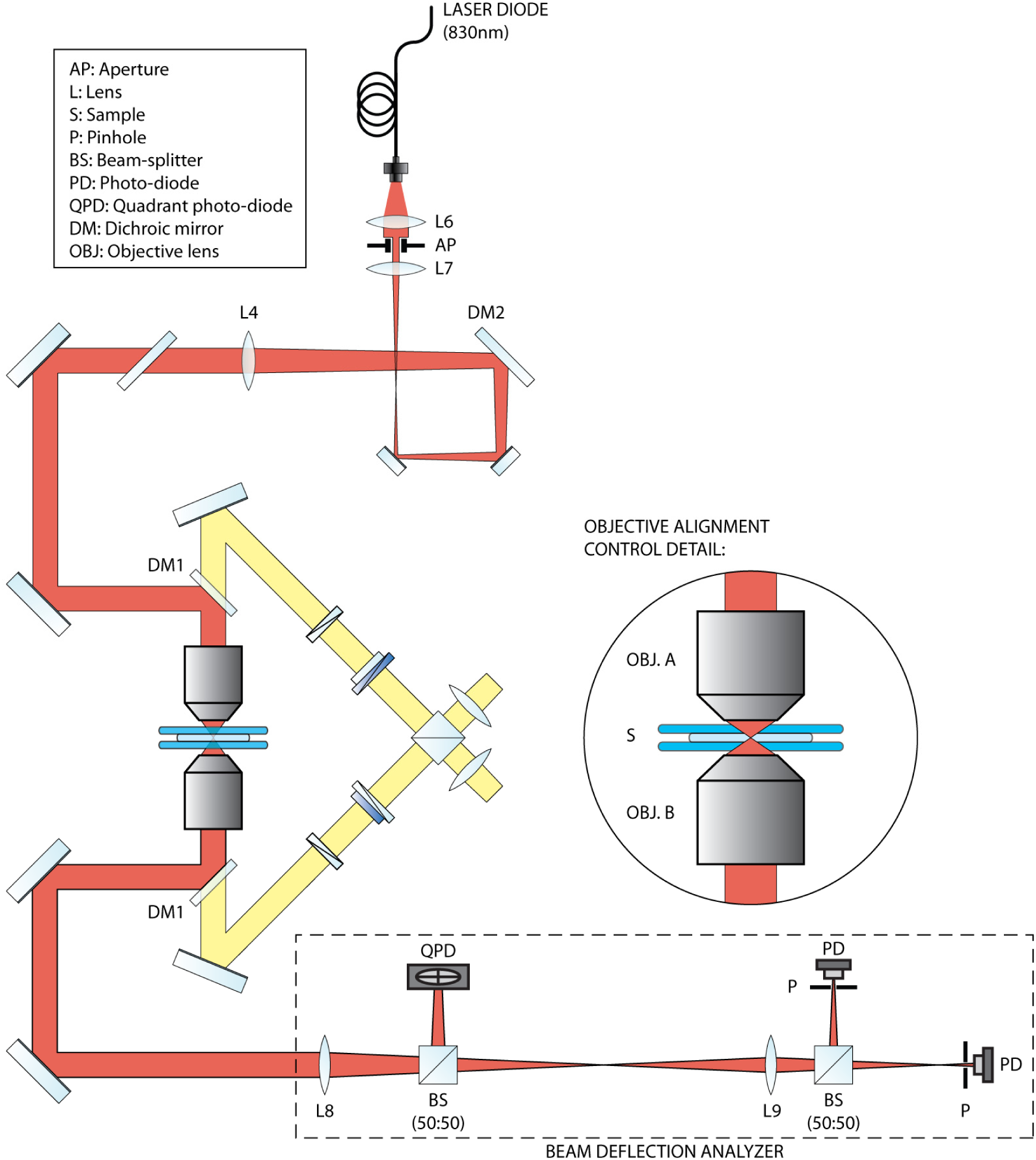
Objective alignment control system. A fiber-pigtailed laser diode at 830nm is collimated by lens L6, and expanded by lenses L7 and L4. The beam fills the back aperture of Objective A. The beam is focused to a point in the sample, and then re-collimated by Objective B (see detail, inset). Any lateral shift between the two objectives is detected as a shift in the angle of the outgoing beam. Any shift in the focus position of the two objectives is detected as a change in the collimation of the outgoing beam. The angle and collimation of the beam, after Objective B, are monitored using the deflection analyzer, consisting of a quadrant photodiode, and two additional photodiodes placed behind pinholes.

**Figure S4:**
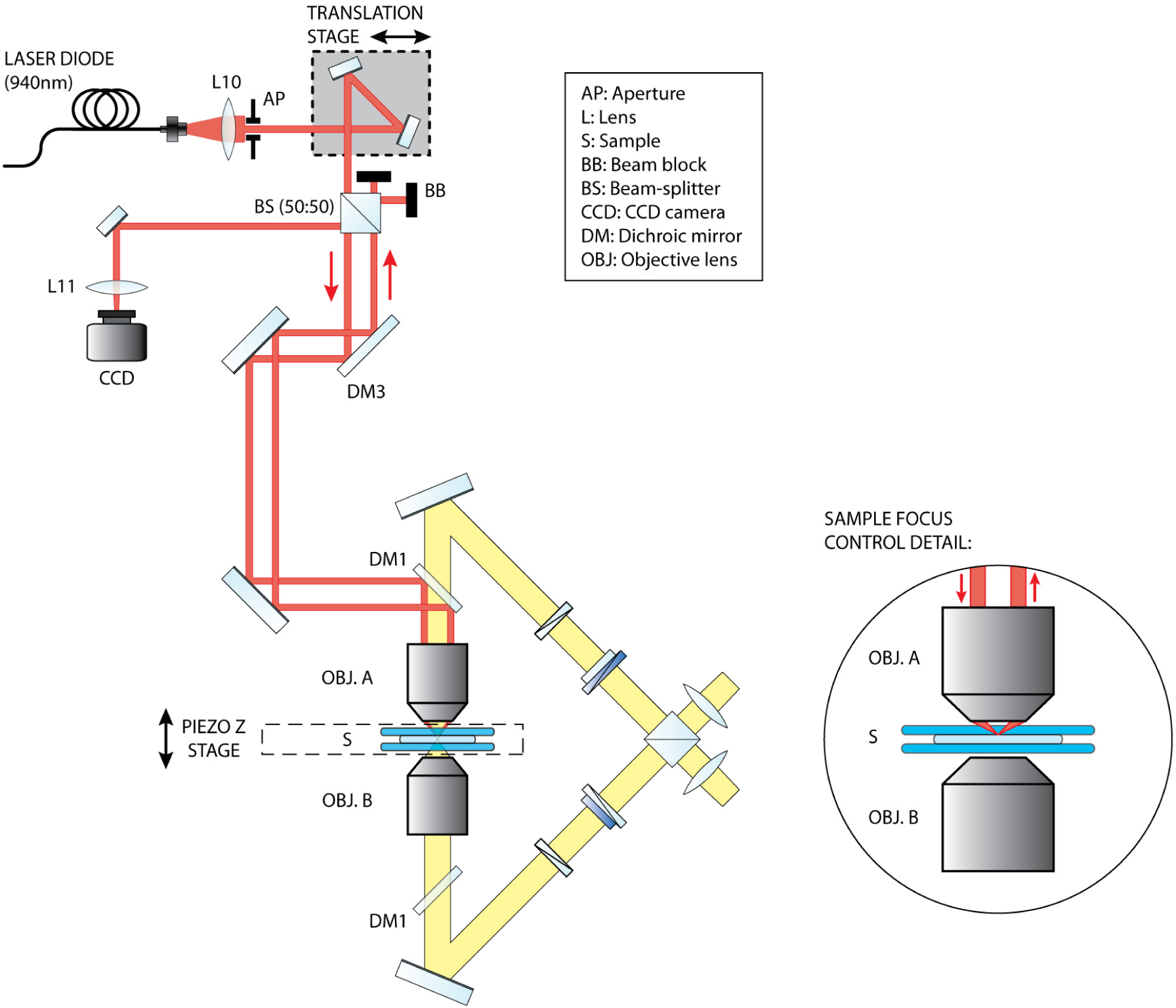
Sample focus control. The position of the sample with respect to Objective lens A is maintained by a feedback system, which is based on an infrared laser beam which reflects from the coverglass-sample interface and is detected on a CCD camera. Light from a fiber-pigtailed laser diode (940nm) is collimated, and directed to the edge of the back aperture of Objective A. A translation stage facilitates adjustment of the beam entrance position. Light emerges from the objective lens at a sharp angle (see inset), is reflected at the glass-water interface in the sample, and is re-collimated by the objective lens, exiting on the other side of the back aperture. The returning light beam is reflected at the beam splitter and focused onto a CCD camera. Any shift in the sample position along the optical axis, relative to the objective lens, is detected as a shift in the beam position on the CCD. By monitoring the beam position, sample drift is continuously corrected by adjusting the piezoelectric transducer (focus piezo) built into the sample translation stage.

**Figure S5:**
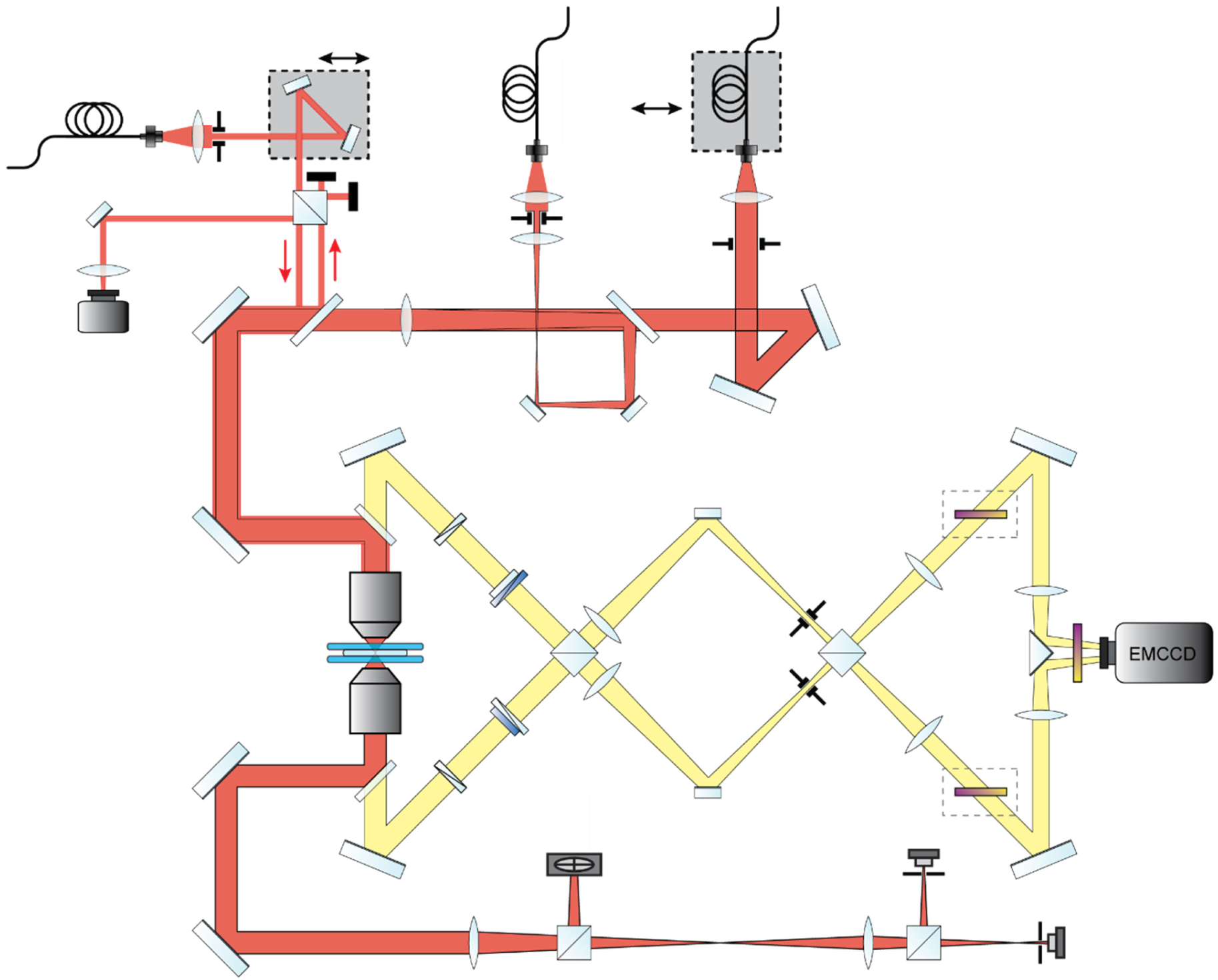
4Pi-STORM optical system overview. The 4Pi fluorescence detection cavity, excitation light sources, objective alignment control system, and sample focus control system, together form the 4Pi-STORM microscope. This figure shows how these systems, shown in detail in Supplementary Figs. S1, S2, S3, and S4, are integrated together on a single optical table. The system was constructed in the horizontal plane, on an optical table measuring 150cm x 120cm. The laser launch was built on a separate optical breadboard (not shown).

**Figure S6:**
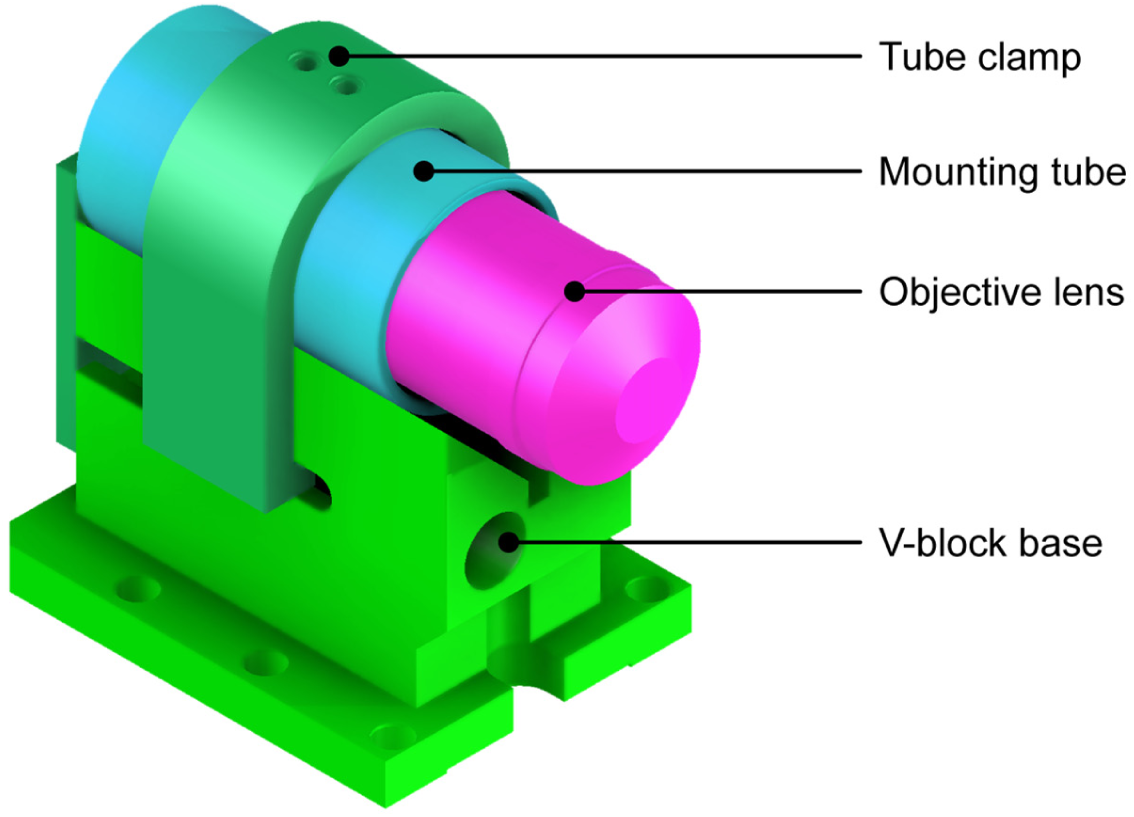
Fixed objective mount. Mechanical assembly drawing of the mounting system for the fixed objective lens (Objective A). The objective lens is mounted in a stainless steel threaded tube, which is supported by a machined aluminum V-block. The tube is clamped to its support using a U-clamp, tightened by two set screws at the top of the clamp. A mechanical stop at one end of the V-block sets the axial position of the mounting tube. This design allows the objective lens to be removed from the system, and replaced, with high positional reproducibility.

**Figure S7:**
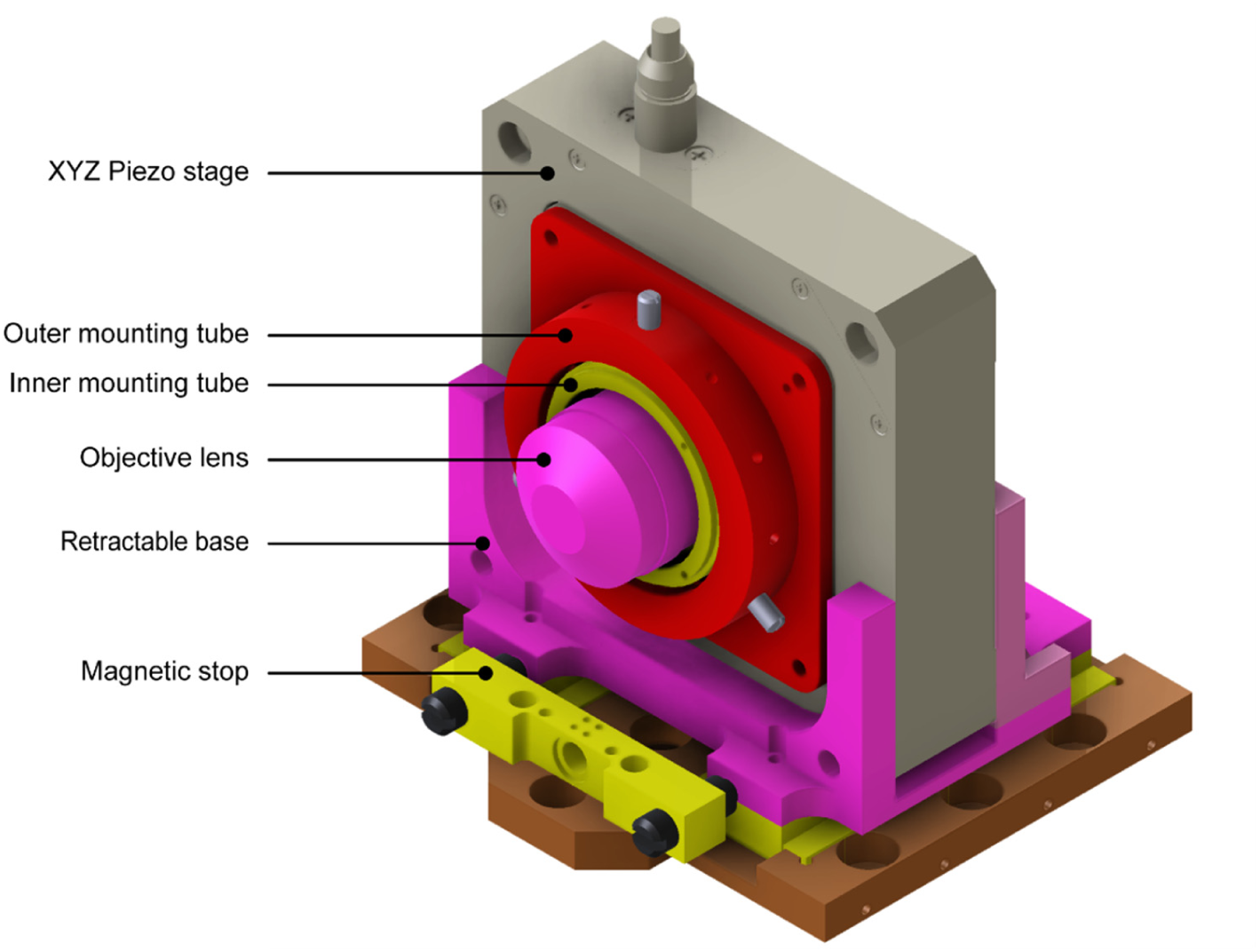
Movable objective mount. The position and orientation of the second objective lens (Objective B) is fully adjustable. The objective is mounting in stainless steel threaded tube, which itself is held within an outer mounting tube (colored red in the drawing) by six set screws. Adjustment of these screws allows coarse positioning of the objective, and adjustment of tip and tilt, during initial microscope alignment. The mounting assembly is held within the aperture of a three-axis piezoelectric stage, which provides fine translation in *x*, *y*, and *z*. The piezo stage is supported in a sliding mount, which can be translated by several centimeters for sample insertion and removal. A magnetic stop at one end of the translation range provides coarse focus adjustment for the movable objective.

**Figure S8:**
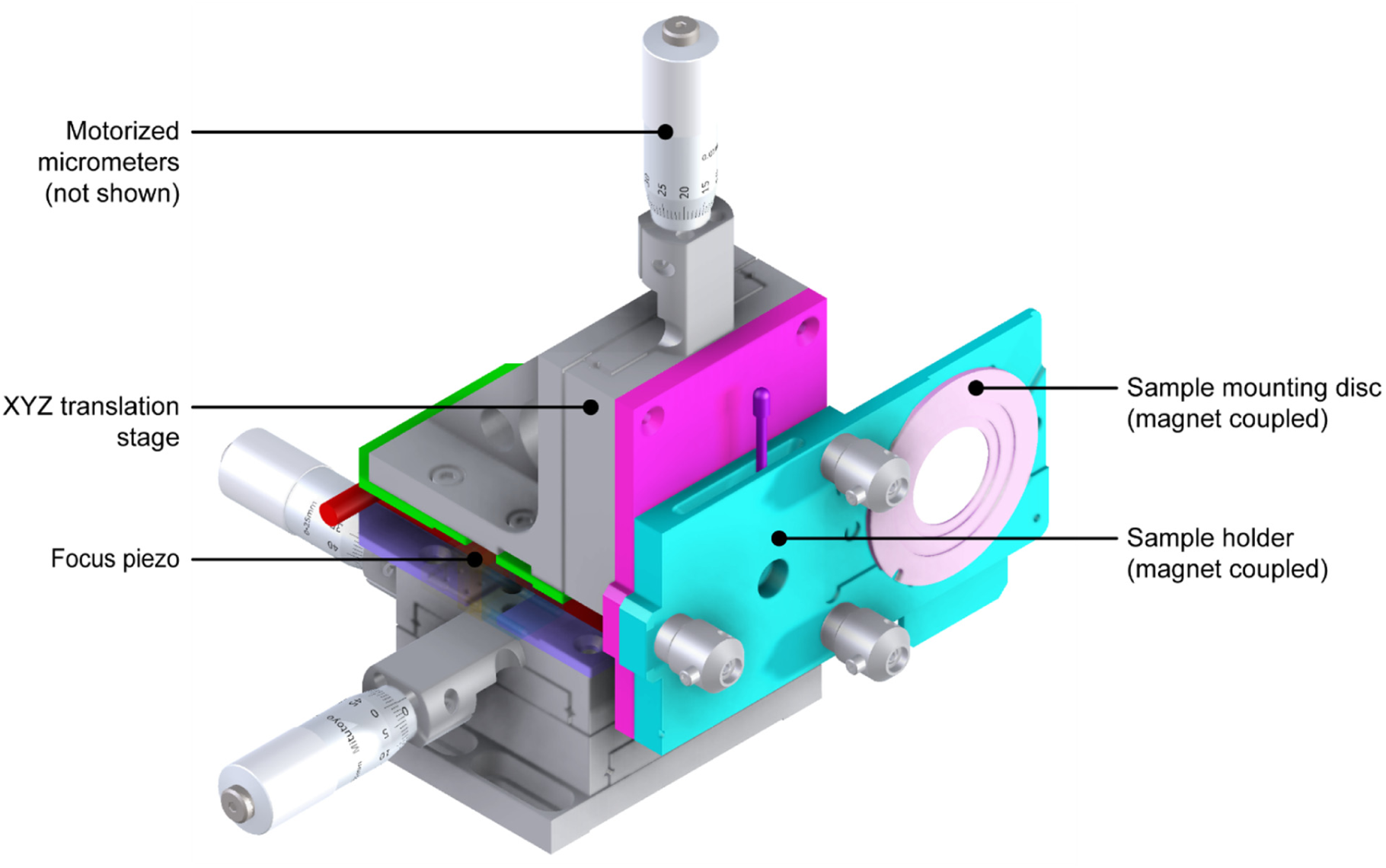
Sample mounting stage. The sample stage consists of a three-axis coarse translation stage, a linear piezoelectric stage, and a removable sample holder. Two axes of the XYZ translation stage are driven by motorized micrometer screws (not shown) to provide lateral sample translation. A manual micrometer screw provides coarse focus control, and a linear piezoelectric transducer provides fine focus control over a range of 30 micrometers. The sample coverslip is mounted on a stainless steel disc, which is magnetically coupled to the sample holder. The disc is oriented vertically, orthogonal to the objective lenses. The sample holder (colored cyan in the drawing) is also magnetically coupled to the translation stage, and may be released by shifting a lever. While coupled together, the sample holder and mounting disc may be inserted or removed from the microscope in order to change the sample.

**Figure S9:**
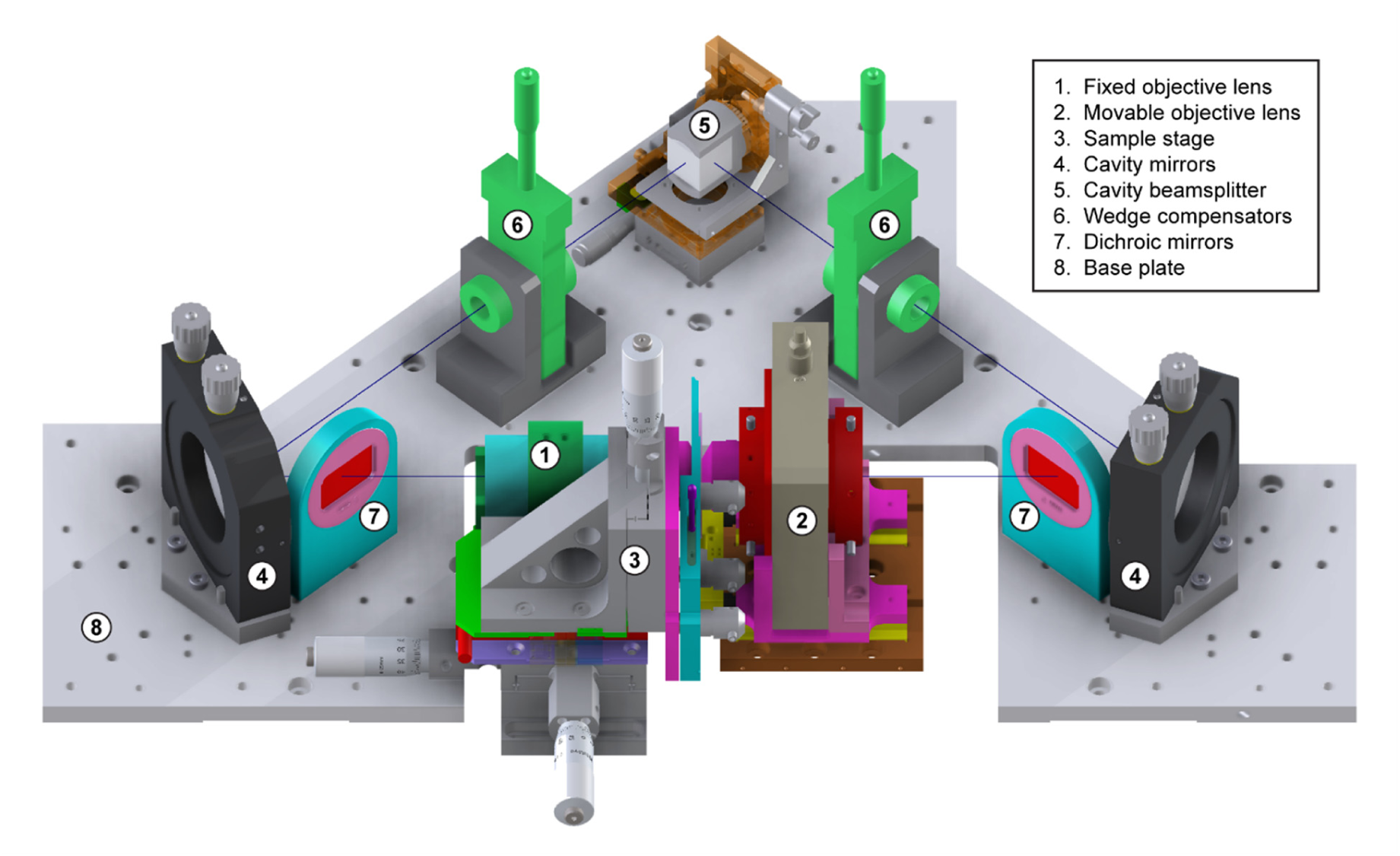
4Pi-STORM interferometric cavity overview. Mechanical assembly drawing showing the 4Pi-STORM interferometric cavity. A machined base plate is used to assist in precise positioning of the cavity mirrors, dichroic mirrors, compensators, and beam splitter. The scaled drawing shows the positioning of the objective lenses and sample stage relative to the other components. Note that the quarter-wave plates are not shown in the drawing.

**Figure S10:**
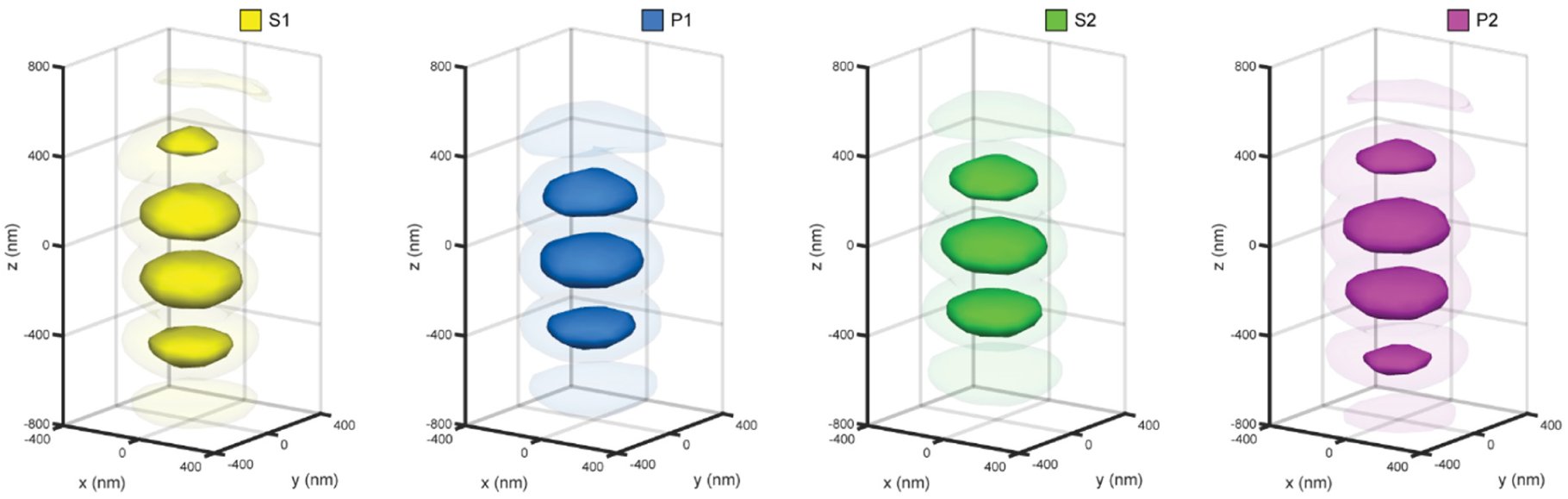
Individual channels of the 4Pi PSF. The four channels of the experimental 4Pi PSF shown in Fig. 1b and 1c, rendered as 3D isosurfaces at relative threshold levels of 35% and 13%, respectively.

**Figure S11:**
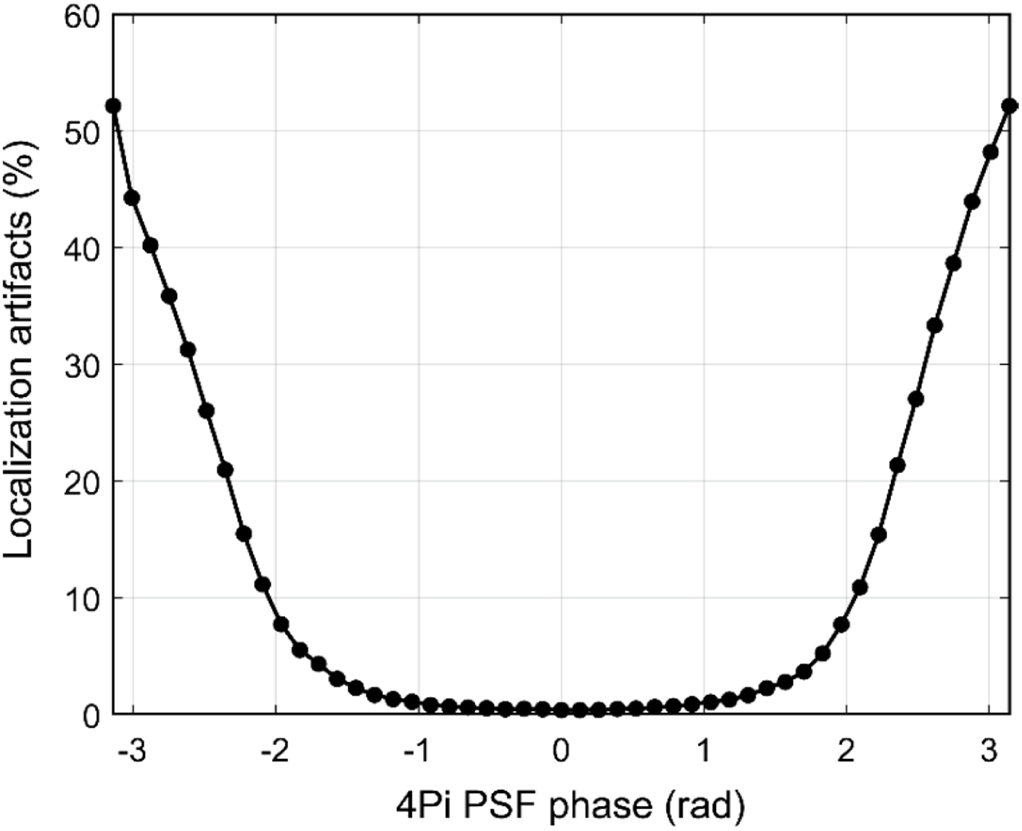
Localization artifact frequency vs. PSF phase error (simulation). A spline model of the 4Pi PSF was created from a fluorescent bead scan. Simulated fluorophore images were generated based on this model, with *z* coordinates evenly spaced over a range of 1 µm around the focal plane, and with a mean brightness of 8000 photons. The phase of the PSF model was then shifted (see Methods) and the model was used to fit the simulated fluorophore data for various PSF phase shifts. The fraction of localization artifacts was determined by generating a histogram of the difference of fitted and simulated z positions and marking all but the largest peak in the histogram as artifacts. This estimation is equivalent to the localization artifact estimation for experimental data. These results were also validated using an experimental dataset (Supplementary Fig. S12).

**Figure S12:**
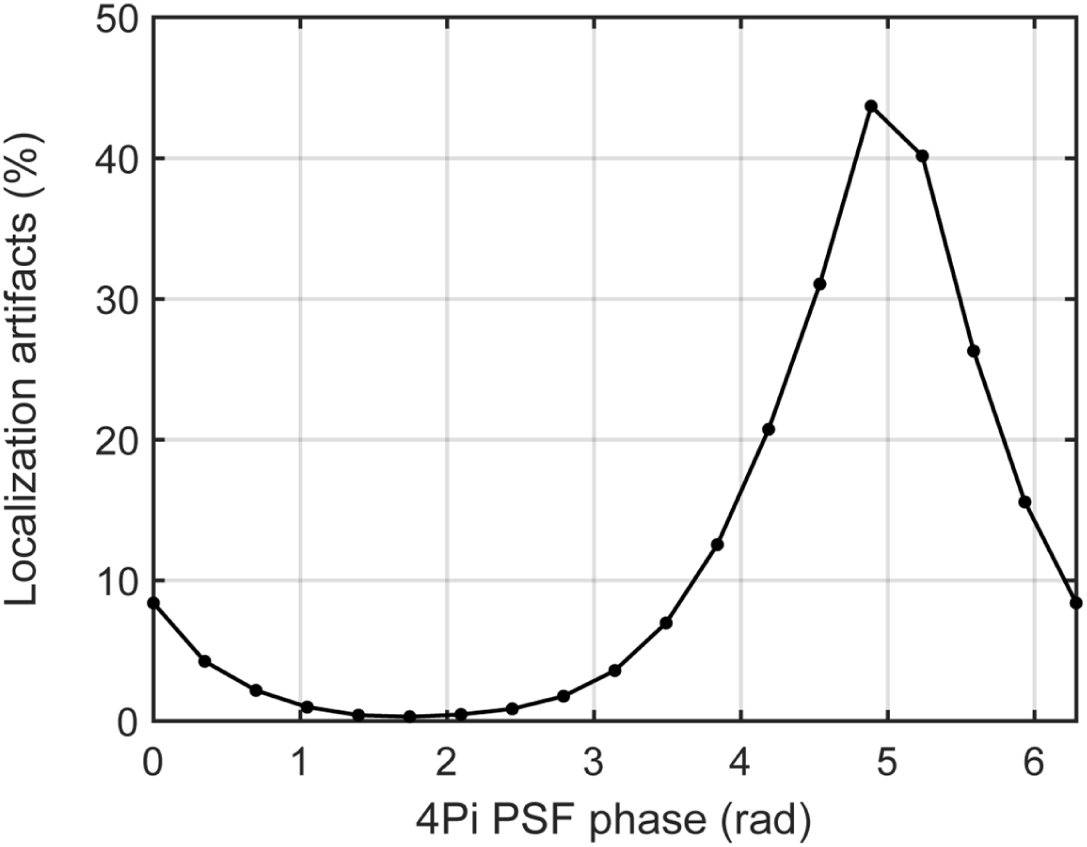
Localization artifact frequency vs. PSF phase error (experiment). Localization artifact frequency for an experimental dataset (Nup96 shown in Fig. 2) versus the phase of the cubic spline PSF model used to analyze the dataset. The PSF phase was numerically shifted over 18 steps, the dataset was analyzed with the PSF model calculated at each step, and the fraction of localization artifacts was determined from the localization results (black circles). The plot shows that the artifact rate is lowest (close to zero) at the global optimum PSF phase for the measurement, and remains low over a range of at least 2 radians centered at the best phase. At the opposite phase, the artifact rate approaches a maximum value of 50%.

**Figure S13:**
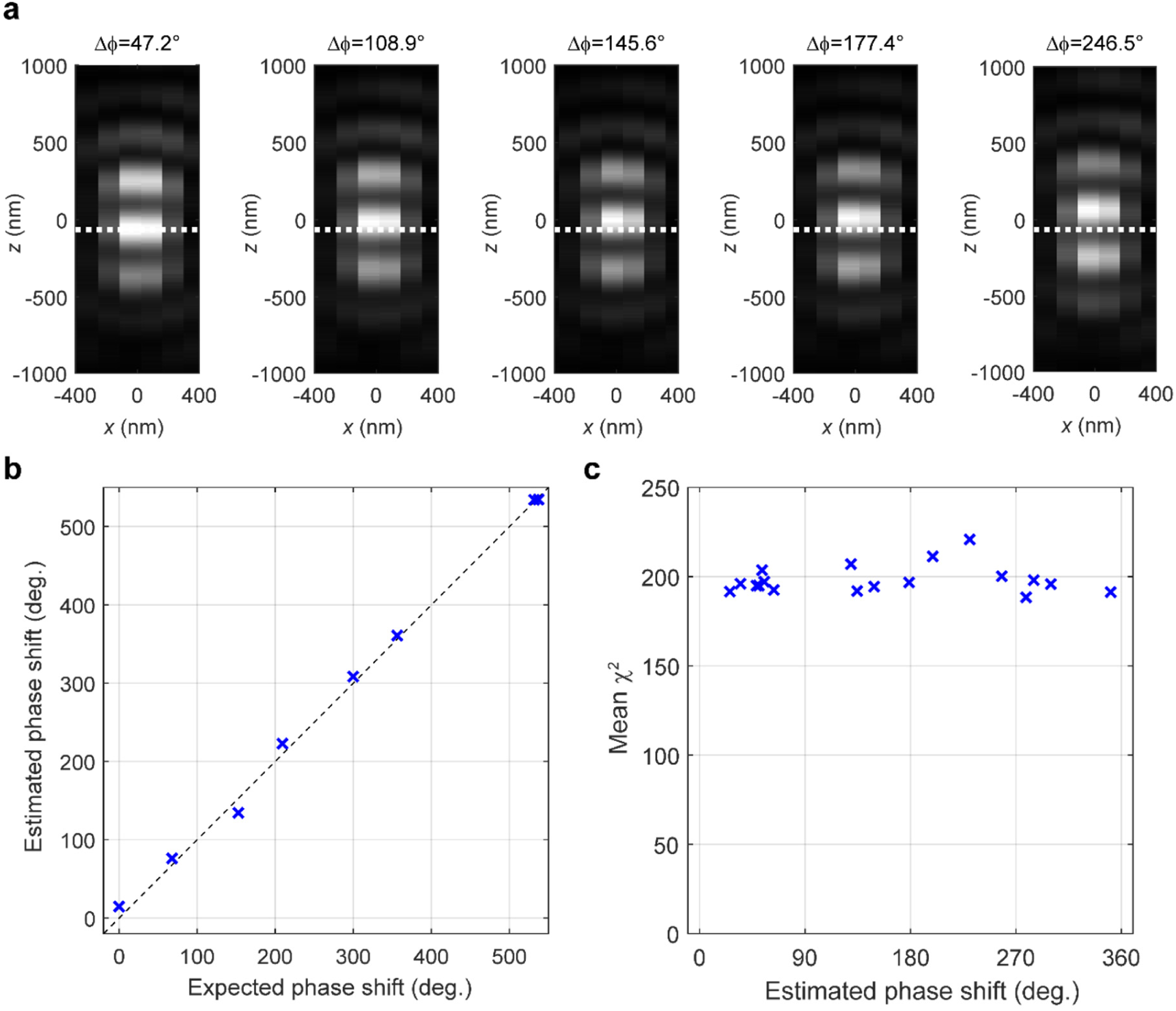
Validation of 4Pi PSF phase shift algorithm. A series of externally applied phase shift were introduced into the 4Pi cavity by manually changing the thickness of the BK7 glass wedge compensator in one arm of the cavity. This control experiment was used to verify the accuracy of the PSF phase estimation, and also the accuracy of the algorithm used to numerically shift the PSF phase. After each manual shift, a bead scan was recorded to measure the PSF, and the phase of the PSF was estimated using the spline fit approach. **(a)** Cross-section views (*x-z*) of the bead scans after each phase shift. The estimated phase shift is shown above the bead image. **(b)** Estimated phase (blue markers) at each shift position versus the expected phase (in degrees). A dashed line with a slope of 1 is plotted for reference. The results show that the estimated phase agrees well with the expected phase shift, within the precision of the micrometer screw on the compensator. The expected phase shift was calculated from the known relationship between the optical path length of the cavity and the glass wedge thickness. **(c)** For each phase estimation, the fit procedure returns a mean value for chi-square (<χ^2^>), which measures of how well the PSF model describes the data. A plot of <χ^2^> vs. the estimated phase showed no dependence on the amount of phase shift, demonstrating that the numerical phase shift algorithm yields physically accurate estimates of the true PSF.

**Figure S14:**
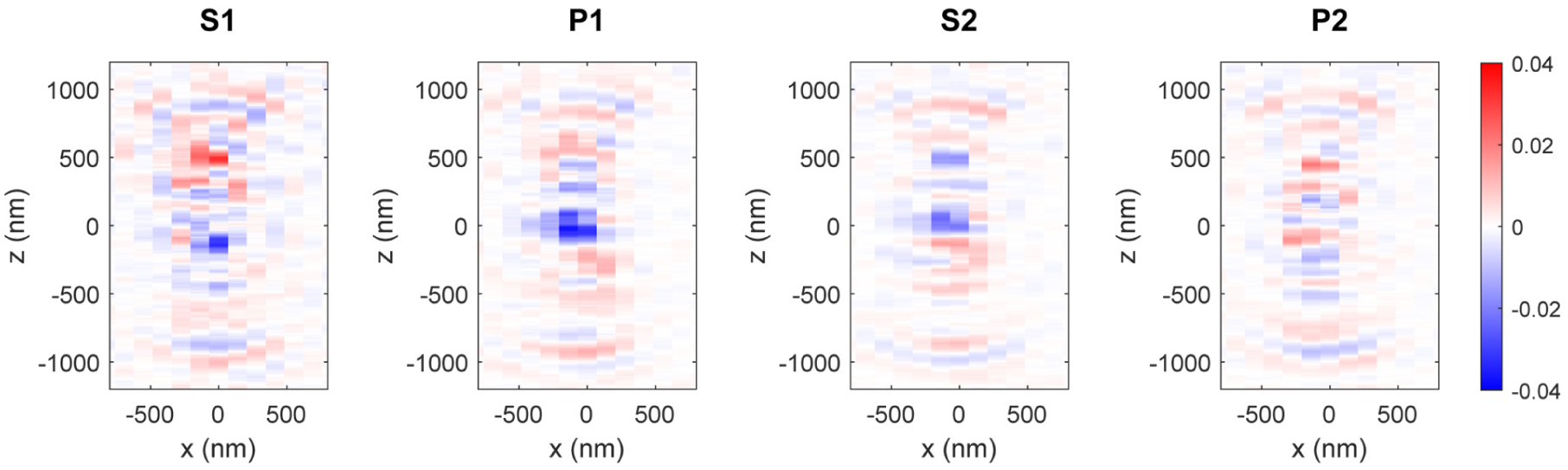
Comparison between PSF spline and measured bead scan. Normalized difference between the cubic spline PSF model obtained from one bead scan and the image stack from a second bead scan, for all detection channels. To match the second bead scan, the cubic spline PSF model was translated in *x*, *y*, and *z*, and its phase was shifted by 26°. An *x-z* cross-section through the center of the difference stack is shown for each detection channel. The maximum absolute deviation between the spline and the bead scan was 4% of the maximum intensity of the PSF.

**Figure S15:**
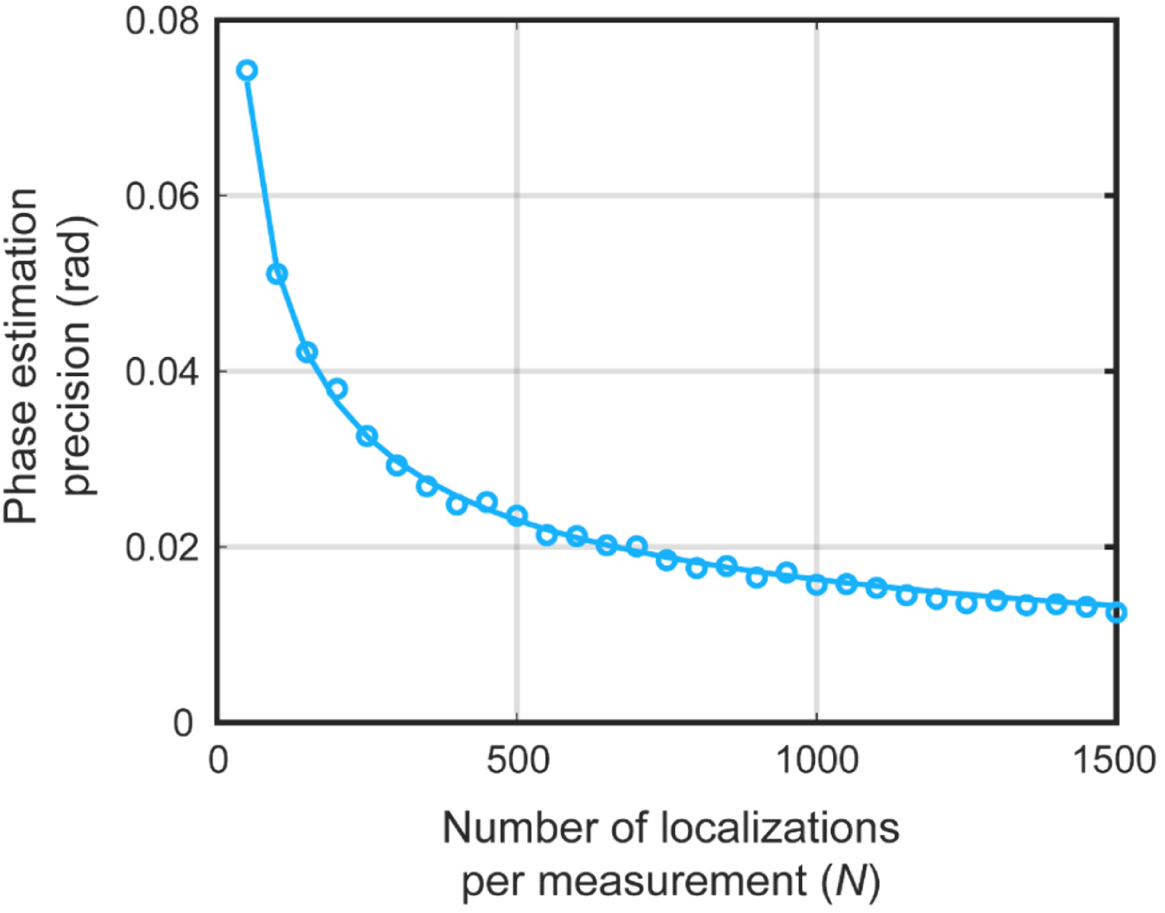
Precision of phase estimation. Precision of the PSF phase estimation as a function of the number of localizations used to estimate the phase. Simulated localization data was generated based on a PSF with a known phase shift, and then phase was then estimated using different numbers of input images. The simulated data featured emitter *z-*positions uniformly distributed over a 1 µm range around the focal plane, an emitter brightness of 8000 photons per event and a background signal of 10 photons per pixel. The simulation results (blue circles) show an improving precision as the number of localizations per measurement increases, as expected. The data were fit with a power law curve which scales as the square root of the number of input images (blue line), with a brightness- and background-dependent amplitude factor.

**Figure S16:**
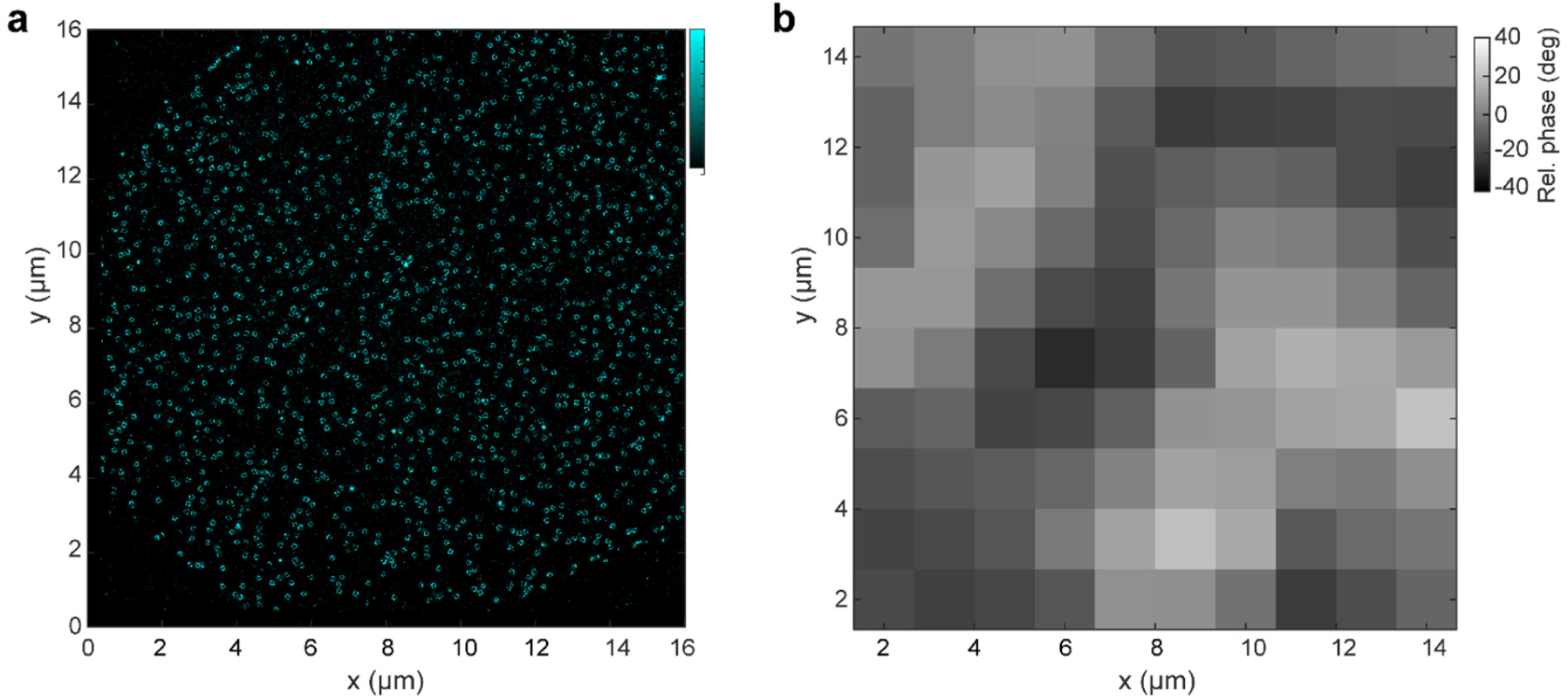
PSF phase variation across the field of view. Phase variation across the field of view of the 4Pi-STORM microscope. The relative phase was determined by analyzing subsets of localizations from an image of nuclear pore complexes (Nup107) in a nuclear membrane which extends across a large area (a). The localization data was subdivided into bins from sample regions 1.33 x 1.33µm in size, and the PSF phase was determined for each bin (b). Within the image, the phase was found to vary over a range of ± 40 degrees (b).

**Figure S17:**
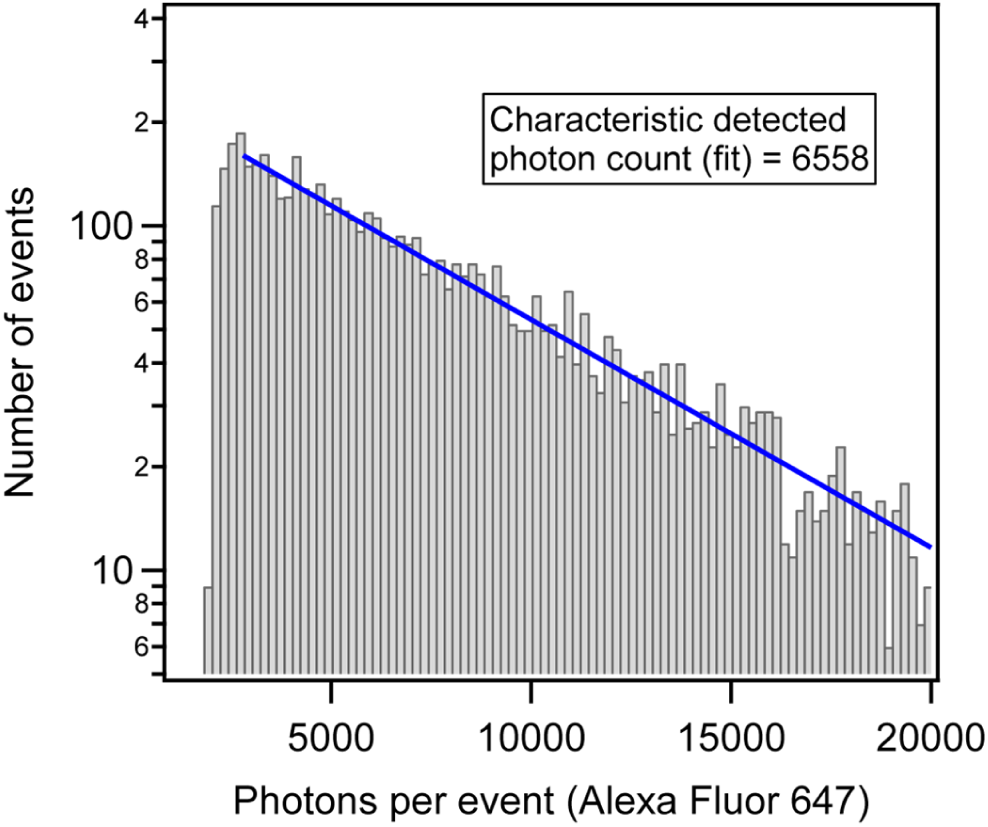
Detected photons per event, Alexa Fluor 647. Single molecules of Alexa Fluor 647-labeled double-stranded DNA, bound to a glass coverslip, were used to measure the localization precision of the 4Pi-STORM microscope. A histogram of the number of photons per on-off switching event (grey bars) was fit with an exponential function (blue line) to determine the characteristic detected photon count. Events with fewer than 1500 photons were discarded from the analysis. The fit yields a characteristic photon count of 6558, and the mean value of the photon distribution was 8002.

**Figure S18:**
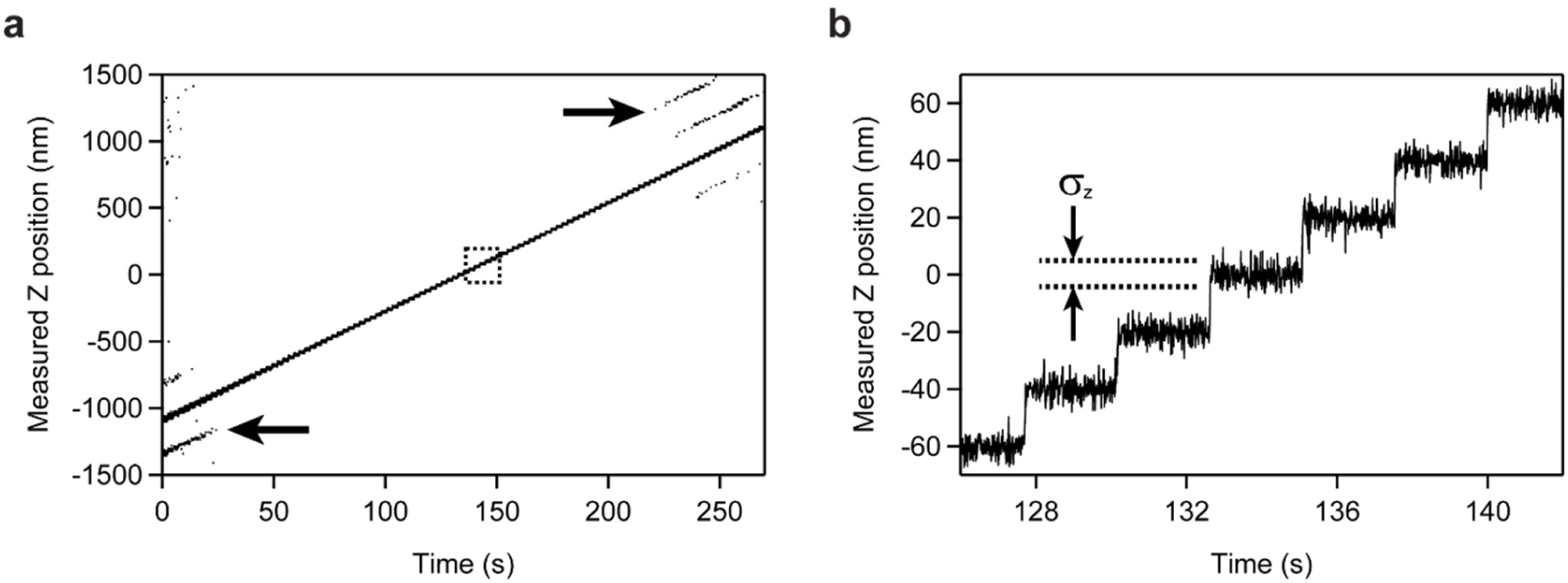
Bead step scan. A fluorescent bead (100 nm diameter) bound to a glass coverslip was scanned through the focal plane of the microscope in 20 nm steps, to measure the localization precision and the frequency of localization artifacts. The illumination was adjusted such that the number of detected photons per camera frame was similar to the brightness of a single fluorophore (∼9200 photons per frame). For every image in the scan, the bead position was measured by fitting the image with the cubic spline PSF model. **(a)** Measured *z* position of the bead during the scan. At the beginning and end of the scan, localization artifacts are visible as periodic bands below and above the true position of the bead (arrows). These are localization errors (artifacts) which arise when the fit algorithm assigns the bead position to the wrong interference fringe of the PSF. The fraction of localization errors can be calculated by comparing the number of correctly localized events to the number of events in the bands above and below the central scan line. **(b)** Zoomed view of the boxed region in (a). The detail of the scan shows the individual steps. At each scan step, the localization precision was measured by calculating the point-to-point variation in the position measurement. Sample drift was estimated using a low-pass filter, and subtracted from the step scan data.

**Figure S19:**
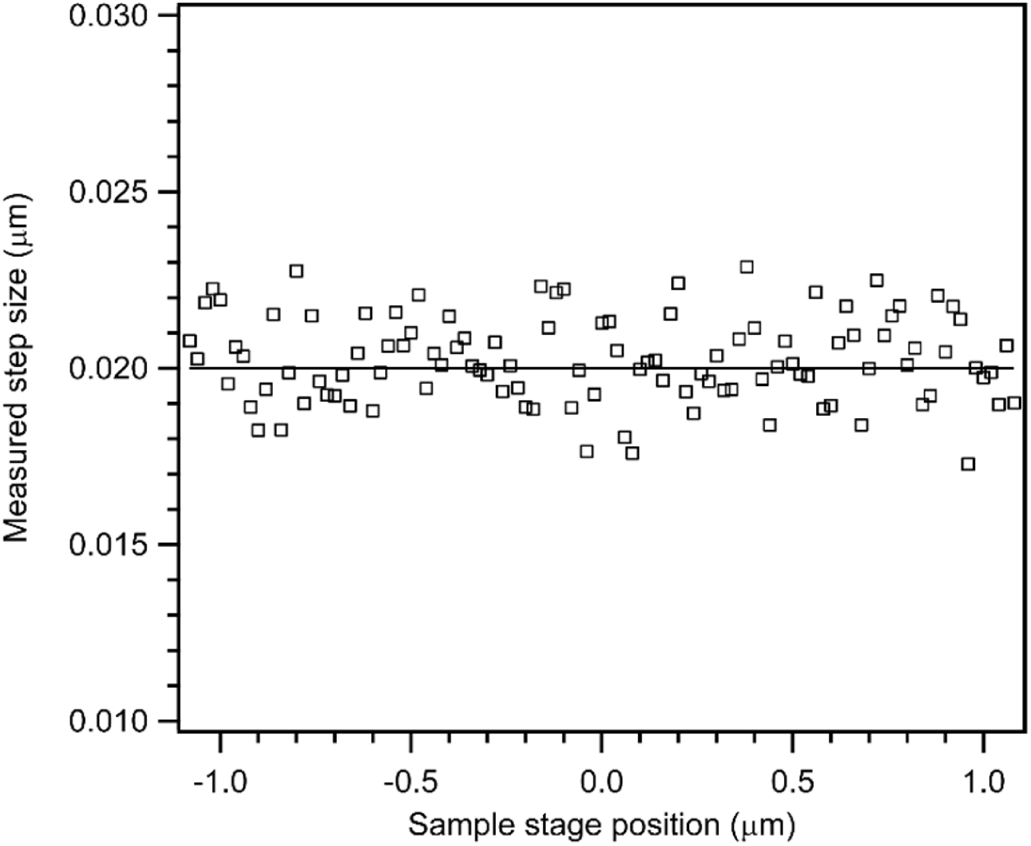
*z*-coordinate localization accuracy. A fluorescent bead (100 nm diameter) bound to a glass coverslip was scanned through the focal plane of the microscope in 20 nm steps, as in Supplementary Fig. S18. For this measurement, the illumination was adjusted such that more photons were detected from the bead (approximately 25000 per exposure). Here, the measured step sizes of the scan (black squares) are plotted vs. the piezo stage position during the scan, to test for bias in the localization procedure. Scatter in the data points is due to random measurement error at each time point. Within the measurement precision, the estimated step sizes do not deviate from the expected value (solid black line) over the full range of the scan, exhibiting no systematic bias with respect to the *z*-position of the emitter.

**Figure S20:**
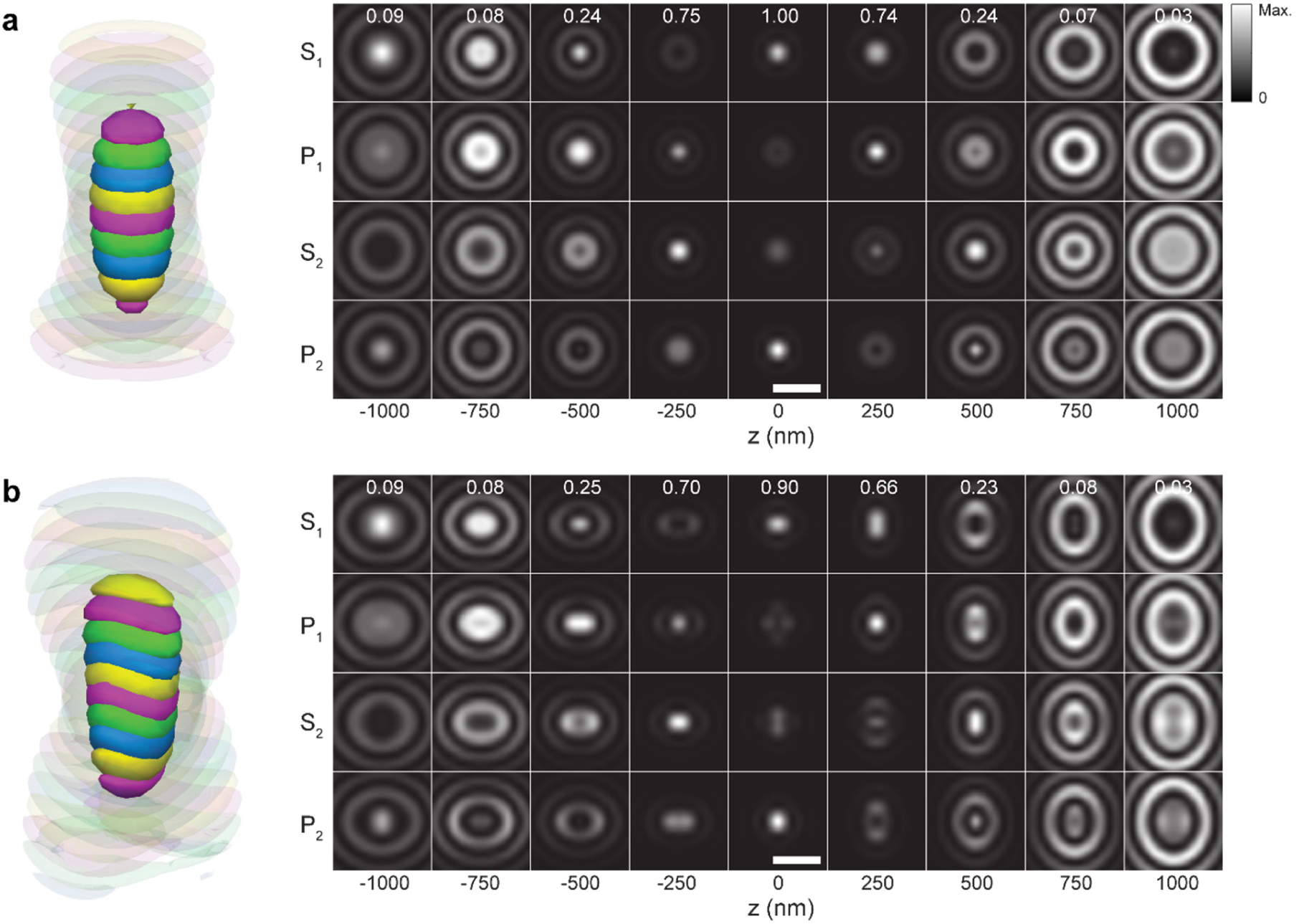
Simulated symmetric and astigmatic 4Pi PSFs. A matrix of *x-y* slices through the simulated symmetric (a) and the astigmatic (b) 4Pi PSFs used for the analysis shown in Fig. 4, at various *z-*positions relative to the focal plane, for the four image channels. Details of the PSF simulations are provided in the Supplementary Methods. A 3D rendering of each PSF is shown to the left of the image matrix. For each z position, all four image channels were scaled to the same maximum value. The value of the maximum, relative to the maximum of the symmetric 4Pi PSF at z=0, is shown at the top of each column. Scale bar: 500 nm.

**Figure S21:**
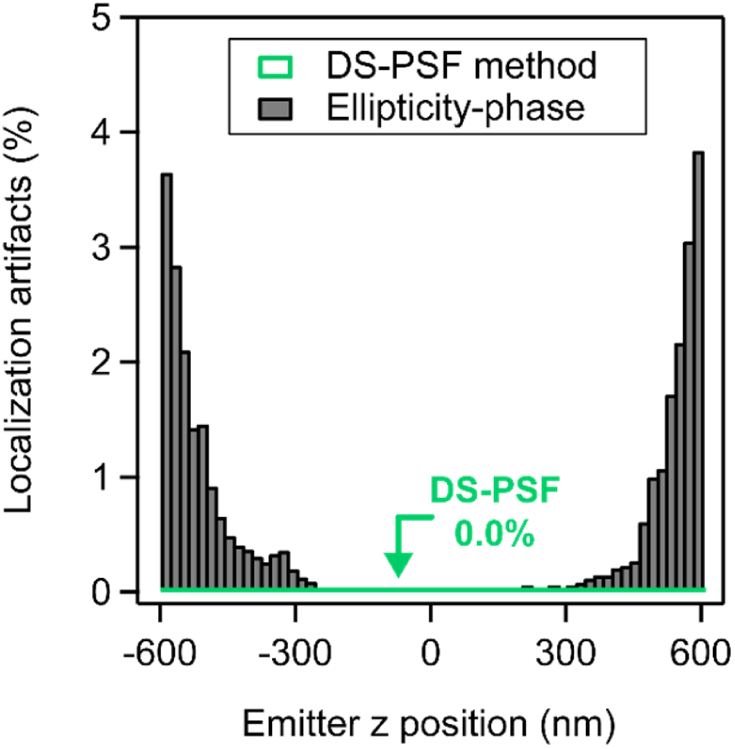
Comparison with Astigmatic 4Pi PSF-based approaches: Localization artifacts. Artifact frequency as a function of the emitter *z-*coordinate, corresponding to the simulated data sets evaluated in Fig. 4. Simulated fluorophore images were generated based on the astigmatic 4Pi PSF and the symmetric 4Pi PSF. Data based on the astigmatic PSF were analysed with the ellipticity-phase method, and data based on the symmetric PSF were analyzed with the DS-PSF method. The simulated datasets were based on experimentally realistic photon counts with a mean of 8000 photons per event, and the size of the emitter cutout region was 13x13 pixels (138 nm pixel size). Both methods obtain a low rate of localization artifacts, with the ellipticity-phase analysis reaching an artifact frequency of ∼4% for positions 600 nm from the focal plane. The artifact frequency for the DS-PSF fit was 0% over the range tested.

**Figure S22:**
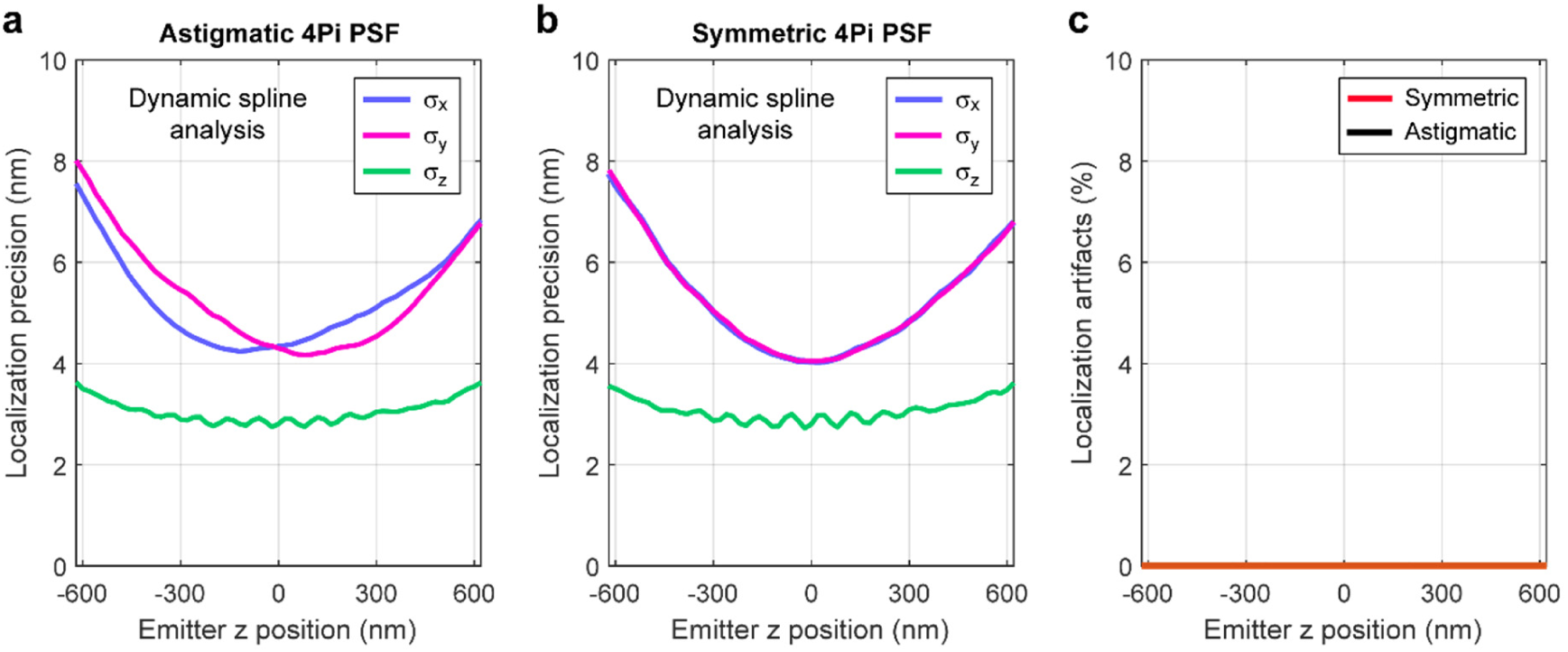
Dynamic spline analysis with Astigmatic or Symmetric 4Pi PSF. Localization precision (a,b) and artifact rate (c) for a simulation of emitters at different *z* positions, using either the astigmatic or the symmetric 4Pi PSF (Fig. 4) to generate and fit the data. Here, both datasets were analyzed with the dynamic spline method, instead of using the ellipticity-phase analysis for the astigmatic PSF. The results show that the localization precision is comparable for the two PSFs with the new analysis, with the symmetric 4Pi PSF giving slightly better results due to its more compact shape. The artifact fraction was < 0.1% over the entire *z-*coordinate range tested. In the simulated data, there were on average 8000 photons per localization, and 10 background photons per pixel.

**Figure S23:**
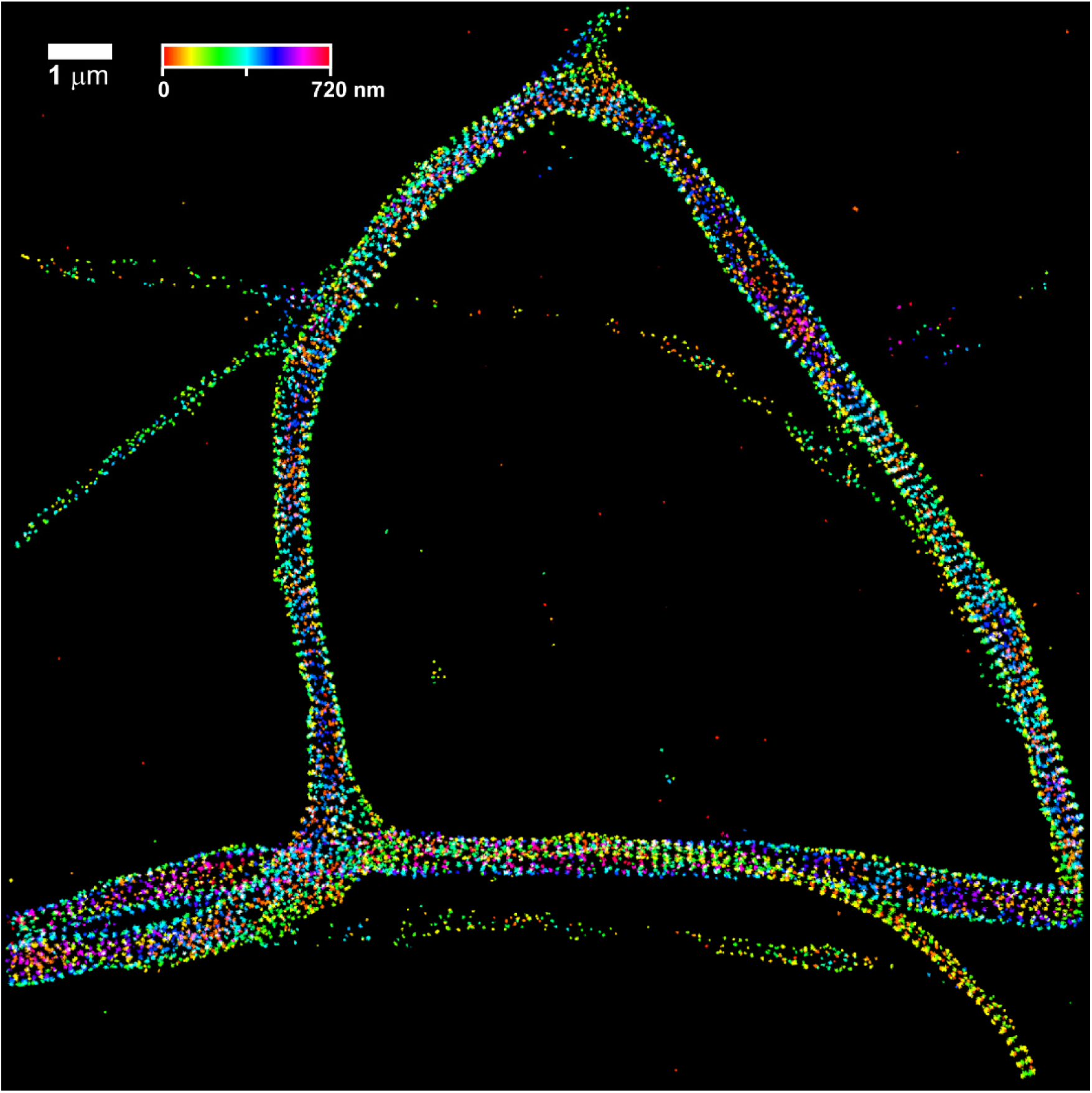
Beta-II spectrin in primary neuron cells. A larger view, showing the entire image, for the neuron sample displayed in Fig. 5. As described, the primary neuron cell is stained with antibodies against beta-II spectrin, and secondary F(ab)2 antibody fragments labeled with Alexa Fluor 647. The localizations were color-coded according to their *z-*coordinate to show the depth information. Three-dimensional renderings of this dataset are shown in Supplementary Videos S2 and S3. Scale bar: 1 µm.

**Figure S24:**
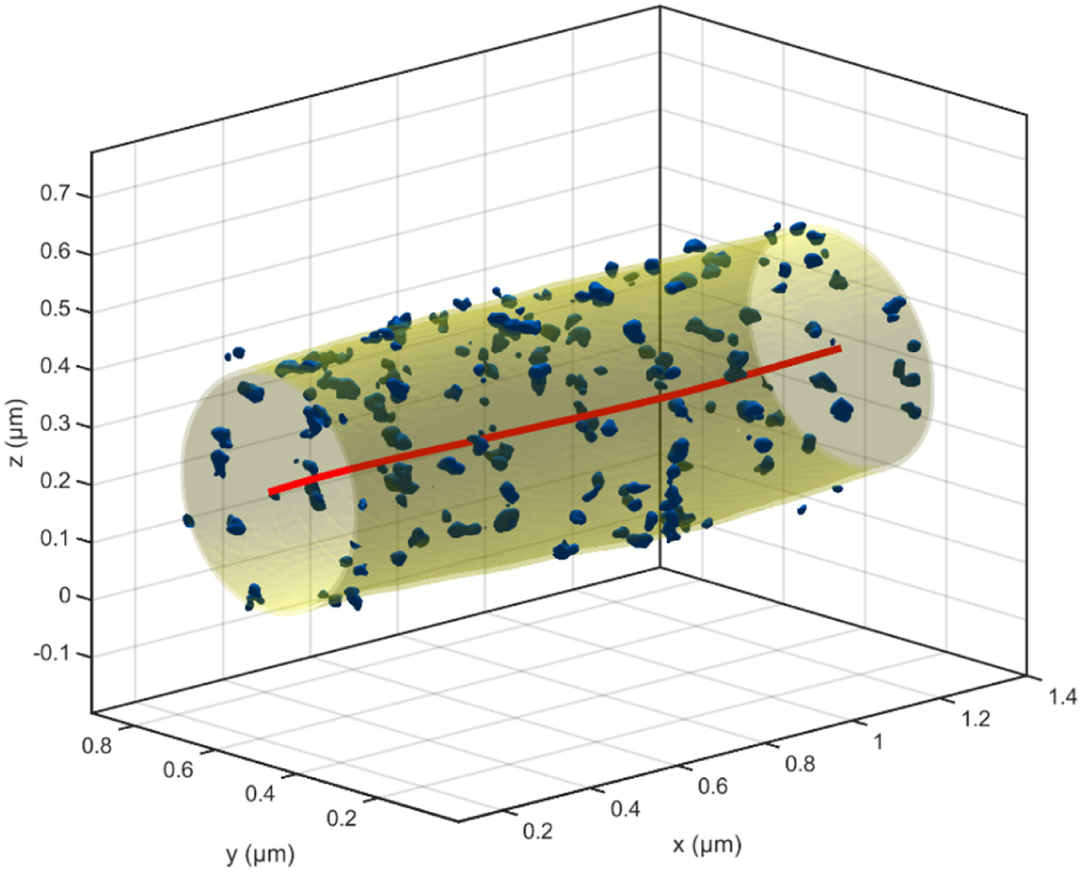
Mapping the surface of a neuronal process. Beta-II spectrin volume rendering corresponding to the data in Fig. 5 b-e including the determination of the center line (red) and the tube-like surface (yellow) that best contains the beta-II spectrin localization data (see Supplementary Notes). An animation illustrating the detection of the axonal membrane region, which is needed to generate the unwrapped view of beta-II spectrin, is shown in Supplementary Video S4.

**Figure S25:**
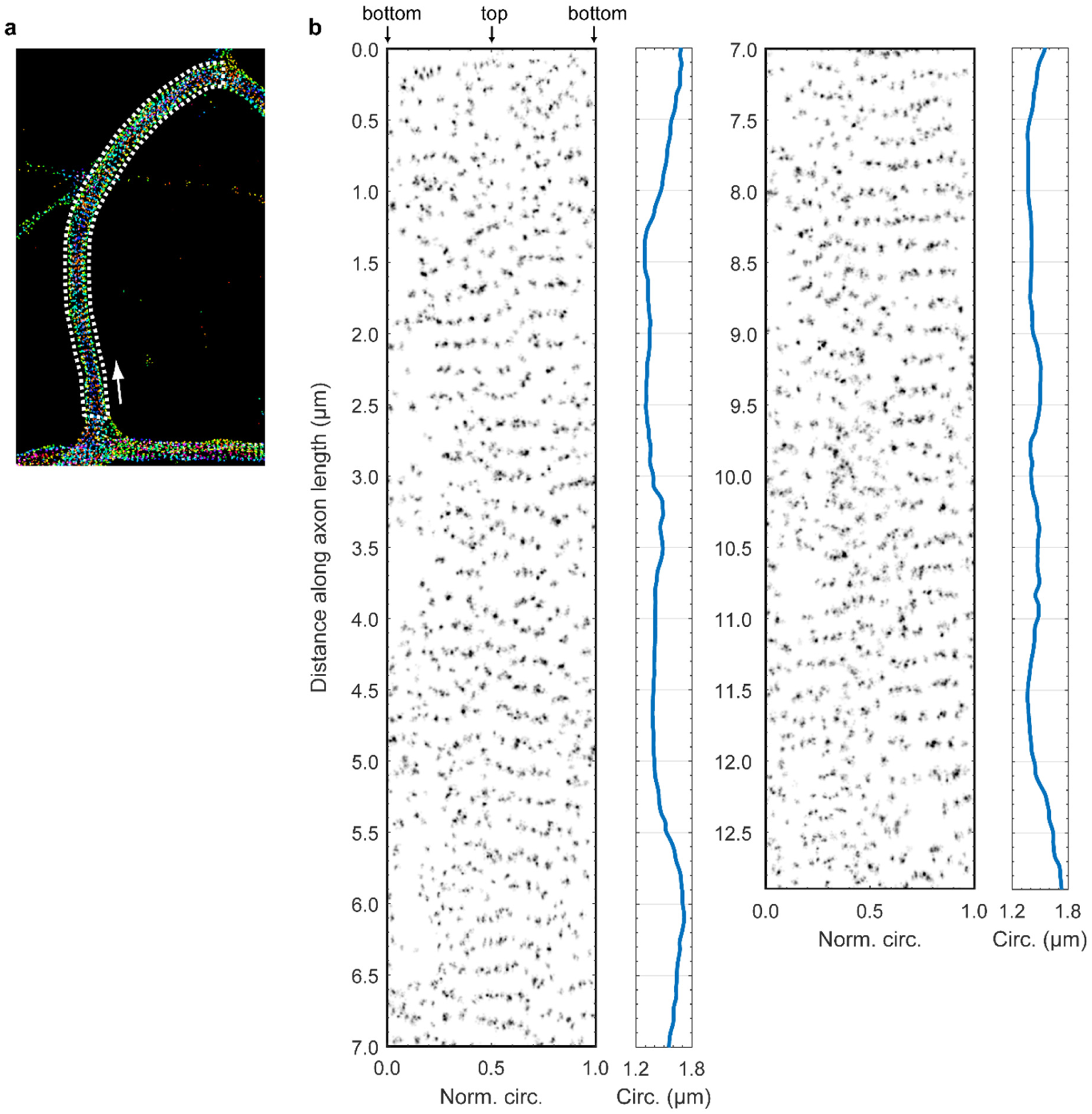
Unwrapped view of Beta-II spectrin (region 1) . A longer section of the neuron shown in Fig. 5, digitally unwrapped to show the organization of beta-II spectrin in the cell. **(a)** Overview image, showing the region which was unwrapped (white box). One end of the box marked as the origin (coordinate 0) of the analysis, and the direction of propagation is indicated by the white arrow. **(b)** The organization of beta-II spectrin relative to the plane of the neuronal membrane. The spectrin density is plotted as a function of the normalized distance around the circumference of the neuron (*x-*axis) and the distance along the length of the process (*y*-axis). The measured circumference of the neuron is plotted to the right of the spectrin density.

**Figure S26:**
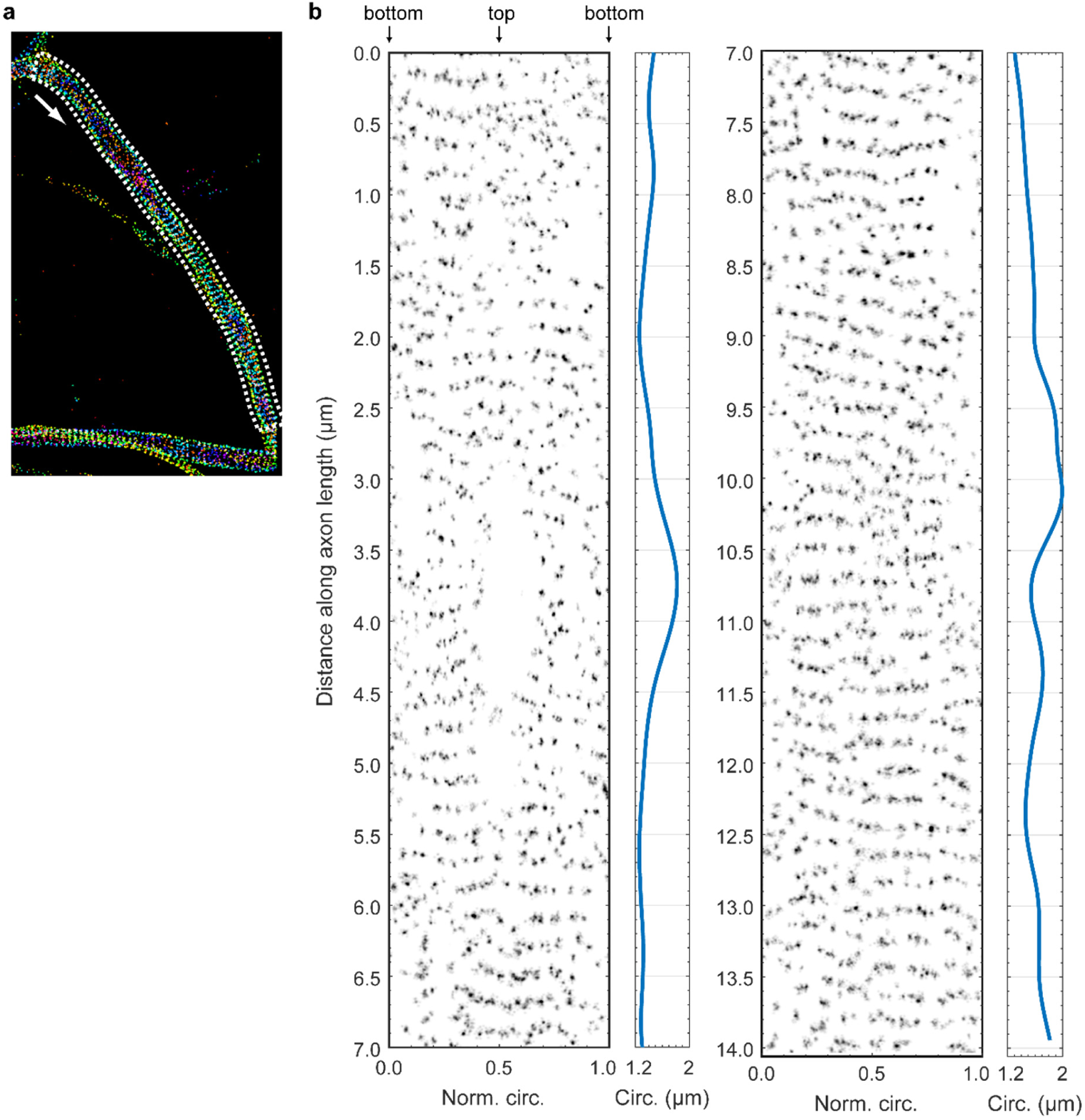
Unwrapped view of Beta-II spectrin (region 2). A longer section of the neuron shown in Fig. 5, digitally unwrapped to show the organization of beta-II spectrin in the cell. **(a)** Overview image, showing the region which was unwrapped (white box). One end of the box marked as the origin (coordinate 0) of the analysis, and the direction of propagation is indicated by the white arrow. **(b)** The organization of beta-II spectrin relative to the plane of the neuronal membrane. The spectrin density is plotted as a function of the normalized distance around the circumference of the neuron (*x-*axis) and the distance along the length of the process (*y*-axis). The measured circumference of the neuron is plotted to the right of the spectrin density.

**Figure S27:**
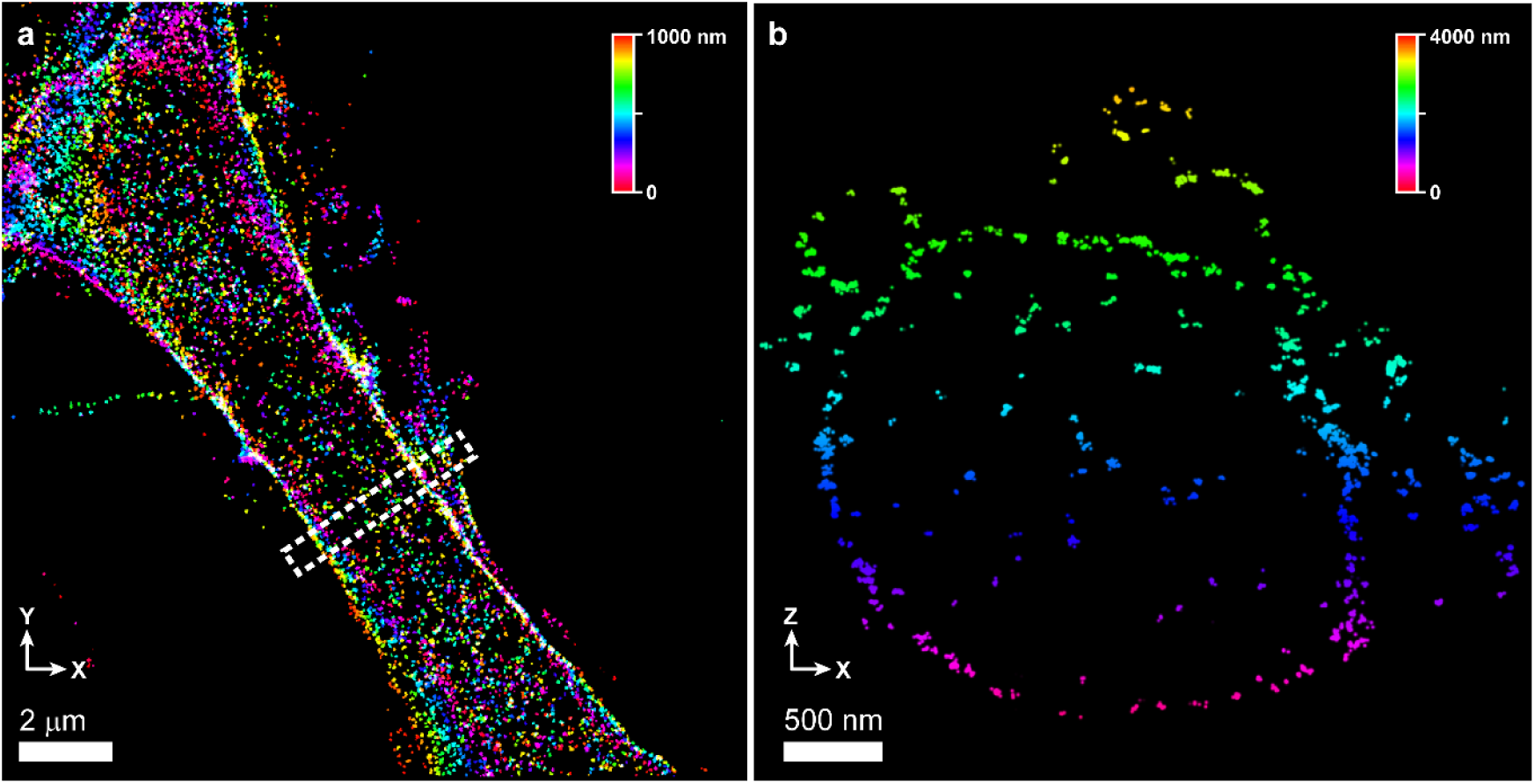
Beta-II spectrin in a thick neuronal process. A primary neuronal cell, several micrometers in thickness, labeled with antibodies against beta-II spectrin. **(a)** An average projection through a top view (*x-y*) of the cell, with the viewplane positioned in the center of the sample. The image shows a high density of spectrin at the surface, and a low density inside the cytoplasm. Scale bar: 2 µm. **(b)** A cross-section view (*x-z*) through the boxed region in (a), showing the circular profile of the cell. Several neuronal processes are visible, clustered around the surface of the larger cell. This dataset was obtained by shifting the sample with respect to the objective lenses during the recording, in ten steps separated by 400 nm, for a total imaging depth of approximately 5 micrometers. Notably, localization artifacts are not evident in the image, despite the large extent of the sample in the *z-*dimension. Scale bar: 500 nm.

**Figure S28:**
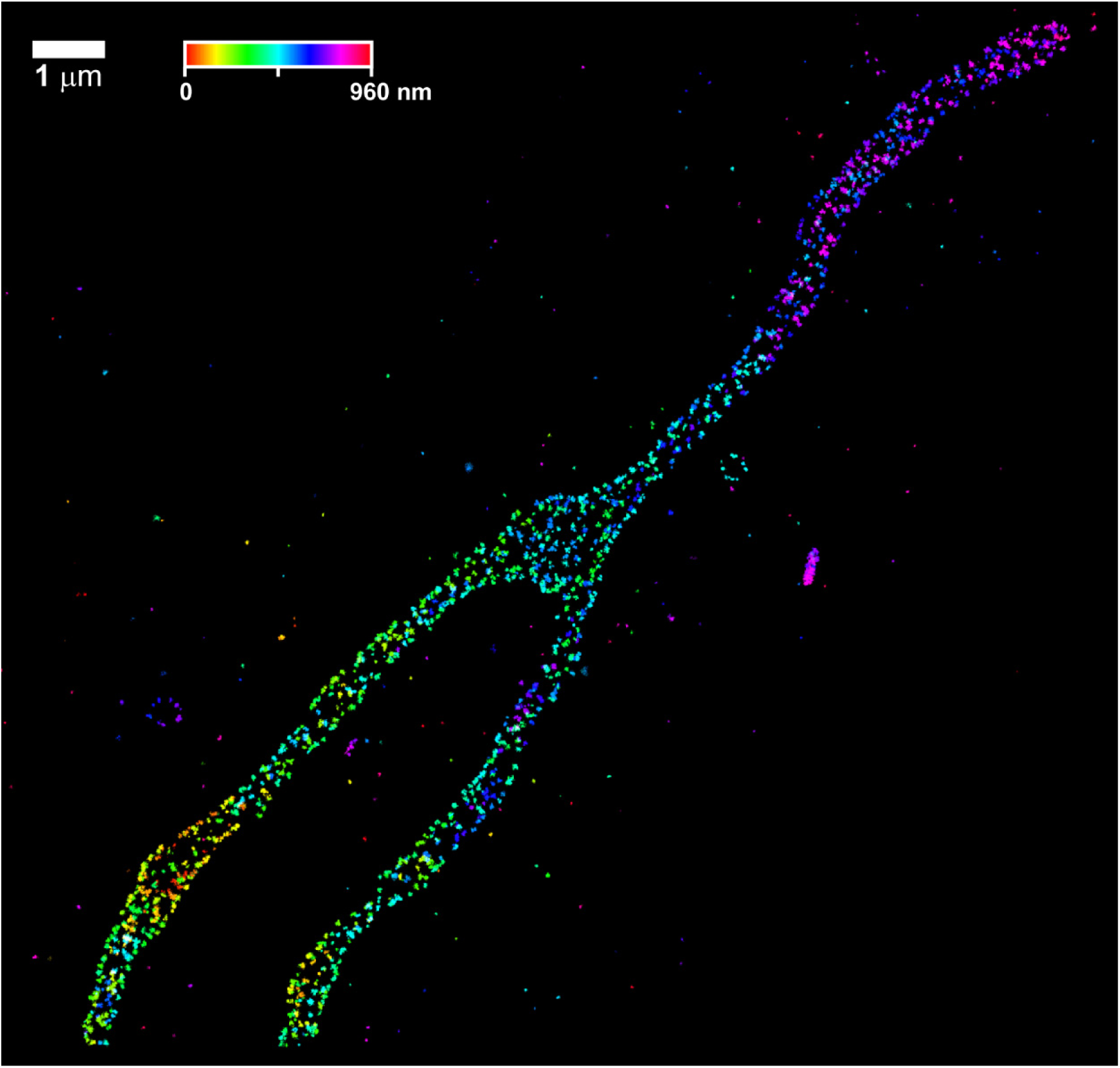
Mitochondrial Mic60 in U-2 OS cells. A larger view, showing the entire image, for the sample displayed in Fig. 6a. As described, the cell is stained with antibodies against Mic60, and secondary antibodies labeled with Alexa Fluor 647. The localizations were color-coded according to their *z-*coordinate to show the depth information. A three-dimensional rendering of this dataset is shown in Supplementary Video S5. Scale bar: 1 µm.

**Figure S29:**
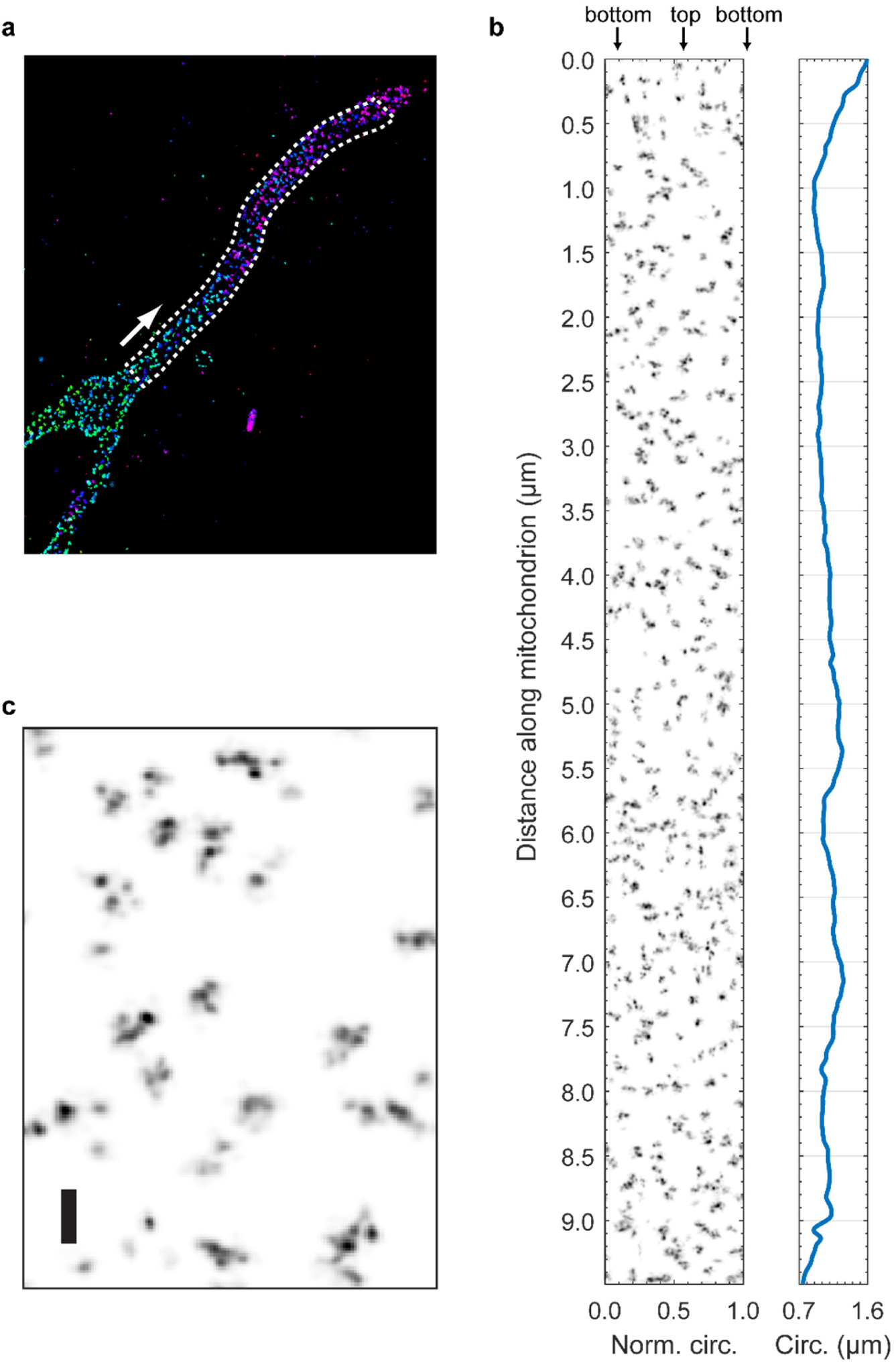
Unwrapped view of Mic60 in a U-2 OS cell. A longer section of the mitochrondrion shown in Fig. 6a, digitally unwrapped to show the organization of Mic60 at the inner boundary membrane. **(a)** Overview image, showing the region which was unwrapped (white box). One end of the box marked as the origin (coordinate 0) of the analysis, and the direction of propagation is indicated by the white arrow. **(b)** The organization of Mic60 relative to the plane of the mitochondrial membrane. Mic60 density is plotted as a function of the normalized distance around the circumference of the membrane (*x-*axis) and the distance along the length of the mitochondrion (y-axis). The measured circumference is plotted to the right of the Mic60 density. **(c)** Zoomed view of data from (b). Scale bar: 100 nm.

**Figure S30:**
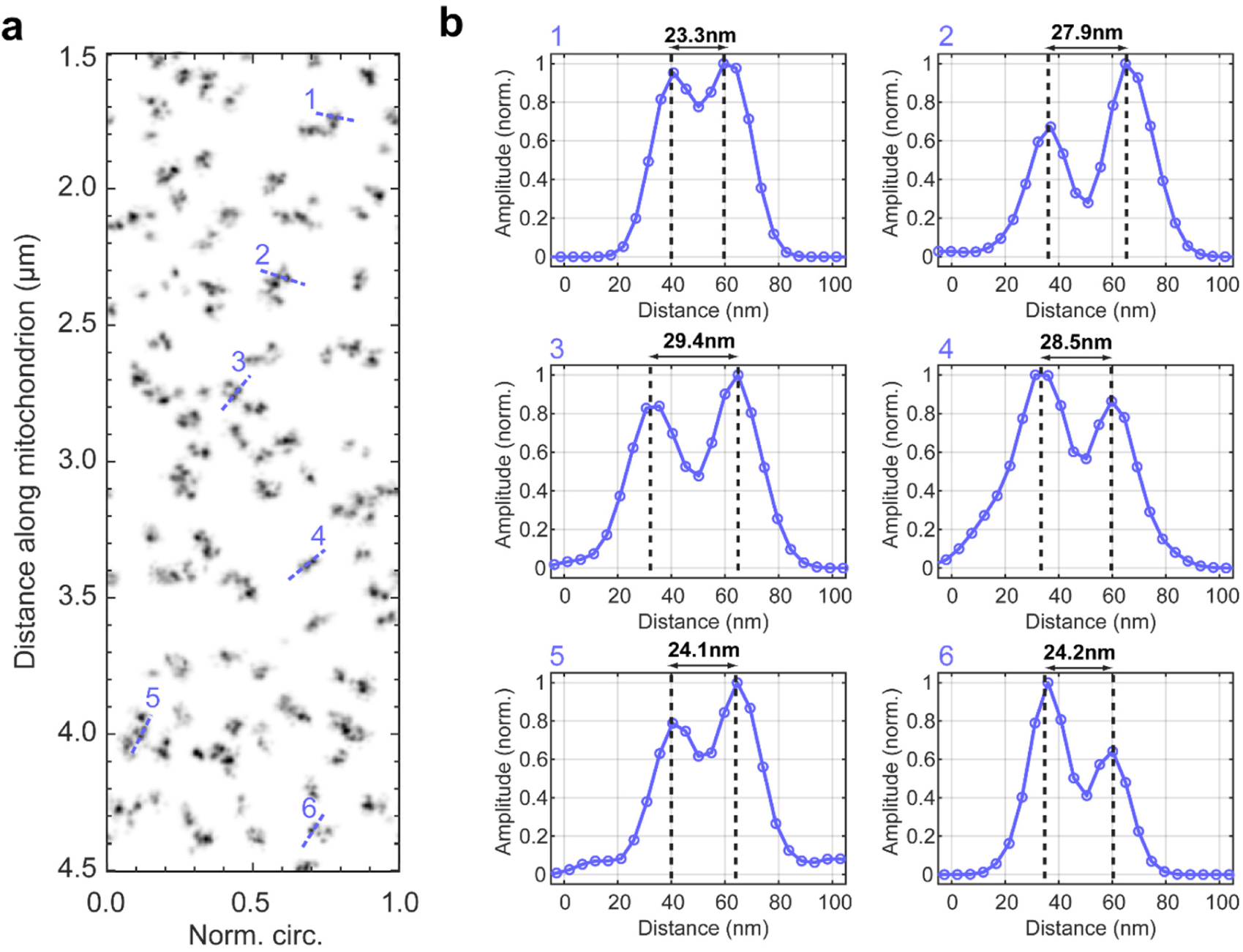
Measurements of Mic60 organization in a U-2 OS cell. **(a)** A 3 µm long section of an unwrapped view of Mic60 in a U-2 OS cell, showing the location of six line profiles (blue, dotted lines) that cross neighboring Mic60 spots in different Mic60 puncta. **(b)** Corresponding line profiles through the unwrapped views and measured distances between peaks. Note that the unwrapping of a curved surface onto a rectangle slightly distorts the scale (average circumference within the section was used).

**Figure S31:**
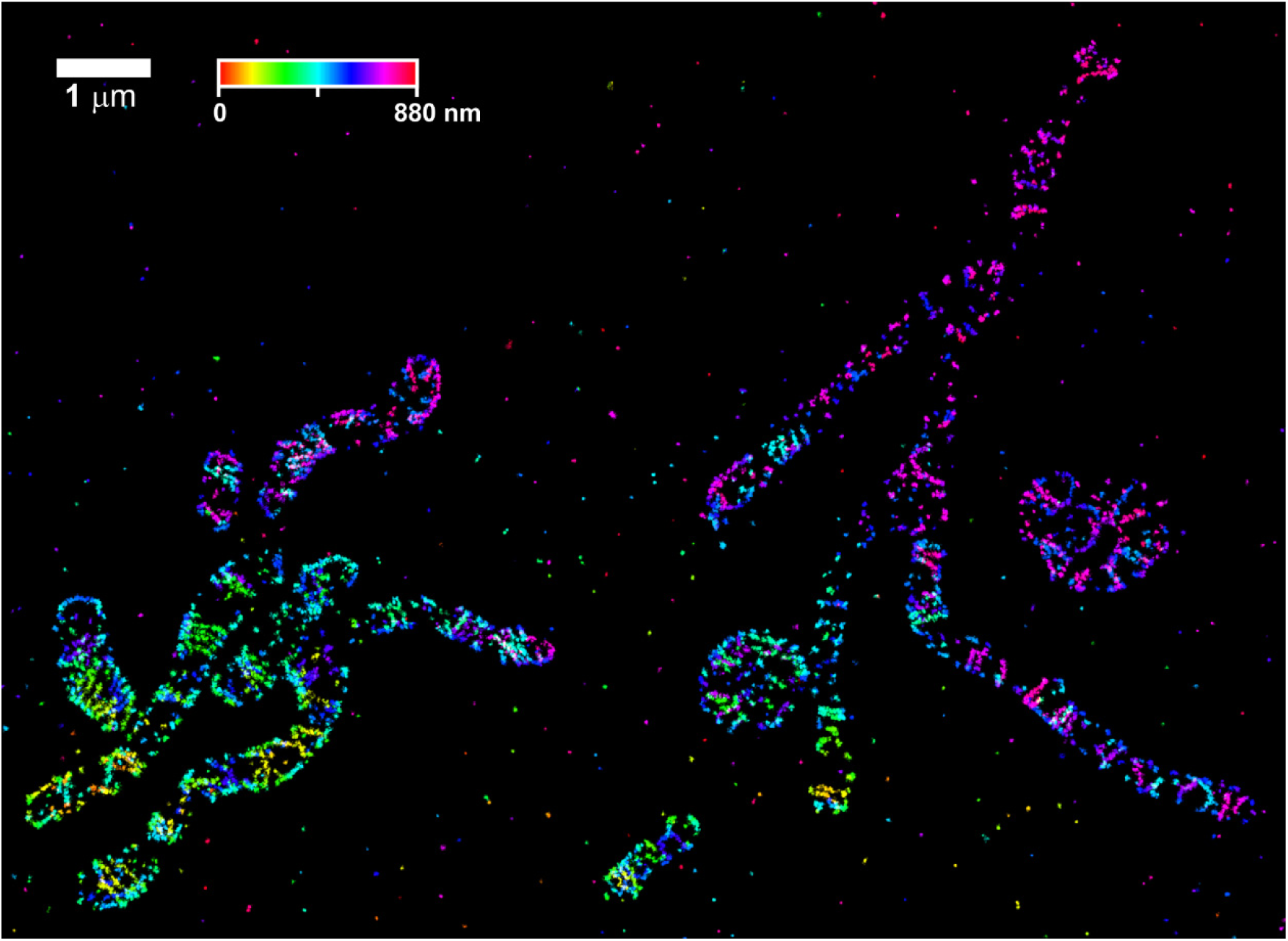
Mitochondrial Mic60 in COS7 cells. A larger view, showing the entire image, for the sample displayed in Fig. 6c. As described, the cell is stained with antibodies against Mic60, and secondary antibodies labeled with Alexa Fluor 647. The localizations were color-coded according to their *z-*coordinate to show the depth information. A three-dimensional rendering of this dataset is shown in Supplementary Video S6. Scale bar: 1 µm.

**Figure S32:**
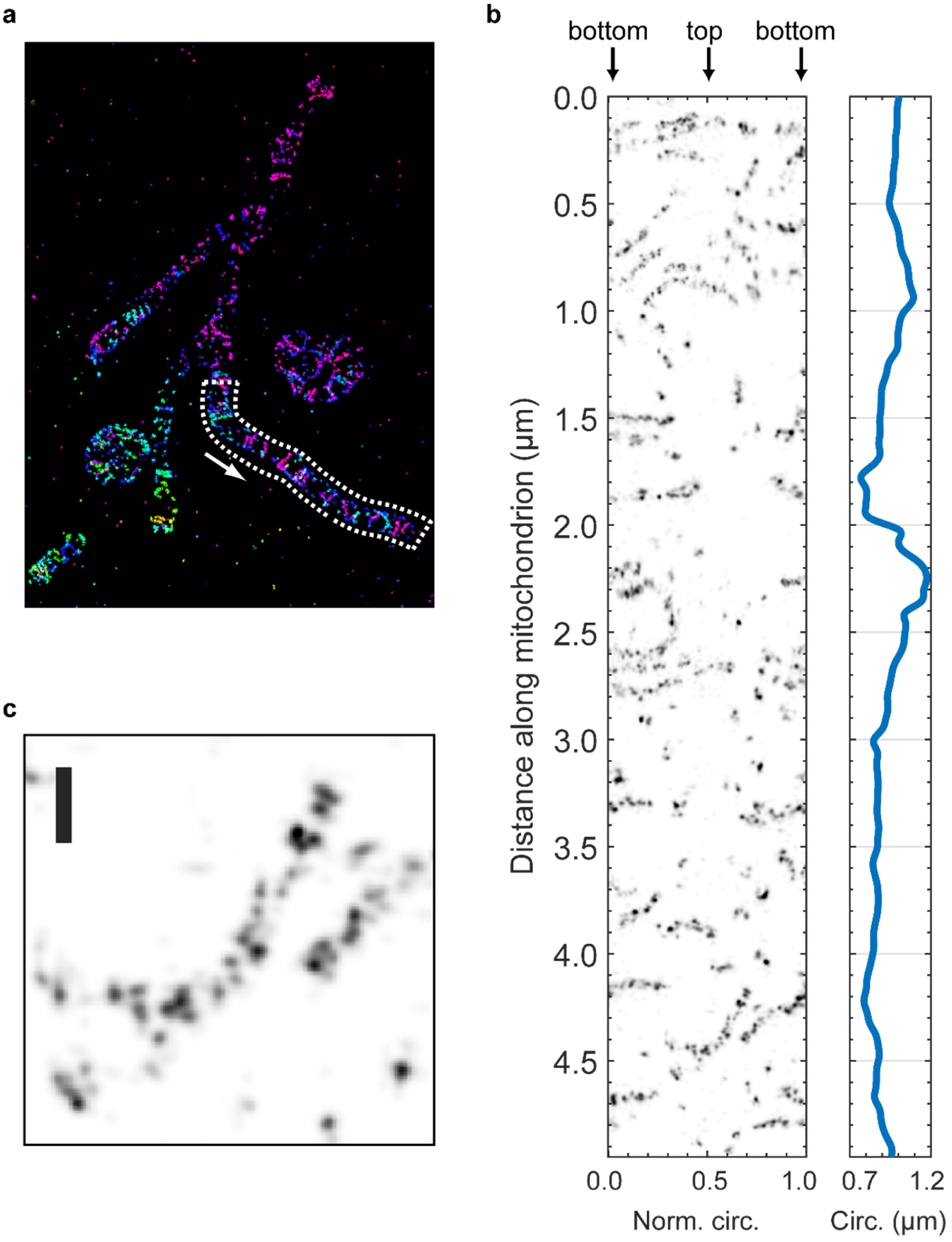
Unwrapped view of Mic60 in a COS-7 cell. A longer section of the mitochrondrion shown in Fig. 6c, digitally unwrapped to show the organization of Mic60 at the inner boundary membrane. **(a)** Overview image, showing the region which was unwrapped (white box). One end of the box marked as the origin (coordinate 0) of the analysis, and the direction of propagation is indicated by the white arrow. **(b)** The organization of Mic60 relative to the plane of the mitochondrial membrane. Mic60 density is plotted as a function of the normalized distance around the circumference of the membrane (*x-*axis) and the distance along the length of the mitochondrion (y-axis). The measured circumference is plotted to the right of the Mic60 density. **(c)** Zoomed view of data from (b). Scale bar: 100 nm.

**Figure S33:**
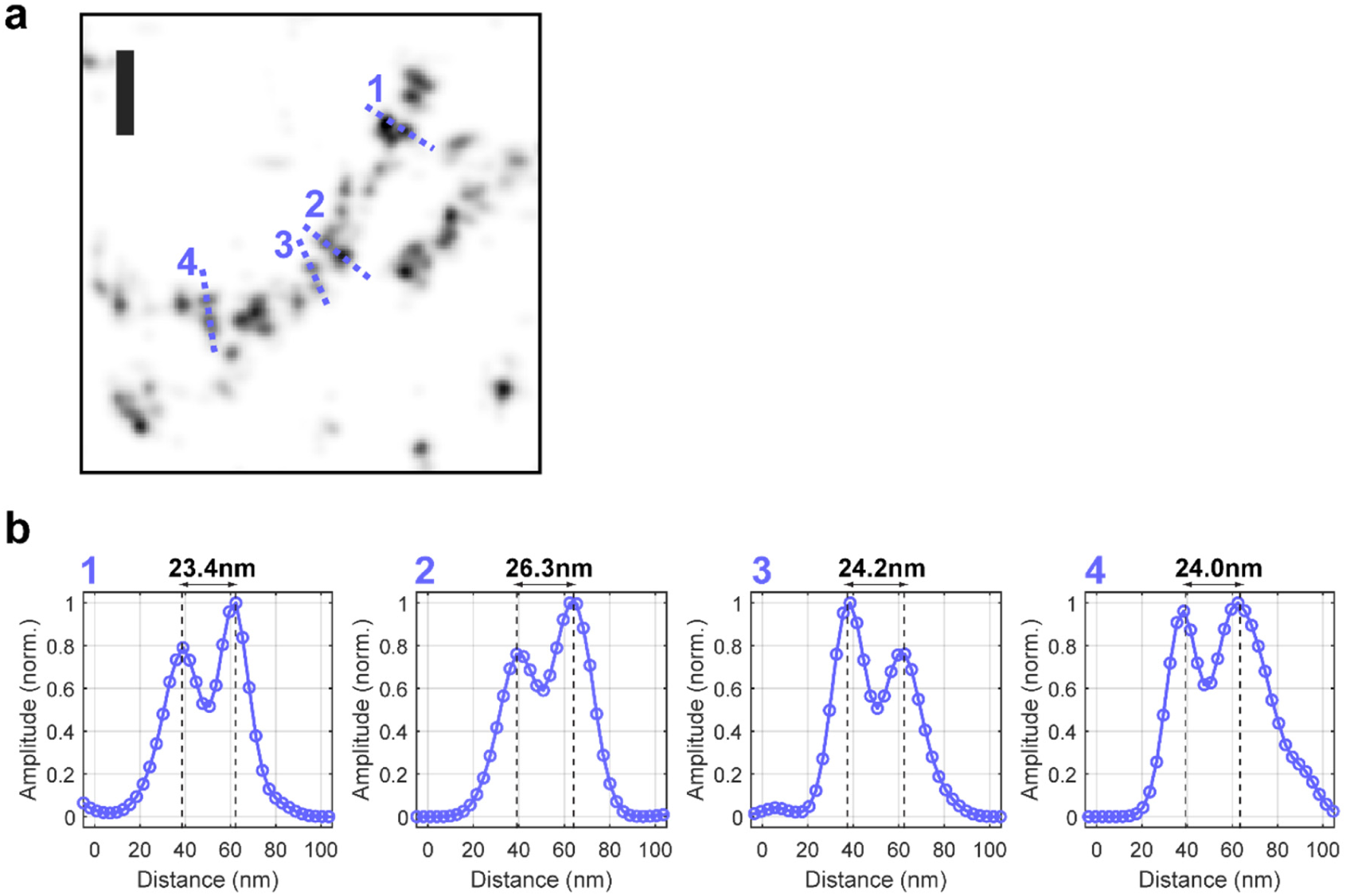
Measurements of Mic60 organization in a COS-7 cell. **(a)** Zoomed view of an unwrapped section of mitochondrial Mic60 in a COS-7 cell showing the location of four line profiles (blue, dotted lines) that are perpendicular to the orientation of the apparent Mic60 stripe. Scale bar: 100 nm **(b)** Corresponding line profiles through the unwrapped views and measured distances between peaks. Note that the unwrapping of a curved surface onto a rectangle slightly distorts the scale (average circumference within the section was used).

**Figure S34:**
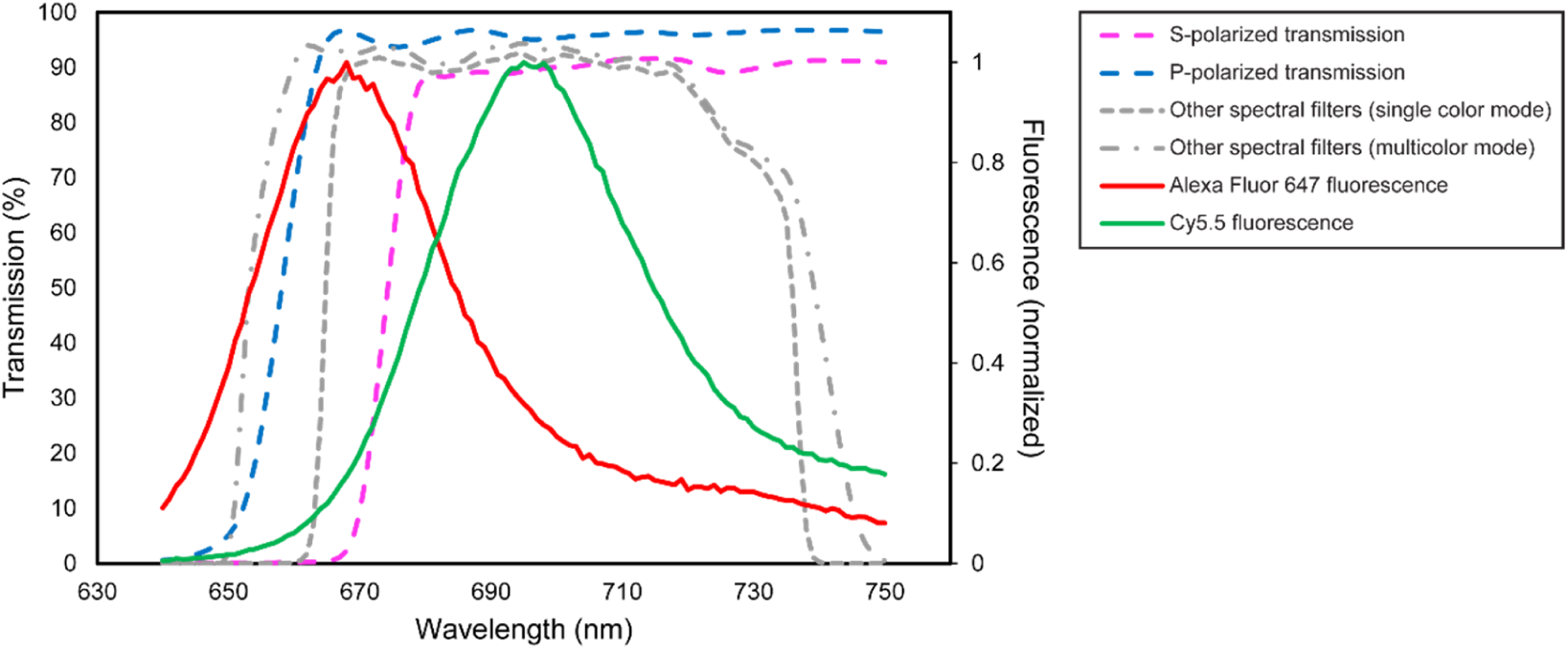
Polarization-specific spectral filter for multicolor imaging. Discrimination of Alexa Fluor 647 and Cy5.5 fluorophores is achieved using a spectral long-pass filter which has a different cutoff wavelength for S- and P-polarized light. The plot shows the fluorescence emission spectra of the two dyes (solid lines) whose emission maxima are separated by approximately 30 nm. A dichroic mirror placed in each detection path (Supplementary Fig. S1), placed at an angle of ∼34 degrees with respect to the fluorescence beam, has a different transmission spectrum for each polarization (blue and magenta dashed lines). Specifically, the S-polarized cutoff wavelength is red-shifted by ∼15 nm with respect to the cutoff for P-polarized light. This filter configuration introduces a significant difference in the ratios of detected S- and P-polarized photons for Alexa Fluor 647 and Cy5.5, allowing the two fluorophores to be distinguished by this parameter. Also shown are the combined spectra for the other dichroic mirrors and spectral filters in the system (grey dashed lines), for the single color and multicolor configurations (see Methods).

**Figure S35:**
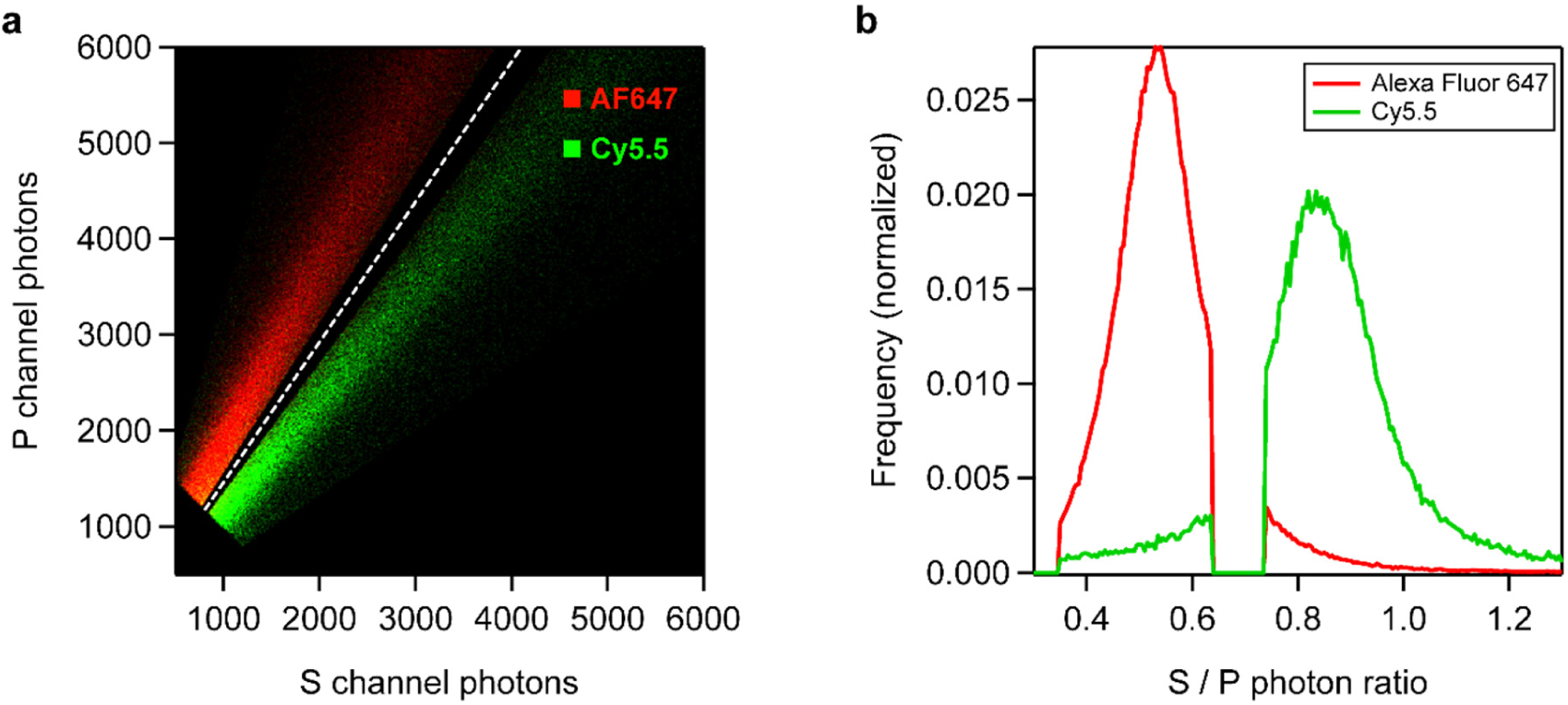
Color discrimination and crosstalk estimation. Control experiment using samples labeled with a single color (Alexa 647 or Cy5.5) and imaged with the two-color imaging configuration (dichroic mirrors DM3 in the detection path, Supplementary Fig. S1). **(a)** Two-dimensional histogram showing the number of photons detected in the P-polarized channels (p1 + p2) vs. the number detected in the S-polarized channels (s1 + s2) for each localization event, for the two control samples (Alexa 647 and Cy5.5). Events for each fluorophore appear in two groups, separated by the ratio of S to P photons. A white dashed line indicates the threshold value used for initial identification of the two fluorophores in the multicolor analysis. **(b)** Histogram of S/P photon ratio for the two control samples, showing the expected mis-identification fraction (crosstalk). Events with an S/P ratio < 0.7 are identified as Alexa 647, and with > 0.7 as Cy5.5. Events with high ambiguity in color identity (S/P ratio between 0.64 and 0.74) were filtered out to reduce crosstalk. Overall, 93% of localization events are correctly identified.

**Figure S36:**
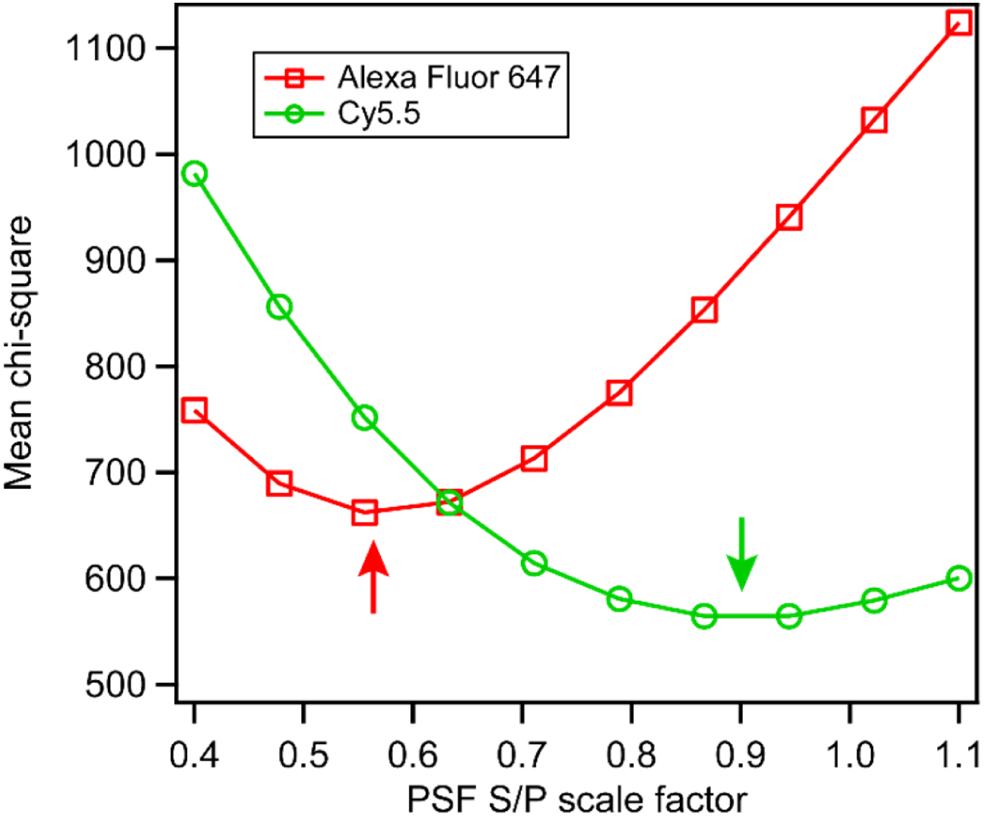
Multicolor PSF S/P scale factor determination. The correct scaling of the PSF S- and P-channel amplitudes may be determined from the localization data. In this example, localizations from the two-color dataset shown in Fig. 6 (Mic60 & DNA) were initially assigned colors based on the ratio of detected S-channel photons to P-channel photons, using a simple threshold of 0.7 (see Supplementary Fig. S35). Next, a series of test PSFs were generated by rescaling the S-channel of the PSF spline relative to the P-channel, over a range of 0.4 to 1.1. Each color group of localizations was fit with the test PSFs, and for each group the mean chi-square of the fits is plotted vs. the channel scaling factor. The minimum of each curve (arrows) reveals the PSF S- to P-channel amplitude ratio for each dye which best corresponds to the experimental data. In this case, the optimal scaling factor was 0.57 for Alexa Fluor 647, and 0.91 for Cy5.5, determined with a parabolic fit.

**Figure S37:**
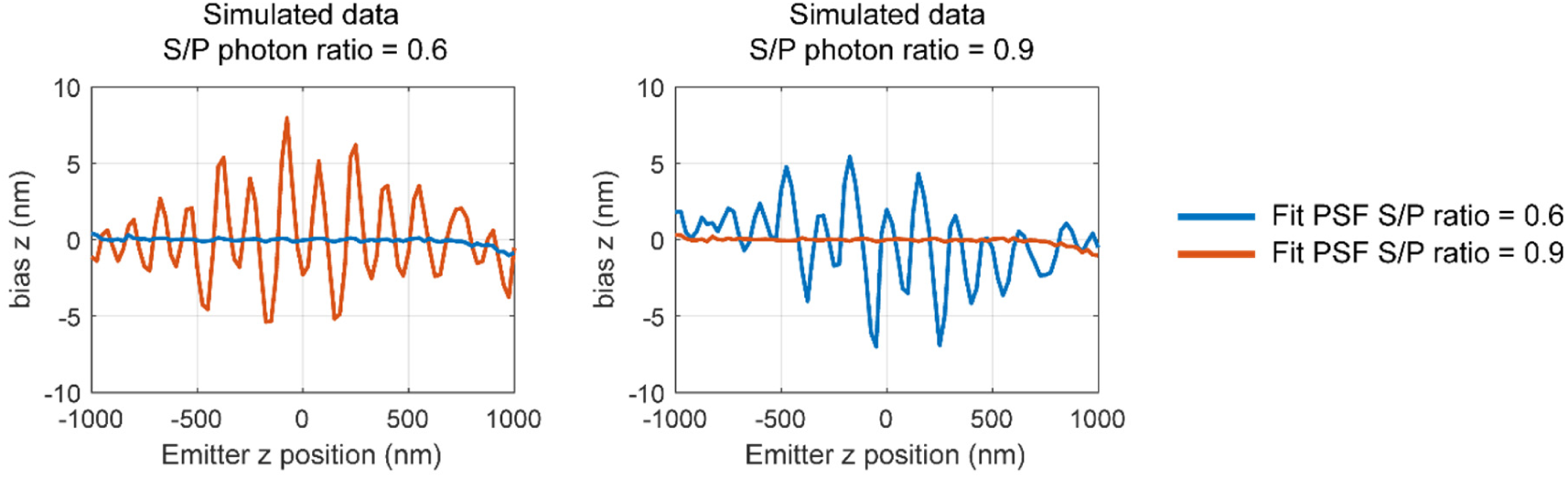
*z-*coordinate bias due to fitting with an incorrectly scaled PSF (simulation). Simulation of the error in the fit results which would be introduced if the incorrectly scaled PSF is used for fitting in multicolor 4Pi-STORM analysis. Simulated emitter images were generated for Alexa Fluor 647 and Cy5.5, having an S- to P-polarized photon ratio of 0.6 and 0.9, respectively. Both sets of data were then fit with the correctly scaled PSF (ratio=0.6 for Alexa Fluor 647 and ratio=0.9 for Cy5.5) and the incorrectly scaled PSF. For each fluorophore, the error in the estimated *z-*coordinate is plotted as a function of the *z-*coordinate of the emitter.

**Figure S38:**
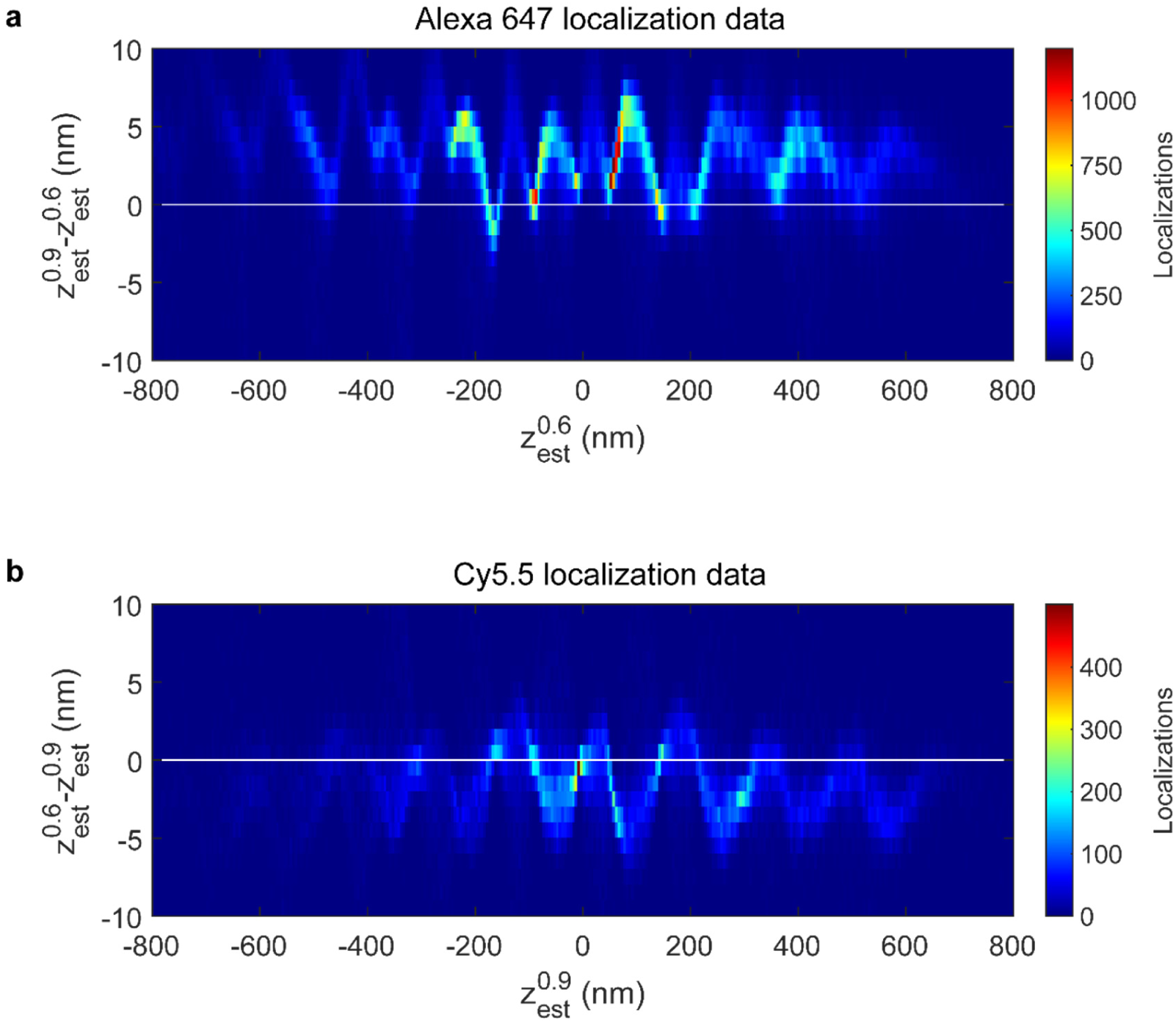
*z-*coordinate bias due to fitting with an incorrectly scaled PSF (experiment). Experimental test of the error introduced when a PSF model with the incorrect S/P channel amplitude ratio is used to fit the emitter images. Localizations from a two-color 4Pi-STORM dataset were sorted into two groups, corresponding to the two fluorophores (Alexa Fluor 647 and Cy5.5), based on the measured ratio of S- to P-polarized detected photons. Next, each group was fit with both the correctly scaled PSF (ratio=0.6 for Alexa 647 and ratio=0.9 for Cy5.5) and the incorrectly scaled PSF. For each group, the discrepancy in the *z-*coordinate between the two fits is plotted as a function of the estimated *z-*coordinate of the emitter obtained by the fit with the correctly scaled PSF model. This amounts to a measurement of the *z-*bias error which would be introduced by using the wrong PSF. **(a)** *z-*bias error for Alexa Fluor 647, as a function of emitter z position. **(b)** *z-*bias error for Cy5.5, as a function of emitter z position.

**Figure S39:**
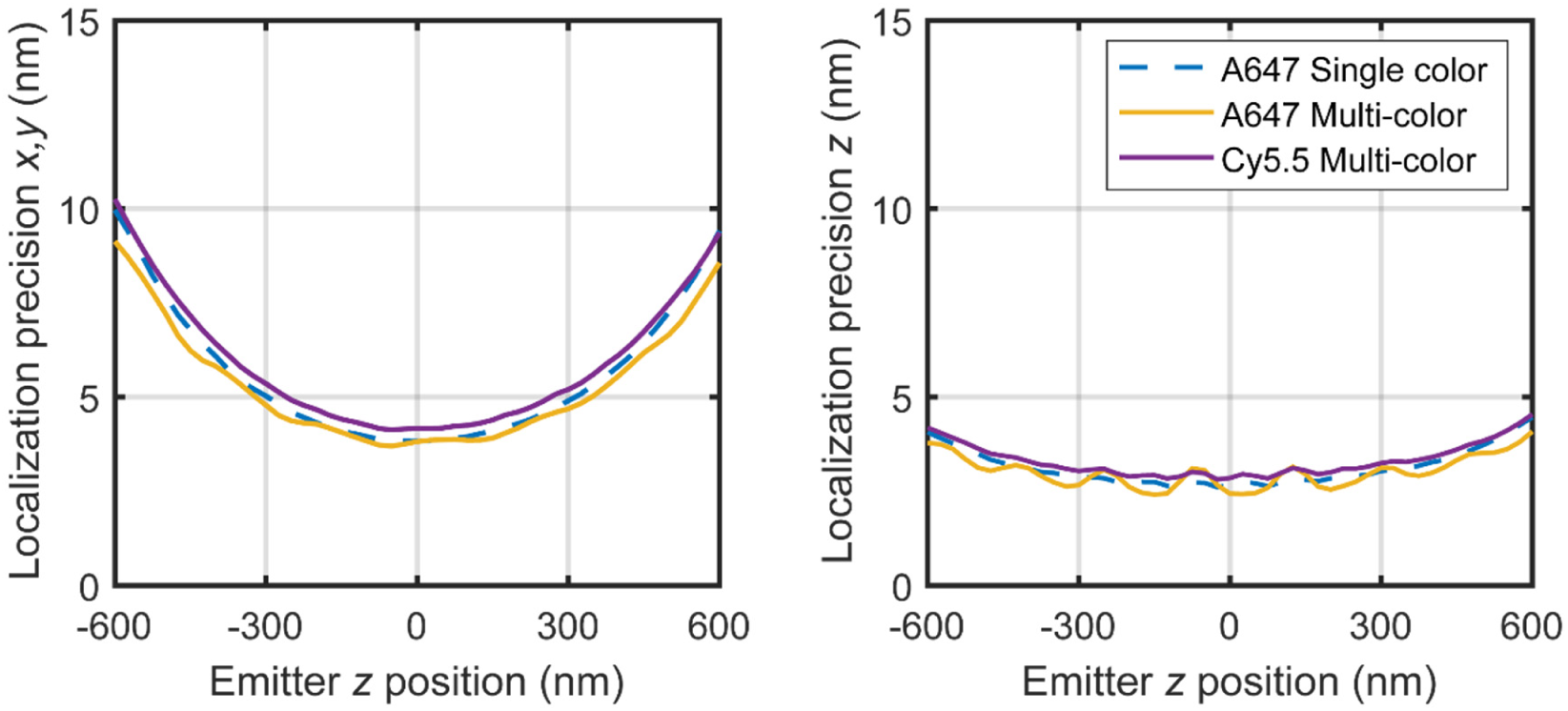
Single color vs. multicolor localization precision. Simulation of localization precision for the fluorophores used in the single color and multicolor imaging configurations. Images of fluorescent emitters were simulated over a range of *z* positions, using mean photon counts and background levels obtained for Alexa Fluor 647 and Cy5.5 in experimental datasets (samples 6 and 7, Supplementary Table S1). On average, the simulated localization uncertainty was 10% higher for Cy5.5 as compared to Alexa Fluor 647, due to the lower number of photons detected. Also, for multicolor imaging of Alexa Fluor 647, the simulated *z* localization precision is observed to modulate with an amplitude of approx. 1 nm close to the focal plane. This variation arises due to the uneven distribution of fluorescence between the S- and P-channels in this configuration.

**Table S1:** 4Pi-STORM localization data statistics

| Sample number | Sample description | Figure | Number of localizations <sup>(1)</sup> | Mean photons per event <sup>(2)</sup> | Mean event duration (frames) <sup>(3)</sup> | Mean background photons per frame <sup>(4)</sup> |
| --- | --- | --- | --- | --- | --- | --- |
| 1 | Nup96 in U2-OS cell | 2 | 27267 | 11923 | 1.4 | 17 |
| 2 | Alexa Fluor 647 - DNA | 3 | 14330 | 8078 | 1.5 | 7 |
| 3 | Beta-II Spectrin in Neuron | 5 | 391994 | 10071 | 2.2 | 26 |
| 4 | Beta-II Spectrin in Neuron (thick sample) | S27 | 242090 | 9246 | 1.8 | 30 |
| 5 | Mic60 in U2-OS cell | 6 | 240711 | 11989 | 1.5 | 52 |
| 6 | Mic60 in COS-7 cell | 6 | 302018 | 10002 | 1.8 | 31 |
| 7 | Mic60 (AF647) and DNA (Cy5.5) in COS-7 cell | 6 | 590623 (AF647)<br>138855 (Cy5.5) | 10066 (AF647)<br>8071 (Cy5.5) | 1.6 (AF647)<br>1.7 (Cy5.5) | 26 (P)<br>16 (S) |

**Notes:**
(1) Number of localizations in the full dataset, after localization filters have been applied (see Supplementary Table S2).
(2) Photons per localization event, calculated as the total of all photons detected per on-off fluorophore switching event, summed over the detection channels and the duration of the event. This value is calculated based on the localization event list after localization filters have been applied.
(3) Mean duration of the switching events, measured in camera exposures. The camera frame rate was 101.8 Hz for all measurements, and the exposure time was 9.3 ms.
(4) Mean background signal level per pixel, per detection channel, per camera exposure. Note that the background level is different in the S- and P-polarized detection channels for the multicolor dataset.

**Table S2:** 4Pi-STORM localization filter parameters

| Filter parameter | Min. value | Max. value | Unit |
| --- | --- | --- | --- |
| Gaussian peak fit width, summed channels <sup>(1)</sup> | 0.8 | 3.0 | pixels |
| Total photons per event, summed channels <sup>(2)</sup> | 1500.0 |  | photons |
| Switching event duration |  | 5 | camera frames |
| S / P photon ratio (multicolor only) | 0.35 | 1.5 |  |

**Notes:**
(1) The four detection channels were transformed to a common coordinate system and summed (see Methods). Each peak was fit with a 2D Gaussian function. This filter is applied to the width (standard deviation) of the Gaussian peak fit results.
(2) Photons per localization event, calculated as the total of all photons detected per on-off fluorophore switching event, summed over the detection channels and the duration of the event.

## Notes

https://www.github.com/gpufit/Gpufit

https://www.github.com/gpufit/Gpuspline

### Competing Interest Statement

The authors have declared no competing interest.

