## Supplementary Video Descriptions for "Optimal Precision and Accuracy in 4Pi-STORM using Dynamic Spline PSF Models"

### Supplemental Video Descriptions

**Supplementary Video S1. The 4Pi PSF numerically phase-shifted over 360 degrees.** The four channels $s_1$ (yellow), $p_1$ (blue), $s_2$ (green), $p_2$ (purple) of the 4Pi PSF were extracted from the scan of a fluorescent bead and rendered as a 3D volume. The phase of the PSF was numerically shifted through several cycles of 360 degrees.

**Supplementary Video S2. Neuronal Cytoskeleton: Detail of Immunolabeled Beta-II Spectrin in a Primary Neuron.** Animation of a relatively linear section of an axon, taken from the dataset shown in Fig. 5. Localizations have been colored according to their *z*-coordinate.

**Supplementary Video S3. Neuronal Cytoskeleton: Immunolabeled Beta-II Spectrin in a Primary Neuron.** Animation of the complete dataset shown in Fig. 5, including a fly-through through the axonal process. Localizations have been colored according to their *z*-coordinate.

**Supplementary Video S4. Neuronal Cytoskeleton: Visualization of the creation of an unwrapped view.** Sequential visualization of perpendicular (*x*-*z*) sections through the 4Pi-STORM image, sliding along the centerline of an axon in which beta-II Spectrin was immunolabeled (as depicted in Supplementary Fig. S24, region 1, first 10 µm). To obtain an unwrapped view, an elliptical band was fit to the Beta-II Spectrin distribution and integrated along the radial direction.

**Supplementary Video S5. Crista Junctions in Mitochondria: Immunolabeled Mic60 in a U-2 OS Cell.** Animation of the dataset shown in Supplementary Fig. S27. Localizations have been colored according to their *z*-coordinate.

**Supplementary Video S6. Crista Junctions in Mitochondria: Immunolabeled Mic60 in a COS-7 Cell.** Animation of the dataset shown in Supplementary Fig. S30. Localizations have been colored according to their *z*-coordinate.

**Supplementary Video S7. Crista Junctions and DNA Nucleoids in Mitochondria: Immunolabeled Mic60 (blue) and dsDNA (yellow) in a COS-7 Cell.** Animation of the multicolor dataset shown in Fig. 6g-j, containing a sequential visualization of thin-slice sections through a single mitochondrion from the 3D dataset.
